# Peripheral positioning of lysosomes supports melanoma aggressiveness

**DOI:** 10.1101/2023.07.07.548108

**Authors:** Katerina Jerabkova-Roda, Marina Peralta, Kuang-Jing Huang, Clara Bourgeat Maudru, Louis Bochler, Antoine Mousson, Ignacio Busnelli, Rabia Karali, Hélène Justiniano, Lucian-Mihai Lisii, Philippe Carl, Vincent Mittelheisser, Nandini Asokan, Annabel Larnicol, Olivier Lefebvre, Hugo Lachuer, Angélique Pichot, Tristan Stemmelen, Anne Molitor, Léa Scheid, Quentin Frenger, Frédéric Gros, Aurélie Hirschler, François Delalande, Emilie Sick, Raphaël Carapito, Christine Carapito, Dan Lipsker, Kristine Schauer, Philippe Rondé, Vincent Hyenne, Jacky G. Goetz

**Affiliations:** Tumor Biomechanics, Strasbourg, France; INSERM UMR_S1109, Strasbourg, France; Université de Strasbourg, Strasbourg, France; Fédération de Médecine Translationnelle de Strasbourg (FMTS), Strasbourg, France; Equipe Labellisée Ligue Contre le Cancer; Institut Curie, PSL, CNRS, UMR144, Paris, France; CNRS UMR7021, Faculté de Pharmacie, Illkirch, France; Institut Gustave Roussy, INSERM UMR1279, Université Paris-Saclay, Villejuif, France; Plateforme GENOMAX, Institut thématique interdisciplinaire (ITI) de Médecine de Précision de Strasbourg Transplantex NG, Fédération Hospitalo-Universitaire OMICARE; 10Service d’Immunologie Biologique, Plateau Technique de Biologie, Pôle de Biologie, Nouvel Hôpital Civil, Hôpitaux Universitaires de Strasbourg, 1 Place de l’Hôpital, 67091, Strasbourg, France; Laboratoire de Spectrométrie de Masse Bio-Organique (LSMBO), IPHC, UMR 7178, CNRS, Université de Strasbourg, Infrastructure Nationale de Protéomique ProFI - FR2048, Strasbourg, France; Faculté de Médecine, Université de Strasbourg et Clinique Dermatologique, Hôpitaux Universitaires de Strasbourg, France; CNRS, SNC5055, Strasbourg, France

## Abstract

Emerging evidences suggest that function and position of organelles are pivotal for tumor cell dissemination. Among them, lysosomes stand out as they integrate metabolic sensing with gene regulation and secretion of proteases. Yet, how their function is linked to their position and how this controls metastasis remains elusive. Here, we analyzed lysosome subcellular distribution in patient-derived melanoma cells and patient biopsies and found that lysosome spreading scales with their aggressiveness. Peripheral lysosomes promote matrix degradation and invasion of melanoma cells which is directly linked to their lysosomal and cell transcriptional programs. When controlling lysosomal positioning using chemo-genetical heterodimerization in patient-derived melanoma cells, we demonstrated that perinuclear clustering impairs lysosomal secretion, matrix degradation and invasion. Impairing lysosomal spreading in two distinct *in vivo* models (mouse and zebrafish) significantly reduces invasive outgrowth. Our study provides a direct demonstration that lysosomal positioning controls cell invasion, illustrating the importance of organelle adaptation in carcinogenesis and suggesting that lysosome positioning could potentially be used for the diagnosis of metastatic melanoma.

## Main

Metastases are responsible for the majority of cancer-related deaths (Dillekås et al., 2019). Melanoma shows strong negative correlation between cancer stage and 5-year patient survival, making it an ideal model to study phenotypic changes leading to cancer cell invasion, adaptation and survival. Melanoma progression consists of multiple sequential events and its early detection is key for patient survival. First, melanocytes are transformed and grow in the epidermis during radial growth phase (RGP), forming a lesion with low potential to develop metastasis. Changes in their transcription program lead to expression of matrix-degrading enzymes and to invasion through the dermis during vertical growth phase (VGP) followed by cancer dissemination through vascular and lymphatic routes, progressing into metastatic stages (Braeuer et al., 2011). To colonize secondary organs during metastasis, melanoma cells sense their microenvironment and react by locally degrading the extracellular matrix (ECM), consisting of structural and specialized proteins (collagen, fibronectin, laminin), which can be remodeled by various proteases, allowing cell dissemination to adjacent tissues (Hua et al., 2011). This suggests that melanoma metastasis requires specific invasion programs for an efficient, targeted delivery and exocytosis of ECM-degrading enzymes. While the implication of lysosome secretion in cell migration has been recently unveiled, how it is orchestrated in invasive cells and whether it might regulate the progression of the disease remains unclear. Distinct reports indicated that lysosomes constitute novel regulators of invasion by allowing cells to sense their microenvironment and trigger adapted responses, notably through the exocytic release of their content (Ballabio and Bonifacino, 2020). For instance, lysosomal exocytosis drives the formation of invasive protrusions resulting in basement membrane breaching in *C. elegans* (Naegeli et al., 2017). In addition, secretion of lysosomal cathepsin B promotes cancer cell invasion and metastasis (Bian et al., 2016). Besides, lysosome secretion contributes to the repair of plasma membrane damages occurring during cell migration and results in better cell survival under mechanical stress (Corrotte and Castro-Gomes, 2019). Importantly, lysosomal activity is regulated by their subcellular location (Johnson et al., 2016; Korolchuk et al., 2011). Peripheral lysosomes are prone to exocytosis and drive growth factor signaling (Jia and Bonifacino, 2019), while perinuclear lysosomes have a decreased pH and higher proteolytic activity (Johnson et al., 2016). Lysosome distribution is in turn directly impacted by endosomal phosphoinositide levels (Mathur et al., 2023) or the cellular microenvironment (Steffan et al., 2009). Lysosomes are transported to the plasma membrane via kinesins (anterograde transport) in response to growth factors and nutrients’ presence, conversely, during starvation and in alkaline environment, lysosomes are transported to the perinuclear region (retrograde transport) in a dynein-dependent manner (Ballabio and Bonifacino, 2020). While the molecular mechanisms driving lysosomal positioning have been broadly studied (Hämälistö and Jäättelä, 2016; Jerabkova-Roda et al., 2024; Pu et al., 2016a), little is known about the *in vivo* functional implications of these changes and whether they can control invasion programs of melanoma cells and their metastatic progression. To answer this question, several studies have described the consequences of manipulating genes involved in lysosome transport on tumor cell invasion (Dykes et al., 2016; Gutierrez-Ruiz et al., 2023; Kundu et al., 2018; Machado et al., 2015; Ping-Hsiu Wu et al., 2020). However, knockdown strategies often impact secondary functions of targeted genes, raising doubts on the precise contribution of lysosome positioning defects in tumor progression. To circumvent these limitations and to fill such gap, we use a unique chemo-genetic tool to control lysosome positioning and to fully demonstrate that lysosomal secretion, regulated via its positioning, controls melanoma invasiveness and impacts its metastatic potential. We reveal a phenotypic switch in the positioning of lysosomes in aggressive melanoma that is supported by distinct transcriptional programs and controls migration and invasion. When impairing the peripheral positioning of lysosomes using a chemo-genetic approach applicable to *in vitro* and *in vivo* models, we reduced the invasion potential of melanoma cells. Importantly, we provide the first evidence in human biopsies that lysosomal positioning strictly correlates with the metastatic progression of the disease. Our study not only illustrates the importance of organelle adaptation in carcinogenesis by providing direct evidences that lysosomal positioning controls secretory pathways of malignant transformation, it also reveals a unique lysosomal phenotype, which could potentially be used for the diagnosis of metastatic melanoma.

### Melanoma invasiveness scales with lysosome spreading

Cells progressing through the metastatic cascade display tremendous phenotypic plasticity in the benefit of increased invasion and ECM degradation potential. In order to investigate invasion-promoting properties, we first characterized a collection of patient-derived melanoma cells from different stages (RGP: WM1552c, WM1862, VGP: WM115, WM983A and metastatic: WM983B, A375 cells). With a collagen invasion assay (Fig S1A), we identified three groups of patient-derived cell lines with low, medium and high invasion potential (Fig S1B), which correlated with their cancer progression state (RGP, VGP and metastatic). Using gelatin degradation assay (Fig S1C), we further showed that cells with high invasion index displayed significantly increased gelatin degradation area and frequency (Fig S1D, E). While gelatin degradation areas were located at invadopodia, specifically identified by the invadopodia markers actin and cortactin (Fig S1C, F), they transiently colocalized with LAMP1, a marker of late endosomes and lysosomes (referred to as lysosomes hereafter) (Fig S1F-H), suggesting that gelatin degradation might involve dynamic shuttling and targeting of lysosomes. These acidic organelles contain metalloproteases (MMPs) and cathepsins that could be released by exocytosis when lysosome transiently co-localize with invadopodia to favor ECM degradation (Jacob and Prekeris, 2015). Indeed, the concerted expression of matrix-degrading enzymes has been previously associated with the transition of melanoma cells from RGP to VGP (Braeuer et al., 2011). To identify the underlying mechanisms, we assessed transcriptional profiles of the three representative patient-derived cell lines of increasing invasion index (WM1862, WM983A, WM983B) using RNAseq and Gene Ontology analysis. This showed an overrepresentation of actin cytoskeleton and cell migration pathways associated with a concomitant reduction of transcripts linked to the lysosomal pathway and metabolism in WM983B cells (Fig 1A and Table 1, 2). We thus hypothesized that increased invasion occurs through a phenotypic switch from a catabolic, lysosomal signature characteristic of RGP cells (WM1862) to a migratory signature found in metastatic cells (WM983B). As lysosome function is linked to its position (Johnson et al., 2016), we investigated their sub-cellular localization in these cell lines by live imaging of freely growing cells and observed that metastatic cells have increased localization of lysosomes to cell periphery as quantified by their 2D radial distribution (Fig 1B, C, Fig S2A, B). We analyzed lysosome positioning of cells plated on micropatterns, which allows for high-throughput study of cells with reproducible shapes (Fig 1D left) and facilitates the comparison and quantitative analysis of lysosome distribution in 2D (Fig 1D right) and in full cell volume (Fig S2C). While RGP cells had mostly perinuclear lysosomes, metastatic cells showed significant dispersion of LAMP1-compartments towards the cell periphery (Fig 1D) characterized by a significant increase in the mean inter-organelle distance and the mean distance to their barycenter (Fig 1E). Moreover, LAMP1-compartments were smaller and more abundant, but globally similarly acidic and degradative in metastatic cells (Fig S2D, E), revealing that the observed transcriptional changes in melanoma cells correlate with changes in LAMP1 distribution. Unbiased analysis of organelles using electron microscopy of the three cell lines revealed strong accumulation of late endosomes/lysosomes in peripheral and protrusive regions of metastatic cells (Fig 1F, G).

**Figure 1:**
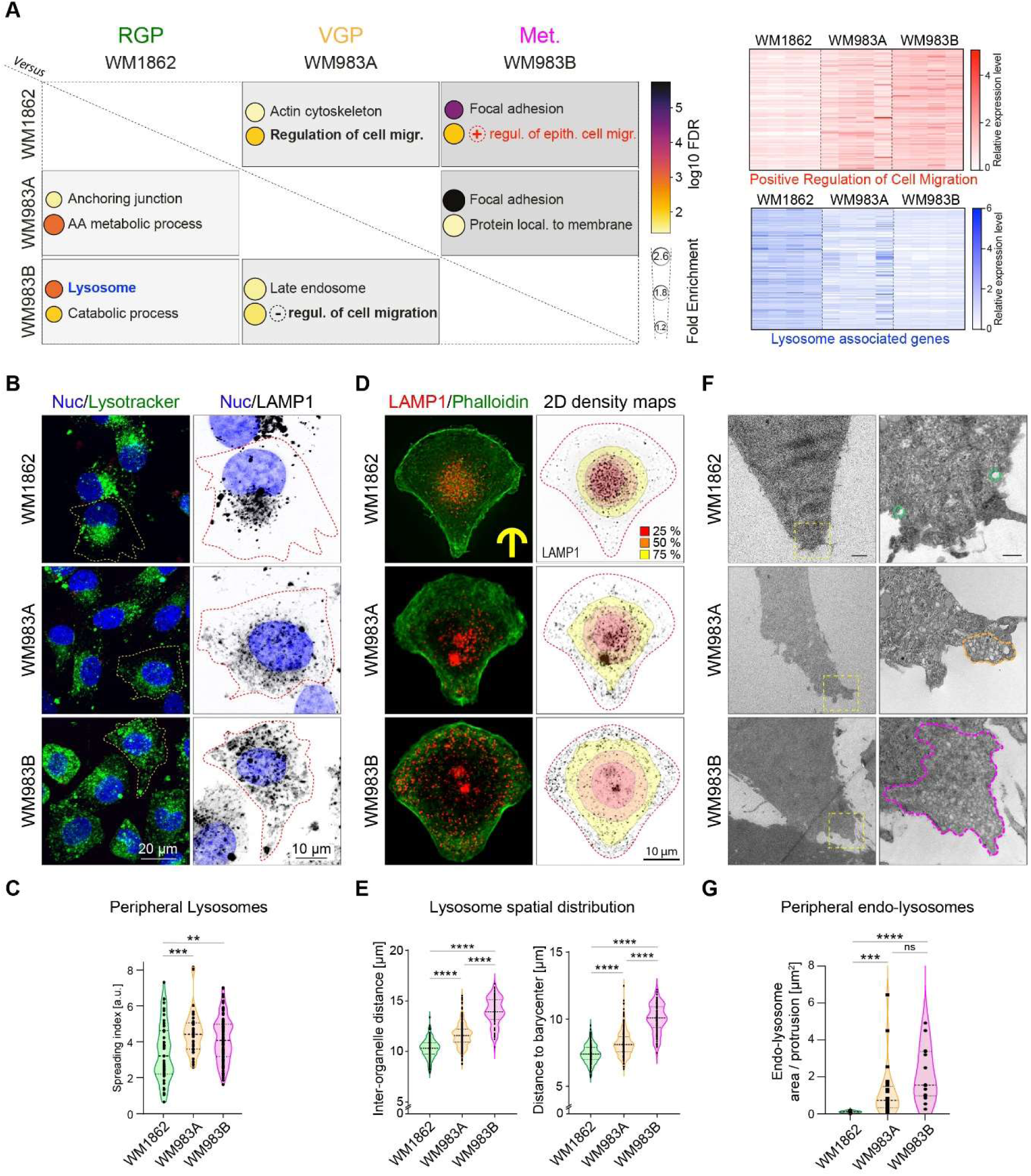
Melanoma invasiveness correlates with lysosome spreading. **A**) Transcriptomics comparison of WM1862, WM983A and WM983B cell lines reveals increased and decreased expression of positive regulation of cell migration and lysosome associated genes, respectively, in metastatic cells. Transcriptomics analysis was performed in quadruplicate, genes showing statistically significant differential expression (pAdj<0.01) were analyzed using gene ontology (GO). Selected GO terms of differentially regulated pathways are listed showing their fold enrichment and log10 FDR in paired comparisons (top panel) and heatmaps are shown for the two main pathways identified (bottom panel), white = low expression, blue/red = high expression. **B-C**) Lysosome positioning in living non-constrained cells. **B**) Representative images of lysosomes stained by NucBlue (nucleus) and green lysotracker (lysosomes) in WM1862 n= 51, WM983A n=40, WM983B n= 108 cells, live cell imaging, in triplicate. **C**) Quantification of lysosome spreading in Cell Profiler. Spreading index (Mean ± SD) = 3.390 ± 1.597, 4.477 ± 1.180, 4.111 ± 1.225, respectively. p values (from top) = 0.0019, 0.0003, Kruskal-Wallis test. One dot represents 1 cell. **D-E**) Lysosome positioning in WM1862, WM983A, WM983B cells grown on 36μm micropatterns. **D**) Representative images of cells stained with LAMP1 antibody and Phalloidin (actin). Top: Immunofluorescence (max z-projection), bottom: overlay of LAMP1 signal and 2D density maps showing LAMP1 distribution (calculated in R software), displaying the smallest area that can be occupied by 25% (red), 50% (orange) and 75% (yellow) of all compartments. **E**) Left: Inter-organelle distance (IOD) represents average of distances between all lysosomes. IOD (Mean ± SD) = 10.30 ± 0.94, 11.56 ± 1.15, 13.99 ± 1.40, respectively. One dot represents 1 cell, p <0.0001 in all conditions. Right: Distance to barycenter (DB) represents the distance from each lysosome to the geometric center (lysosomes). DB (Mean ± SD) = 7.40 ± 0.69, 8.21 ± 0.93, 10.03 ± 1.07, respectively. One dot represents 1 cell, p <0.0001 in all conditions. Kruskal-Wallis with Dunn’s multiple comparison post-hoc test. WM1862, n= 182 cells, WM983A, n= 231 cells, WM983B, n= 82 cells, in triplicate. **F-G**) Electron microscopy reveals that the presence of endolysosomal compartments in protrusions scales with melanoma aggressiveness. **F)** Representative images, **G**) Quantification showing the area occupied by endo-lysosomes in protrusions for each cell line: WM1862 n= 14, WM983A n= 24, WM983B n= 15, Area ± SD = 0.1075 ± 0.05614, 1.219 ± 1.474, 2.106 ± 1.457, respectively. One dot represents 1 field of view, p values (from top) = 0.0001, 0.16, <0.0001, one-way ANOVA. * p<0.05; ** p<0.01; *** p<0.001; **** p<0.0001.

Our results identify and characterize a remarkable cellular phenotype of spread lysosomes associated with aggressive malignancy in melanoma. Since peripheral lysosome positioning has been reported earlier in other cancer types (breast cancer (Ping-Hsiu Wu et al., 2020) and bladder cancer (Mathur et al., 2023)), we next investigated potential molecular mechanisms underlying this phenotype. Transcriptomics data analysis demonstrates that genes known to promote perinuclear localization of lysosomes, such as RILP (Pu et al., 2016b) or RNF167 (Nair et al., 2020), show decreased expression, and conversely, genes linked to anterograde transport, such as KIF1B and KIF5B (Moamer et al., 2019), are overexpressed in metastatic cells (Fig S3A). Besides, other lysosomal genes previously associated with cancer progression such as LAMP1 (Machado et al., 2015; Wang et al., 2017), lysosomal Ca2+ channel MCOLN1 (Medina et al., 2011) and several MMPs (Hua et al., 2011) (MMP2, MMP15, MP17) are also upregulated in metastatic cells (Fig S3A). To probe the molecular mechanism promoting lysosome spreading in melanoma, we focused on the two kinesins, KIF1B and KIF5B, whose expression levels gradually increase with the aggressiveness of patient-derived cell lines (Fig S3B). When any of the two kinesins was downregulated using siRNA (Fig S3C, D), lysosomes relocalized from the periphery to the perinuclear region (Fig S3E, F). In addition, we analyzed sequencing data of primary and/or metastatic melanomas from 331 patients, available at the TCGA database (Akbani et al., 2015). Interestingly, both KIF1B and KIF5B, show an increased expression in samples of metastatic melanomas when compared to primary tumors (Fig S3G), suggesting that they could, in part, be responsible for a correlation between lysosomal positioning and melanoma progression.

### In patients with metastatic melanoma, lysosomes are relocated to the cell periphery

Following our observations on patient-derived cell lines, we tested whether changes in lysosome positioning can also be observed in biopsies from melanoma patients and whether specific patterns correlate with the progression stage. For that purpose, we analyzed lysosome subcellular distribution in samples from human biopsies of healthy skin, benign naevi, primary melanoma and melanoma skin metastases (Fig 2A). Strikingly, while Sox10-positive melanocytes of healthy and benign tissues displayed strong lysosome perinuclear clustering, cells of melanoma cutaneous metastases showed striking peripheral positioning of lysosomes (Fig 1B, C). In the primary melanoma, lysosome spreading was variable but rather low when compared to metastatic melanoma, suggesting that peripheral lysosomes are only appearing in cells that are bound to be metastatic, and could potentially serve as a prognostic marker. In addition, while the naevi samples displayed heterogeneous positioning of lysosomes, the proportion of cells with peripheral lysosomes increased with the depth away from the epidermis (Fig 2D, E), suggesting that more complex programs might be at play to stimulate the changes in lysosome positioning and to promote tumorigenesis. Whether this is also true for primary melanoma remains to be investigated as the lesions were too small to perform spatial analysis. Taken together, these results show that lysosome spreading correlates with melanoma stage, in particular, metastatic melanoma cells redistribute their lysosomes towards cell periphery, as seen in collection of tumor biopsies from a cohort of patients. Our results connect lysosome spreading to melanoma metastasis and support the importance of this phenotype for future clinical applications as for example, improving the classification criteria for malignant and non-malignant lesions and thus allowing for early detection of lesions with metastatic potential and timely intervention.

**Figure 2:**
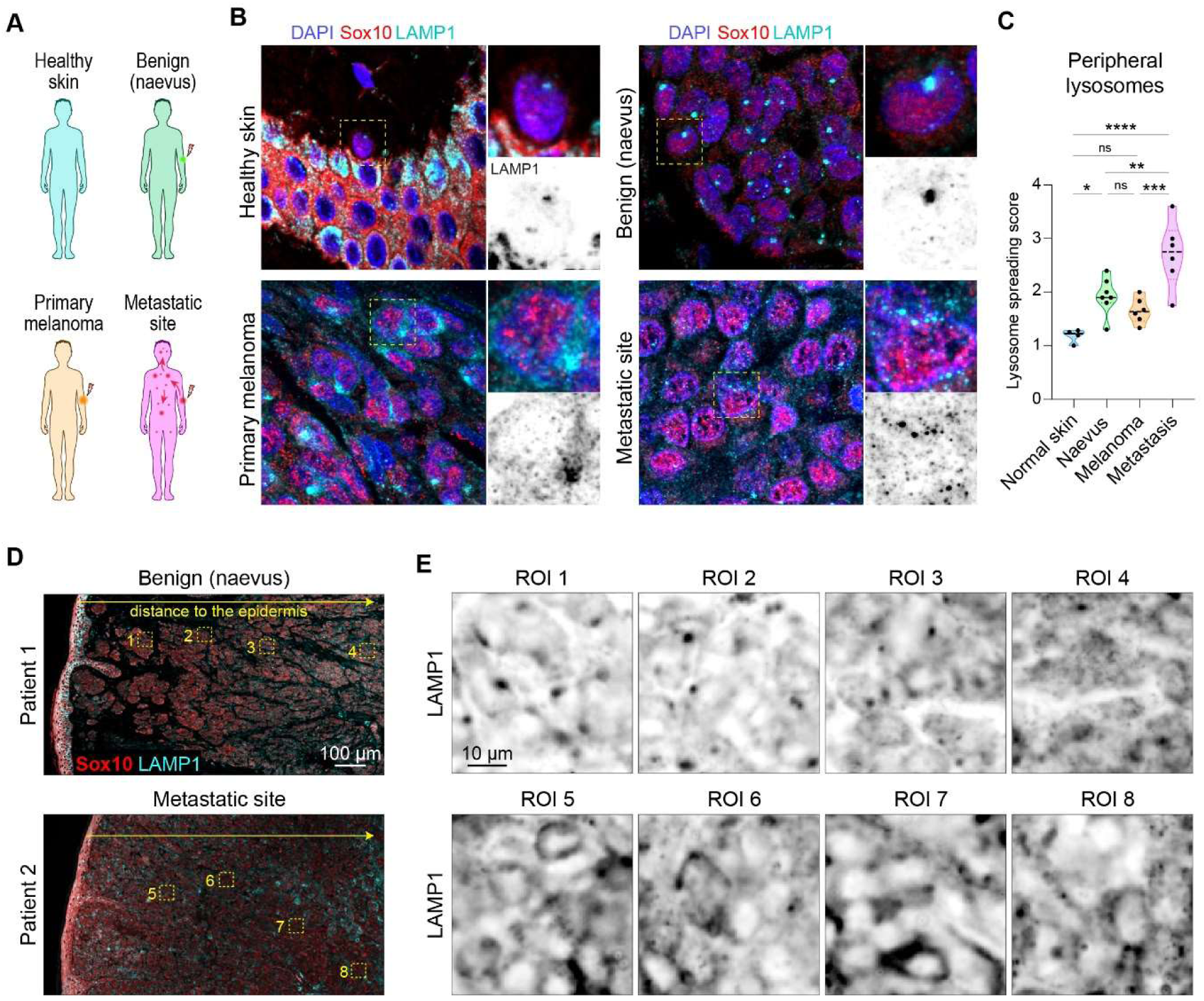
Lysosomes spread as melanoma progresses towards metastasis in human biopsies. **A)** Samples of patient biopsies were obtained from healthy skin donors and from patients with benign tissue (naevus), primary melanoma and metastatic melanoma (skin metastasis). **B)** Representative images of patient biopsies sections: samples were labelled for SOX10 (red), LAMP1 (cyan) and nuclei (blue) by immunofluorescence, full tissue section was imaged by slide scanner. **C**) Quantification of lysosome spreading. Ten random regions were analyzed in a blinded setup for lysosome positioning. Lysosome spreading, number of patients: healthy skin n= 4, naevus n=7, primary melanoma n=6 and metastatic melanoma n=6. Spreading score (Mean ± SD) = 1.184 ± 0.1276, 1.929 ± 0.3450, 1.656 ± 0.2373, 2.793 ± 0.6094, respectively. One dot represents 1 patient, p values (from top, left) = <0.0001, 0.1543, 0.0087, 0.0218, 0.2299, 0.001, one-way ANOVA, * p<0.05; ** p<0.01; *** p<0.001; **** p<0.0001. **D-E**) Lysosome spreading shows a spatial heterogeneity within the tumor mass. **D**) Representative images from Patient 1 = benign tissue (Naevus) and Patient 2 = metastatic melanoma showing low magnification image mapping the tissue section, immunohistochemistry: LAMP1 (cyan), SOX10 (red). **E**) Zoomed regions from panel D, showing the progressive change in lysosome spreading with increasing distance to the epidermis, single channel: LAMP1 antibody staining.

### Forcing lysosome perinuclear clustering in metastatic melanoma cells

To study the consequences of lysosome repositioning for metastatic evolution of melanoma, we engineered melanoma cells employing a chemo-genetic strategy based on the fast and strong heterodimerization of the FKBP and FRB domains by Rapalog (Kapitein et al., 2010). We chose WM983A and WM983B cell lines, derived from the primary tumor and metastatic site of the same patient, respectively. We stably expressed FKBP domain fused to LAMP1 and the FRB domain fused to dynein adaptor BicD2 (Fig 3A). Rapalog treatment induced binding of BicD2 and recruitment of Dynein to LAMP1, forcing lysosome movement towards the minus end of microtubules, and thus perinuclear clustering of the LAMP1 compartment around the microtubule organizing center (Fig 3B, S4F). Clustering was fast, dose-dependent (Fig S4A, B) and persistent in time (Fig S4C, D, E), as previously described (Kapitein et al., 2010), allowing precise control of lysosomal positioning *in vitro* and *in vivo.* Engineered control cells expressing only single domain FKBP (FKBP only) did not show lysosome clustering upon Rapalog treatment (Fig S4A, C, D). Correlative light and electron microscopy (CLEM) of control- or Rapalog-treated WM983B-LAMP1-mCherry cells showed colocalization between LAMP1-mCherry and vesicular compartments, which clustered as expected in the perinuclear region upon Rapalog treatment (Fig 3C). Notably, Rapalog does not impair the colocalization between LAMP1-mCherry and the BODIPY-Pepstatin A (Fig S4G), which is delivered to lysosomes via endocytic pathway (Chen et al., 2000) and binds to the active site of cathepsin D in acidic conditions, suggesting that clustering by Rapalog did not disrupt cargo delivery and lysosomal catabolic activity in melanoma cells. Of note, Rapalog is a chemically modified analog of Rapamycin unable to bind lysosome-related mTor and to thus perturb mTORC1 signaling (Bayle et al., 2006). Accordingly, we could not detect any changes in mTORC1 signaling in our model under normal conditions of culture (Fig S4H-I). As such, this tool brings great advantage over genetic studies targeting the known regulators of lysosome positioning as it allows to reposition specifically the LAMP1 positive organelles without impacting secondary functions of selected genes, caused by protein depletion. Proteins controlling lysosome positioning often have other functions: KIF1B and KIF5B for instance are known to regulate the localization of other organelles, endosomes and mitochondria (Hooikaas et al., 2019; Serra-Marques et al., 2020) while RILP controls lysosomal pH (Mulligan et al., 2024), and RNF167 regulates ubiquitination (Nair et al., 2020). By contrast, Rapalog can be used to study the direct link between lysosome position and cancer invasiveness.

**Figure 3:**
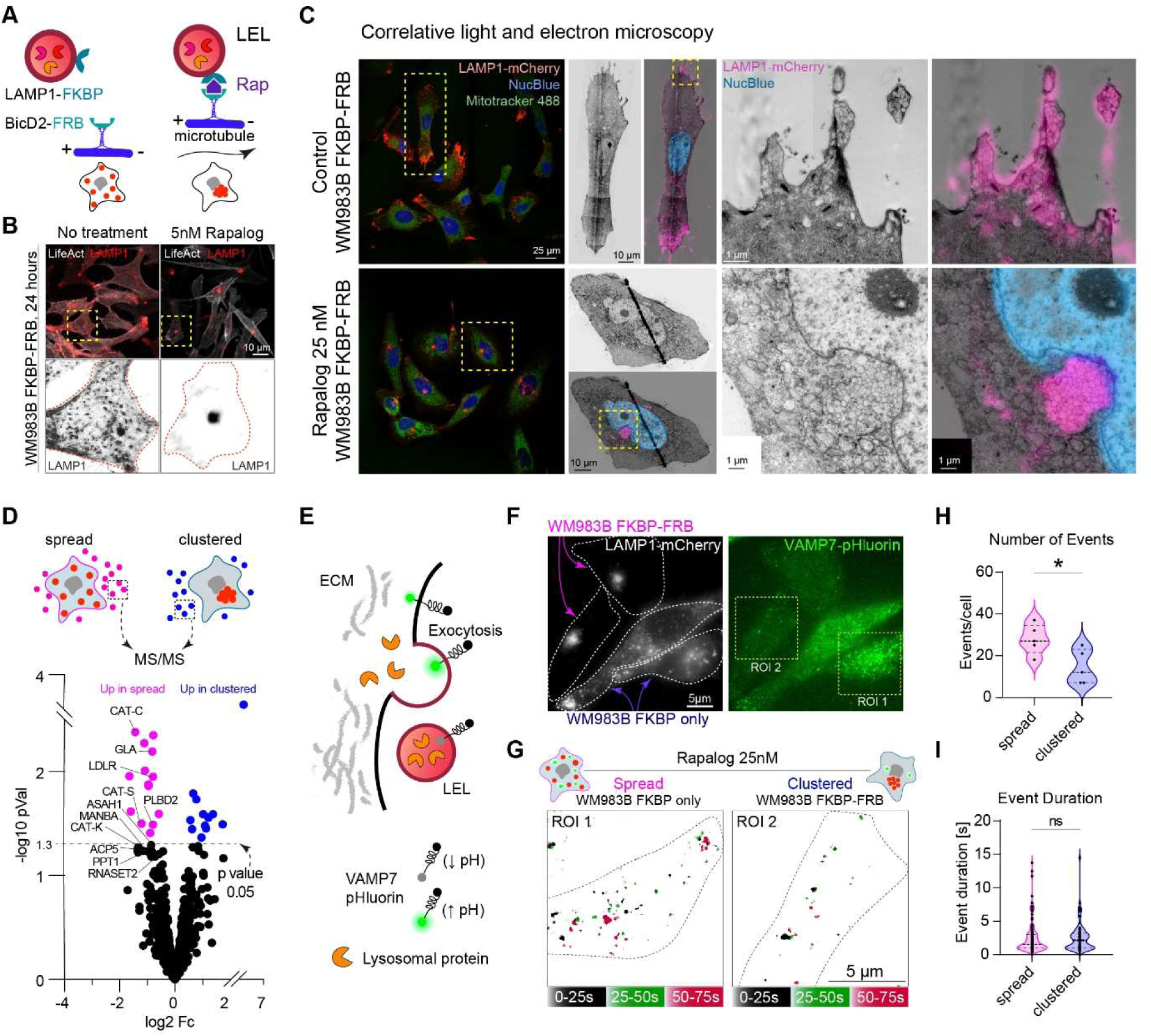
Lysosome perinuclear clustering inhibits lysosome exocytosis, matrix degradation and cell invasion. **A-B**) Rapalog mediated-perinuclear clustering of lysosomes. **A**) Lysosomes in WM983A or WM983B cells stably expressing LAMP1-mCherry-FKBP and BicD2-FRB can be clustered using the compound Rapalog (Rap) which induces FKBP-FRB rapid heterodimerization. **B**) Representative images showing stable lysosome perinuclear clustering 24h after 5nM Rapalog treatment. **C**) Correlative light and electron microscopy reveals clustered perinuclear LAMP1 positive compartments. Representative immunofluorescent images and a high magnification overlay of fluorescent and electron microscopy images (LAMP1 appears in pink, nucleus in blue) are shown for each condition. **D**) Melanoma secretome is altered by lysosome perinuclear clustering. Differential quantitative mass spectrometry analysis of proteins secreted to the cell medium by WM983B (FKBP-FRB) cells in the absence or presence of 5nM Rapalog. Proteins known to be lysosome-associated are labelled with their name. Magenta = proteins upregulated in cells with spread lysosomes, blue = proteins upregulated in cells with clustered lysosomes (p<0.05). **E-I**) Rapalog-induced perinuclear clustering inhibits lysosome exocytosis. **E**) Lysosome secretion was assessed by TIRF microscopy using VAMP7-pHluorin probe, which is quenched in lysosome acidic environment and brightly fluorescent once exposed to the alkaline pH of the extracellular space. **F**) Representative images of co-cultured WM983B-LAMP1-mCherry cells expressing either single heterodimerizing domain (FKBP only) or both the domains (FKBP-FRB) treated with 25nM Rapalog, TIRF microscopy. **G**) TIRF movie was divided into three time-segments (black: 0-25 seconds, blue: 25-50 seconds, magenta: 50-75 seconds) and displayed as maximum projection showing the number of events per each time-segment. Two representative examples: ROI 1 = spread lysosomes, ROI 2 = clustered lysosomes. **H-I**) Graphs showing number of lysosome secretion events (**H**, one dot represents one cell) and event duration (**I**, one dot represents one event) (five cells each). Events per cell (Mean ± SD): spread = 27.80 ± 7.19, clustered = 14.40 ± 8.23, p value = 0.0397, Mann-Whitney test. Event duration (Mean ± SD): spread = 2.46 ± 2.36, clustered = 2.56 ± 2.20 seconds, p value = 0.3518, Mann-Whitney test. **J-L**) Lysosome perinuclear clustering inhibits gelatin degradation.

### Peripheral lysosomes promote secretion and matrix degradation

Lysosome exocytosis of different proteases and subsequent extracellular matrix degradation promotes invasion to adjacent tissues (Monteiro et al., 2013; Naegeli et al., 2017). We thus first investigated how altering positioning of the LAMP1 compartment impacts the cell secretome. We analyzed the concentrated cell supernatant of WM983B cells in the presence and absence of Rapalog treatment by mass spectrometry (Fig 3D, Table 3) and by human protease array (Fig S5A, Table 4). Together, these results show an overall decrease in the total protease content secreted by the cells when lysosomes are clustered close to the nucleus (Fig S5A, Table 3, 4). We identified a significant decrease of several lysosome-associated proteins upon lysosome perinuclear clustering, including cathepsins and various metalloproteases from the ADAM, Kallikrein and MMP families (Fig 3D, Table 4). These enzymes contribute to ECM degradation (Vidak et al., 2019), tumor growth and invasion (Gocheva et al., 2006) and they have been linked to metastatic progression, for instance in the case of cathepsin S or B in gastric and colorectal cancers respectively (Bian et al., 2016; da Costa et al., 2020). To confirm that Rapalog-induced perinuclear clustering inhibits lysosome exocytosis, we imaged VAMP7-pHluorin, a v-SNARE involved in the fusion of lysosome with plasma membrane (Chaineau et al., 2009; Lachuer et al., 2023) using TIRF microscopy (Fig 3E, F). Quantification reveals that cells with perinuclear clustering showed significantly reduced numbers of VAMP7 secretory events (Fig 3G, H, Fig S5B), with no impact on the duration of the secretion process (Fig 3I). These experiments demonstrate that perinuclear clustering of lysosomes impairs lysosome exocytosis by metastatic melanoma cells. These results are in line with a previous observation of peripheral lysosomes promoting their fusion with plasma membrane and thus exocytosis in a model of lysosomal storage disease (Medina et al., 2011). Next, we investigated whether perinuclear lysosome clustering would impair the ECM degradation machinery (Fig 4). Gelatin degradation area was reduced in both the primary tumor (WM983A) and the metastatic cells (WM983B) upon perinuclear lysosome clustering (Fig 4A, B, Fig S5C, D). While degradation frequency in WM983A (VGP) cells remained unaltered (Fig S5E), the metastatic WM983B cells showed a significant decrease upon lysosome peri-nuclear clustering (Fig 4C). We then tested the impact of forced peri-nuclear clustering of lysosomes on invasion of melanoma cells (Fig 4D). Untreated cells (FKBP-FRB) and Rapalog-treated control cells (FKBP only) have peripheral lysosomes, which were perinuclear in Rapalog-treated cells expressing both FKBP-FRB domains (Fig 4D bottom). Rapalog treatment alone had no effect on the invasion index, but Rapalog-induced perinuclear lysosome clustering significantly decreased the invasion potential of the metastatic melanoma cells (Fig 4E). These results further confirmed that peripheral lysosome positioning promotes lysosome exocytosis which contributes to ECM degradation and cell invasion. Forcing perinuclear lysosome localization rescues these phenotypes, suggesting that lysosome position and subsequent secretion promotes ECM remodeling, as often seen in aggressive cancers (Winkler et al., 2020). More globally, cancer cell secretion is likely to further shape pro-metastatic features of the tumor microenvironment, and to favor the emergence of, for example, cancer associated fibroblasts, whose ECM remodeling expertise is pivotal during tumor progression (Kalluri, 2016; Sahai et al., 2020). Of note, we have observed transient localization of lysosomes to invadopodia, in line with earlier observation that targeted secretion of CD63-positive multi-vesicular bodies promotes invadopodia formation (Hoshino et al., 2013). Yet, while LAMP1 is highly enriched in protrusive regions of melanoma cells, its co-localization with invadopodia markers was only partial and transient, suggesting that lysosomal exocytosis likely occurs outside of active invadopodia as well. It has been reported that microtubule-transported post-Golgi carriers are secreted near focal adhesions, which capture and stabilize microtubules (Fourriere et al., 2019). Lysosomal exocytosis visualized by VAMP7-pHluorin can occur at focal adhesions (Lachuer et al., 2023) that have been associated with invasive matrix degradation (Wang and McNiven, 2012). Alternatively, CD63+ late endosomes/lysosomes are re-localized to the cell periphery and protrusive structures, where they could, for example, promote the secretion of pro-tumorigenic extracellular vesicles (Hoshino et al., 2013). Whether distinct types of late endosomes act in concert to favor ECM degradation, or whether it involves hybrid late endo-lysosome compartments remains to be determined.

**Figure 4:**
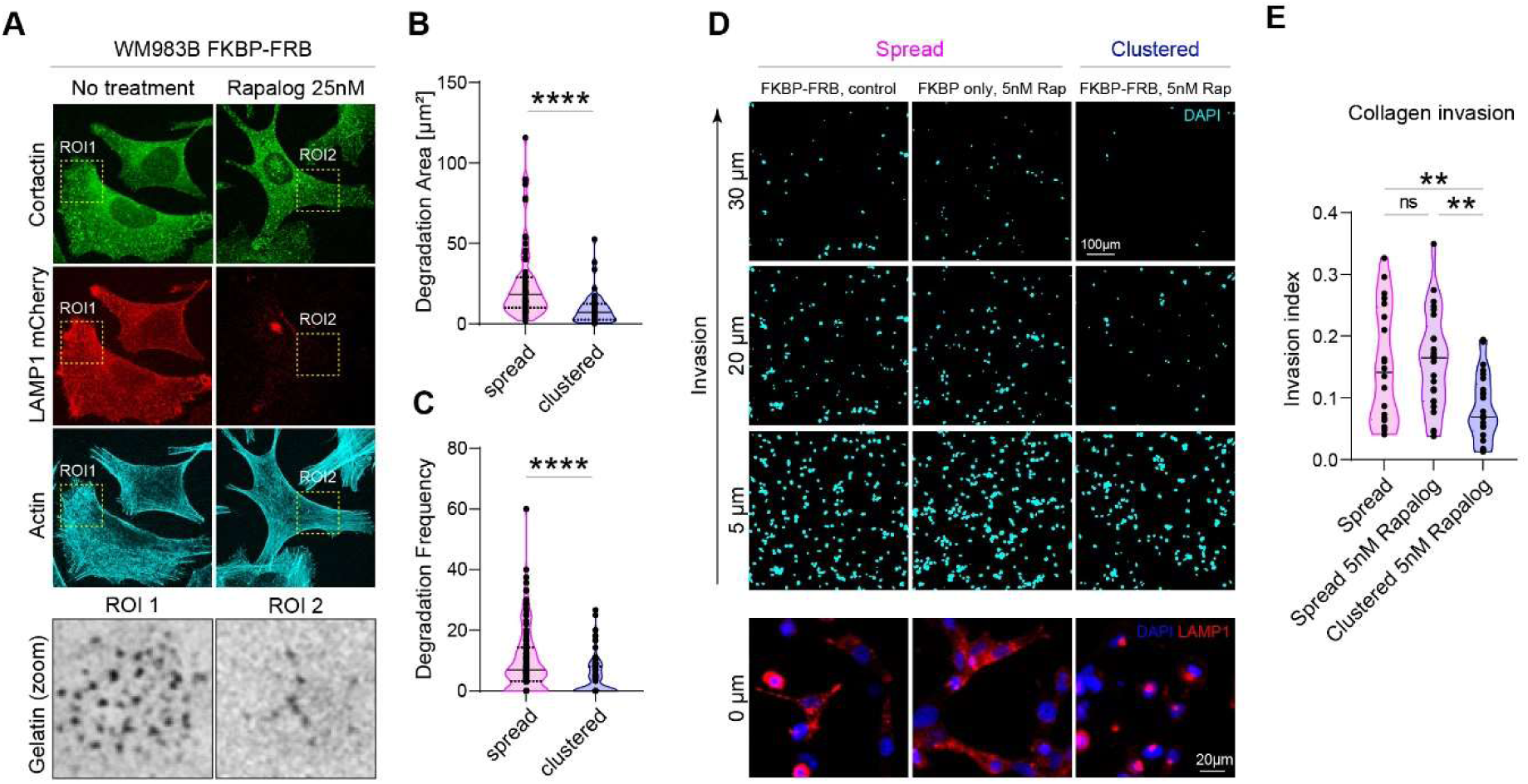
Lysosome clustering impairs cell secretion, matrix degradation and cell invasion. **A-C)** Peripherally positioned lysosomes promote gelatin degradation. **A**) Representative images of WM983B (LAMP1-mCherry-FKBP-FRB), cultured on FITC-gelatin in presence or absence of 25nM Rapalo and stained with cortactin/phalloidin, TIRF microscopy. **B**) Degradation Area (Mean ± SD) = 24.75 ± 23.29 n= 73 cells, 9.41 ± 9.94 n= 50 cells, respectively, p <0.0001 Mann-Whitney. **C)** Degradation frequency (Mean ± SD) = 10.14 ± 10.45 n= 138 cells, 4.43 ± 6.04 n= 102 cells, respectively, p <0.0001 Mann-Whitney. One dot represents 1 cell. **D-E**) Lysosome perinuclear clustering decreases collagen invasion. **D**) WM983B cells with spread (FKBP-FRB control and FKBP only with Rapalog 5nM) and clustered lysosomes (FKBP-FRB with Rapalog 5nM), showing representative images of lysosome positioning in the three groups (bottom) and representative images of cell invasion after 24h (top). **E**) Quantification of the invasion index: FKBP-FRB n= 20, FKBP only n= 25, FKBP-FRB (clustered) n= 25, (Mean ± SD) = 0.1526 ± 0.094, 0.1592 ±0.079, 0.0836 ± 0.053, respectively, in triplicate, p value (from top, left) = 0.0093, 0.9548, 0.0021, one-way ANOVA. One dot represents 1 field of view. * p<0.05; ** p<0.01; *** p<0.001; **** p<0.0001.

### Forcing lysosomal clustering impairs invasion potential of melanoma cells

Building on the stability of Rapalog-mediated lysosomal clustering in patient-derived cells and the demonstration that it impairs ECM degradation and cell invasion *in vitro*, we sought to monitor if lysosome position impacted melanoma dissemination *in vivo*. We first set-up an experimental approach based on subcutaneous grafting of fluorescently-labeled (Cell Trace) melanoma cells in ears of nude mice (Fig 5A). Tumors made of control cells (FKBP only) and cells with both FKBP-FRB domains were subjected to topic treatment with Rapalog (5nM) every 2 days. Rapalog treatment alone had no effect on cell proliferation or cell viability *in vitro* (Fig S5F, G). Upon 11 days of growth and dissemination, we serially sectioned the ears and quantified the range of invasion. Tumors (n=2) made of melanoma cells with peripheral lysosomes displayed increased levels of dissemination (Fig 5B, C) as quantified by the area of fluorescently-labeled cells in each section (Fig S6A). In contrast, melanoma tumors (n=2) with perinuclear lysosome clustering displayed limited dissemination potential (Fig 5B, C) demonstrating that forced perinuclear clustering of lysosomes impairs the metastatic properties of melanoma tumors *in vivo*. To further demonstrate that melanoma invasiveness is altered by lysosome perinuclear clustering, we switched to an alternative experimental metastasis model where invasion can be tracked in real time. To do so, we injected melanoma cells with different lysosome clustering status intravenously in two days post-fertilization (dpf) zebrafish embryos (Follain et al., 2018) to probe lysosomal clustering while assessing the metastatic and invasive potential of melanoma cells over time (Fig 5D, Fig S6B). Lysosome clustering was stable *in vivo* and visible in round circulating tumor cells that had just arrested in the vasculature after injection (Fig 5E) as well as in the growing non-invasive tumor cells three days post injection (Fig 5F, bottom). At day 3, while all melanoma cells had efficiently extravasated (Fig 5G), independently of their lysosomal positioning, cells with peripheral lysosomes displayed increased post-extravasation invasion potential *in vivo* (Fig 5H, Fig S6C, D). Notably, Rapalog treatment alone had no impact on the invasion index (Fig 5H) when we compared untreated cells (FKBP-FRB) and Rapalog-treated control cells (FRB only) which both have spread lysosomes (Fig 5E, F). On the contrary, when lysosomes were clustered in melanoma cells before injection, their ability to invade after extravasation was strongly impaired (Fig 3F, H). Collectively, these data show that lysosome positioning is an important driver of melanoma invasiveness, both at the level of the primary tumor (Fig 5A-C) and in metastatic dissemination (Fig 5D-H). As seen in our study, cells with peripheral lysosomes have higher invasion potential *in vitro* as well as *in vivo*, which can be rescued by promoting the perinuclear lysosome clustering and thus reducing their malignancy, providing the first *in vivo* molecular control of lysosome positioning in metastatic progression.

**Figure 5:**
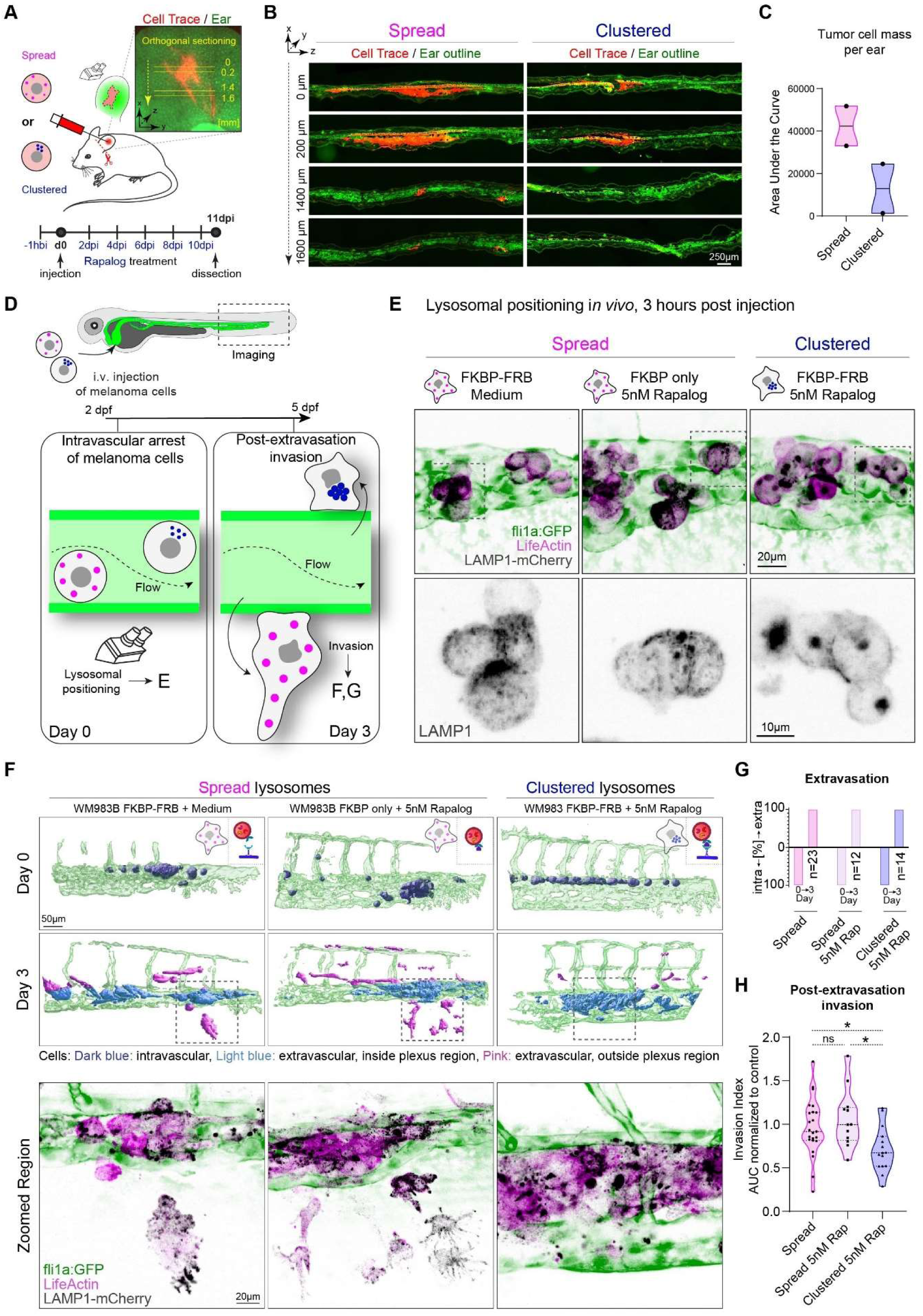
Lysosome clustering impairs cancer cell invasion *in vivo*. **A-C)** Lysosome clustering prevents tumor progression in mice. Cells with spread (FKBP only, 5nM Rap) and clustered (FKBP-FRB, 5nM Rap) lysosomes were injected intradermally in mouse ear. **B)** Representative Immunofluorescence examples of mouse ear sections. Red = tumor cells, green = ear (autofluorescence). **C**) Quantification of cell mass in the ears when lysosomes are spread (n=2) and clustered (n=2), calculated from cell area per each section and distance in tumor: (Mean ± SD) = 42 380 ± 13 200, 12 901 ± 16 456, respectively. **D-H**) Perinuclear lysosome clustering prevents cell invasion in zebrafish. **D**) Illustration of the invasion assay in zebrafish. Fli1a:GFP (green endothelium) zebrafish embryos (48 hpf) were injected intra-vascularly with WM983B cells and imaged at day 0 and day 3 post injection to assess the post-extravasation invasion potential of cancer cells with different lysosomal positioning. **E**) Representative confocal images of WM983B cells (stably expressing LAMP1-mCherry= black, and LifeActin-miRFP670= magenta) with spread (FKBP-FRB control; FKBP only, 5nM Rapalog) and clustered lysosomes (FKBP-FRB, 5nM Rapalog) at 3 hours post-injection shown as maximum z projection. **F**) IMARIS segmentation (*top*), representative examples, day 0 (3 hours after cell injection) and at day 3 (72 hours post injection). (bottom) Zoomed regions showing confocal images corresponding to the IMARIS segmentation. images are shown as a maximum z projection (LAMP1-mCherry= black, LifeActin-miRFP670= magenta). **G**) Graph showing the cell extravasation at day 0 and day 3. Negative axis= intravascular cells, positive axis= extravascular cells. **H**) Post-extravasation invasion (PEI) potential (at 3dpi) was calculated as a ratio: area of cells that migrated outside of the vasculature region to the total area of cells. FKBP-FRB (spread) n= 23 embryos, FKBP only (spread) n= 12 embryos, FKBP-FRB (clustered) n= 15 embryos, PEI (Mean ± SD) = 0.9616 ± 0.3284, 1.060 ± 0.3316, 0.7085 ± 0.2568, respectively, in triplicate, p values (from top, left) = 0.0317, >0.9999, 0.0172, Kruskal-Wallis test. One dot represents 1 fish, normalized to spread condition. * p<0.05; ** p<0.01; *** p<0.001; **** p<0.0001.

Organelles are dynamic, self-organized structures whose specific function is inevitably linked to their position and morphology (Schauer et al., 2010), in space and time within cells (Ballabio and Bonifacino, 2020; van Bergeijk et al., 2016). In this study, we provide first direct demonstration that the position of lysosomes within cells tightly controls the targeted secretion of matrix-degrading enzymes which subsequently promotes melanoma cell invasion and metastatic progression. While our work provide a direct link with melanoma metastasis, a recent study shows that relocalization of lysosomes to the cell periphery promotes the emergence of leader cells in collective epithelial cell migration (Marwaha et al., 2023). Re-positioning at the periphery enables localized secretion of proteolytic enzymes contained in lysosomes at focal adhesion (Lachuer et al., 2023), invadopodia (Nakahara et al., 1997) or within broader protrusive regions of the cell. While more work is required to understand the switches that relocalize lysosomes, our study demonstrates that peripheral lysosome positioning promotes cell invasion through lysosomal secretion and ECM remodeling and thus contributes to metastatic progression. In addition, although we did not detect any change in lysosomal degradative capacities upon lysosome peri-nuclear clustering, it could potentially alter tumor progression by mediating nutrient sensing (Jia and Bonifacino, 2019; Korolchuk et al., 2011) or chemoresistance through the secretion of chemotherapeutics stored in lysosomes (Machado et al., 2015). Our study exploited a chemo-genetic model of forced lysosome clustering which requires cell engineering and would therefore benefit from the complementary use of small molecules that regulate lysosome positioning, particularly for clinical applications. We previously identified PI3K inhibitors as potent lysosome clustering agents in bladder cancer (Mathur et al., 2023). Indeed, lysosome positioning, and more general organelle topology (Wang et al., 2023), were used as a readout in a screen for novel therapeutic drugs and targets (Circu et al., 2016). This opens an exciting area of research leading to a wider drug discovery approach centered on organelle positioning.

Probing of spatial distribution and morphology of lysosomes in human tumors could constitute a novel indicator of tumor progression, as it is the case for cell shape or additional morphometric analysis (Sero et al., 2015; Pei-Hsun Wu et al., 2020). We now demonstrate that lysosomal peripheral distribution can be used as a proxy for melanoma metastatic dissemination. We further show that deeply invading cells away from the epidermis tend to display increased lysosome spreading as if such repositioning increased with the invasive properties of cells. This suggests that probing lysosomal positioning could allow to refine classification criteria for malignant and non-malignant cutaneous lesions. Several other organelles, such as mitochondria, are intimately linked to cancer progression and probing simultaneously multiple organelles, as it can be done using whole-cell segmentation of high-resolution images (Heinrich et al., 2021), could document precisely which organelles, and their contacts, are repositioned during melanoma progression. Interestingly, deep learning-based approaches recently demonstrated the power of correlating breast cancer status with organelle topology, which out-performed morphology-based features, further highlighting the need to consider organelles positioning, and their interactome, as a new cancer rheostat that could be exploited for better diagnosis (Wang et al., 2023).

## Acknowledgements

We thank all members of JGG’s and KS’ teams for discussions on this topic. JGG is the coordinator of the NANOTUMOR Consortium, a program from ITMO Cancer of AVIESAN (Alliance Nationale pour les Sciences de la Vie et de la Santé, National Alliance for Life Sciences & Health) within the framework of the Cancer Plan (France) that has mostly supported this work, including the teams of KS and PR. Work and people in the lab of JGG are also supported by the INCa (Institut National Du Cancer, French National Cancer Institute), charities (La Ligue contre le Cancer, which also supports PR, and ARC (Association pour la Recherche contre le Cancer), FRM (Fondation pour la Recherche Médicale)), the National Plan Cancer initiative, the Region Est, INSERM and the University of Strasbourg. Proteomics experiments were supported by the French Proteomic Infrastructure (ProFI FR2048, ANR-10-INBS-08-03). KJR is supported by a post-doctoral fellowship SPF202004011876 from FRM and the NANOTUMOR consortium. HL was supported by ARC and FRM. KJH is supported by the NANOTUMOR consortium and Taiwan National Science and Technology Council (NSTC-112-2917-I-564-038). We are also thankful to recent donators (Rohan Athlétisme Saverne) to support our work. The imaging was supported by CRBS imaging platform PIC-STRA with assistance from P. Kessler, by the Imaging Center of IGBMC with assistance from E. Grandgirard and E. Guiot, by the PIQ Platform with assistance from R. Vauchelles and by the Cell and Tissue Imaging Platform (PICT-IBiSA), member of the national infrastructure France-BioImaging supported by the French National Research Agency (ANR-10-INBS-04). Electron microscopy was performed with help from C. Royer and V. Demais (INCI, Strasbourg, France). We would like to thank C. Renaud and C. Brubach for technical assistance during the preparation of patient biopsies. We thank T. Galli (IPNP, INSERM U1266, Paris) for sharing the VAMP7-pHLuorin constructs, A-C. Reymann and R. Benoit for sharing their resources, D. Sampaio Goncalves for help with ImageJ macro writing, Mei Li and A. Perrin for help with mouse intradermal injections, M. Durik for valuable feedback on the work and F. Colin for help with graphical representation of the data. Parts of the figures were drawn using pictures from Servier Medical Art which is licensed under a Creative Commons Attribution 3.0 Unported License (creativecommons.org/licenses/by/3.0/). We are grateful to F. Saltel and B. Bonnard (BRIC, Inserm U1312, Bordeaux) for their contribution on invadopodia analysis.

## Contributions

KJR, VH, KS, PR and JGG conceived project and designed the experiments. KJR performed most of the experiments and analysis. AM, RK, HJ, LML, PC, ES and PR performed collagen gel invasion assays and invadopodia experiments. KJR, NA and MP performed zebrafish experiments. KJR and IB performed CLEM experiments. KJH performed EM analysis. LB, OL, VH and KJR performed mice experiments. CBM was involved in organelle positioning imaging and analysis. VM analyzed TCGA data. QF and FG designed and performed cytometry analysis. AL and OL performed molecular biology. KJR performed and analyzed micropatterning experiments using tools developed by HL. AP, TS, AM, OL, VH and RC performed and analyzed RNAseq experiments. AH, FD and CC performed and analyzed mass spectrometry experiments. DL and LS provided human biopsies. VH, KS, PR and JGG provided funding. VH and JGG supervised the study. KS and PR co-supervised the study. KJR, VH and JGG wrote the manuscript, with input from KS and PR. All authors proofread the manuscript.

---

**Figure S1:**
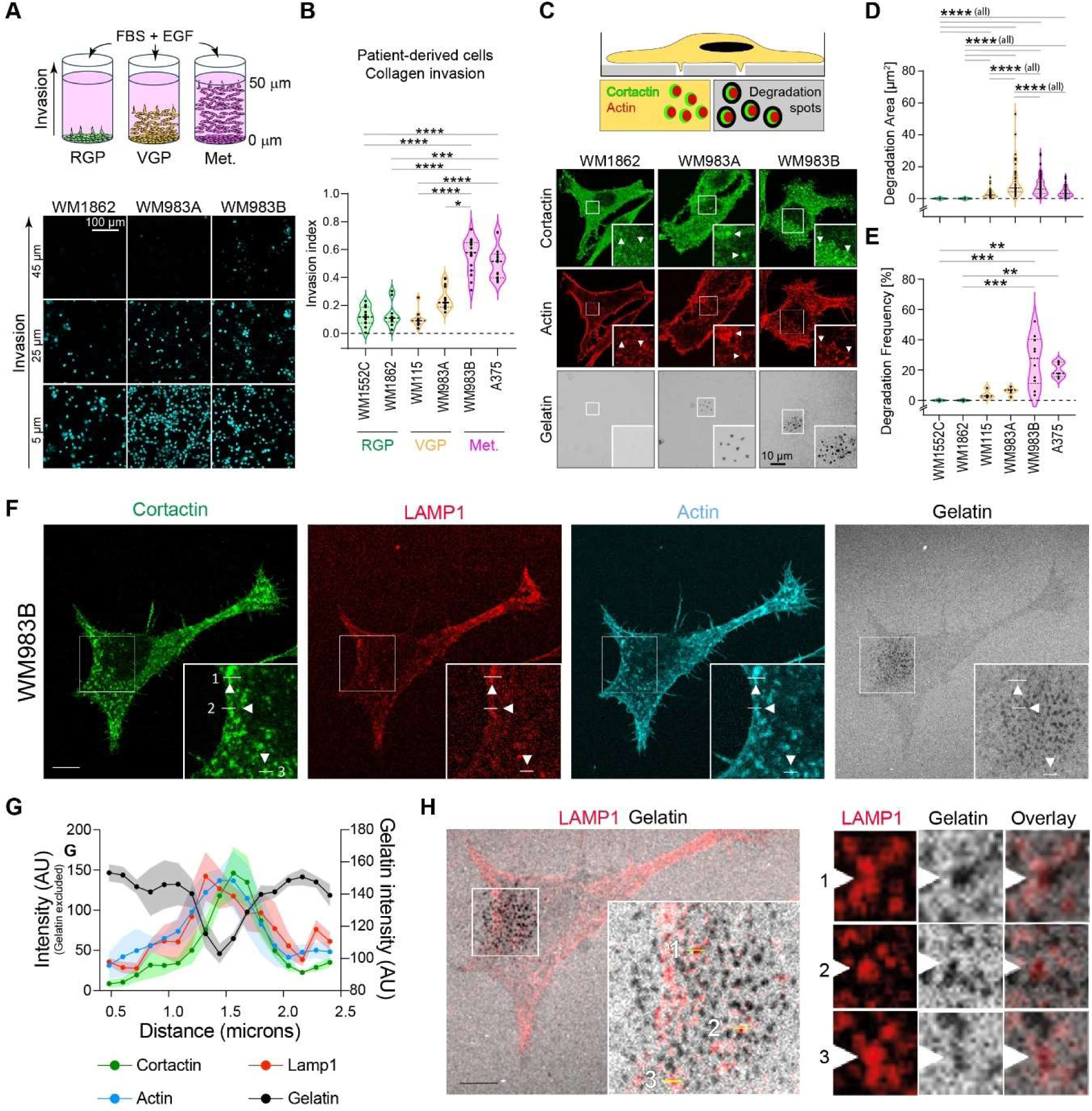
Melanoma invasiveness correlates with matrix degradation. **A-B**) Invasive potential of human melanoma cells (WM1552c, WM1862, WM115, WM983A, WM983B, A375) was analyzed using collagen invasion assay. **A**) Representative images showing cell invasion after 24 hours. Spreading distance was analyzed by confocal imaging. **B**) For each cell line, the invasion index equals the number of nuclei above 10 µm distance divided by the total number of cells, (Mean ± SD) = 0.1213 ± 0.070, 0.1334 ± 0.089, 0.2500 ± 0.080, 0.1036 ± 0.060, 0.5562 ± 0.125, 0.5070 ± 0.117, respectively, in triplicate, one dot represents 1 field of view. Kruskal-Wallis with Dunn’s multiple comparison post-hoc test. **C-E**) Degradation capacity of the melanoma cell lines was assessed using gelatin degradation assay allowing to visualize actin (red), cortactin (green) and degradation spots (black). **C**) Representative images of cells seeded on FITC-gelatin for 24 hours. Stained with cortactin and phalloidin, confocal microscopy. **D**) For cells displaying degradation activity, the degradation area (total area per cell) was quantified, (Mean ± SD) = 0.00, 0.00, 11.13 ± 11.00, 3.19 ± 2.76, 7.48 ± 5.59, 3.99 ± 2.97, respectively, in triplicate, one dot represents 1 cell. **E**) Degradation frequency was calculated as a percentage of cells displaying gelatin degradation activity, (Mean ± SD) = 0.00, 0.00, 6.19 ± 2.26, 4.00 ± 2.94, 26.08 ± 16.52, 20.08 ± 4.79, respectively, in triplicate, one dot represents 1 field of view. Kruskal-Wallis with Dunn’s multiple comparison post-hoc test. **F-H**) WM983B cells were seeded on gelatin 24 hours prior to the experiment, fixed and stained with cortactin, and LAMP1 antibody and with phalloidin (actin). **G**) The level of colocalization was quantified by line profiling in ImageJ. **H**) Zoom on partial colocalization between LAMP1 and gelatin degradation spots in three different regions. * p<0.05; ** p<0.01; *** p<0.001; **** p<0.0001.

**Figure S2:**
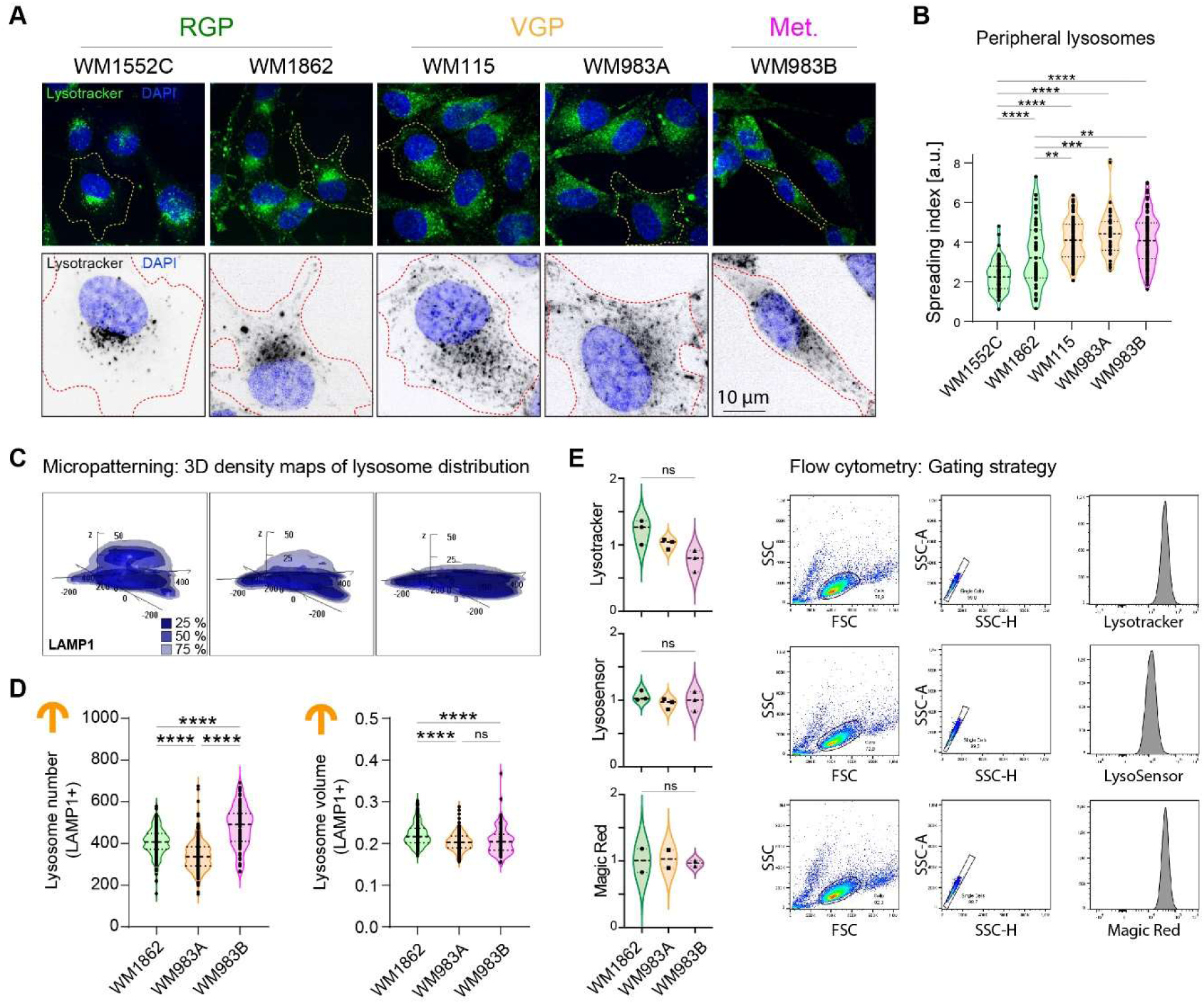
KIF1B and KIF5B drive lysosome positioning in metastatic melanoma cells. **A-B**) Lysosome positioning in non-constrained cells scales with metastatic potential in human melanoma cells. **A**) Representative images of Lysotracker staining in live imaging in RGP (WM1552C and WM1862), VGP (WM115 and WM983A) and metastatic (WM983B) cells. Similar images than in Figure 1B for WM1862, WM983A and WM983B. **B)** Graph depicting lysosome spreading index in the different cell lines (Mean ± SD) = 2.294 ± 0.8067, 3.390 ± 1.597, 4.124 ± 1.036, 4.477 ± 1.180, 4.111 ± 1.225, respectively, p values (from top) = <0.0001, <0.0001, <0.0001, <0.0001, 0.0019, 0.0003, 0.0012 **C-D**) Lysosome distribution, number and volume in melanoma cells grown on micropatterns. **C**) 3D density maps of WM1862, WM983A, WM983B cells imaged using micropatterning. Density maps of LAMP1 staining were calculated using R software, displaying the smallest area that can be occupied by 25, 50 and 75% of all compartments. **D**) Graphs showing (*left*) lysosome number (Mean ± SD) = 405.9 ± 68, 341.9 ± 74, 479.7 ± 99, respectively, and (*right*) volume (Mean ± SD) = 0.2211 ± 0.023, 0.2035 ± 0.021, 0.2071 ± 0.034, respectively, in cells grown on micropatterns. One dot represents one cell, p value <0.0001 in all significant comparisons. **E**) Graphs showing normalized fluorescence of lysosome mass with Lysotracker (*top*) (Mean ± SD) = 1.21 ± 0.18, 1.02 ± 0,08, 0.77 ± 0.16, respectively, (*middle*) lysosome acidity with Lysosensor (Mean ± SD) = 1.06 ± 0.08, 0.95 ± 0.08, 0.98 ± 0.15 respectively and (*bottom*) cathepsin B degradative activity with Magic Red, (Mean ± SD) = 1.007 ± 0.25, 1.03 ± 0.19, 0.96 ± 0.06 respectively, analyzed by flow cytometry. One dot represents the average of one experiment. One-way Anova. No comparison is statistically different. With Lysotracker, WM1862 Vs WM983B, p=0.0512. Right: Gating strategy used for each dye.

**Figure S3:**
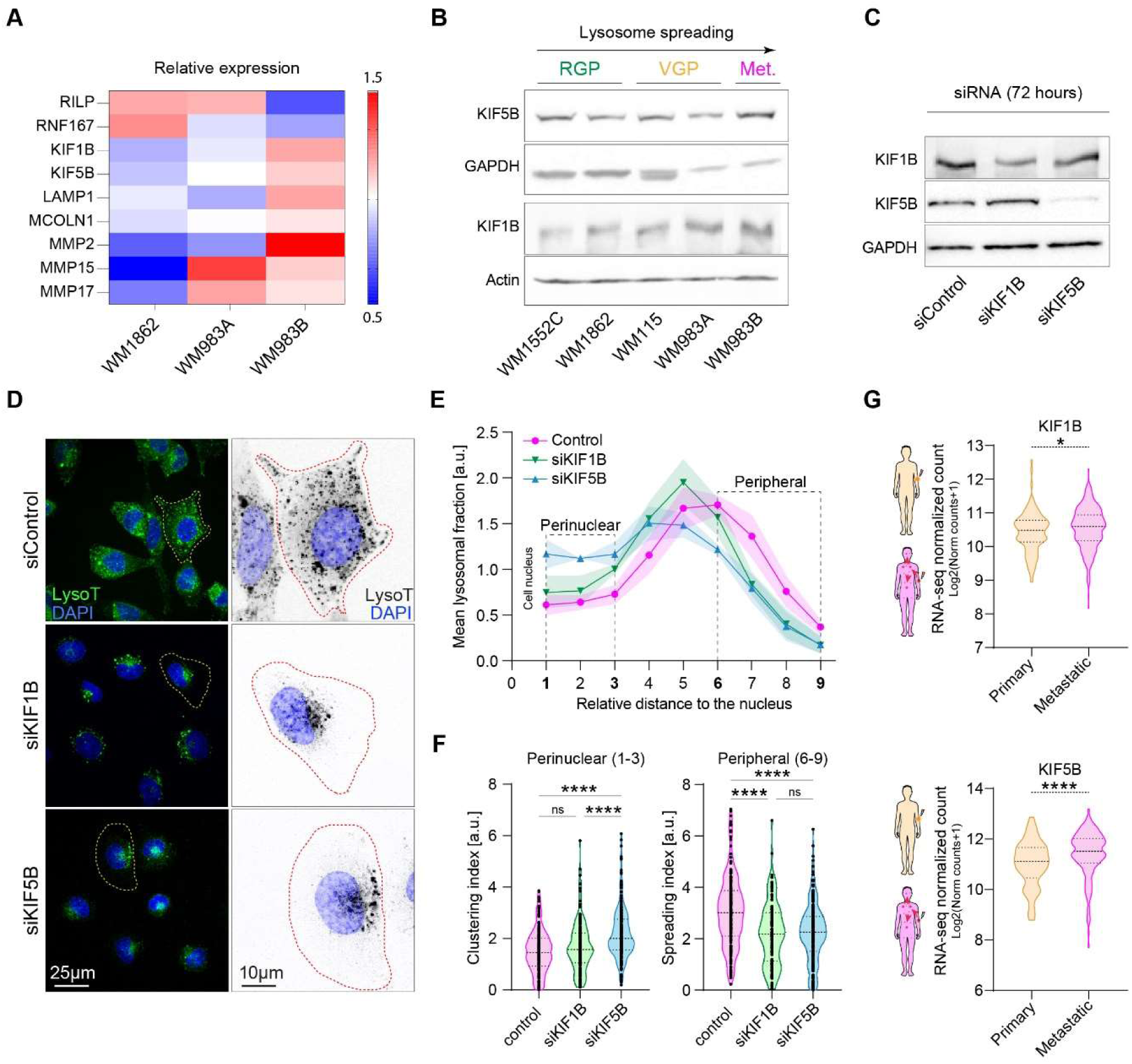
In human melanoma, KIF5B and KIF1B are overexpressed and promote lysosome spreading to cell periphery. **A-B**) KIF1B and KIF5B expression is increased in metastatic melanoma cells, as shown by RNA sequencing together with (**A**) a selection of genes, color code indicates relative expression levels. All genes show a statistically significant difference between WM1862 and WM983B with pAdj<0.0001, except for MCOLN1 where pAdj=0.026. **B**) Western blot analysis of KIF5B, KIF1B expression in patient-derived cell lines. **C-F**) KIF1B and KIF5B promote lysosome spreading in metastatic melanoma cells. **C**) Expression of KIFs upon siRNA treatment. **D**) Representative images of WM983B cells treated with control, KIF1B or KIF5B siRNAs and imaged for lysosomes positioning by live spinning disk imaging using lysotracker. **E**) Lysosome positioning relative to the nucleus shows an increase of lysosomes in the perinuclear area and a decrease in the peripheral region upon KIF1B or KIF5B depletion. **F**) Quantification of (left) Perinuclear index (Mean ± SD) = 1.505 ±0.830, 1.696 ± 0.983, 2.198 ± 0.9996 and (right) Peripheral index (Mean ± SD) = 3.051 ± 1.358, 2.185 ± 1.210, 2.222 ± 1.128, 2.294 ± 0.8067, respectively. One dot represents one cell. **G**) Analysis of sequencing data from tumors of melanoma patients. KIF5B (top) and KIF1B (bottom) show increased expression in metastatic cohort compared to primary melanoma TCGA cohort. * p<0.05; ** p<0.01; *** p<0.001; **** p<0.0001.

**Figure S4:**
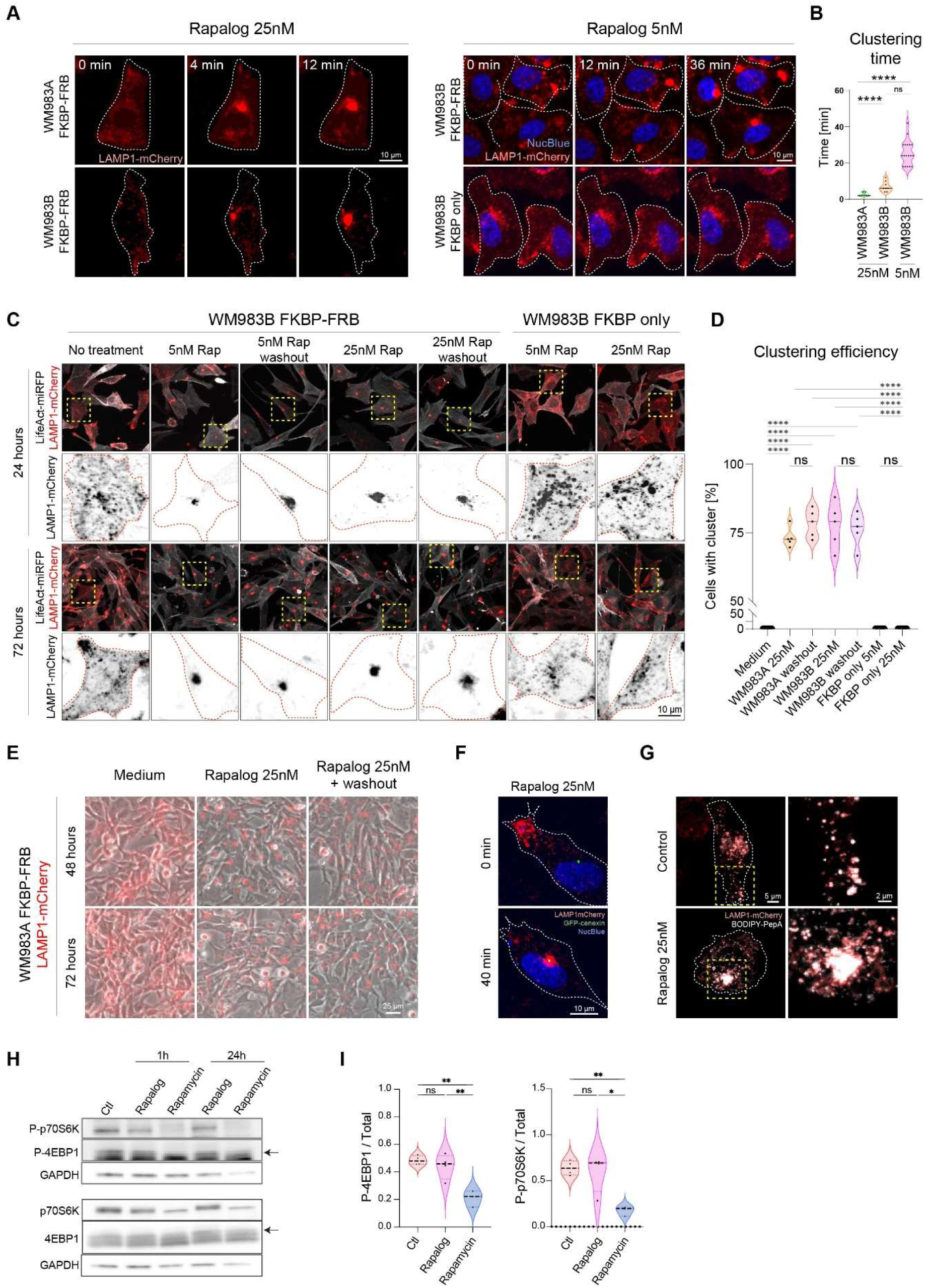
Fast and stable Rapalog-induced lysosome perinuclear clustering. **A)** Rapalog induces a fast lysosome perinuclear clustering. Representative images of WM983A and WM983B cells expressing two heterodimerizing domains (FKBP-FRB) and WM983B cells expressing only one heterodimerizing domain (FKBP only) extracted from a 1-hour time-lapse. After acquiring the first time-frame, cells were treated with 25 nM Rapalog (left) or with 5 nM Rapalog (right). **B**) Graph showing clustering timing at different Rapalog concentrations. Cluster formation was quantified as the time needed until a visible cluster appeared. Time [min] = 2.6 ± 1.0, 7.0 ± 2.8, 25.7 ± 6.8, respectively. One dot represents 1 cell, one-way ANOVA, p values (from top): <0.0001, 0.2541, <0.0001. **C-E**) Rapalog treatment induces stable lysosome perinuclear clustering. Representative images of C) WM983B and E) WM983A (FKBP-FRB) cells treated with 5nM or 25 nM Rapalog and imaged at 48 hours and 72 hours timepoints. Washout condition – cells were treated for 1 hour with Rapalog, washed 3x in PBS and cultured in normal growth medium for the duration of the experiment. **D**) Analysis of C-E showing percentage of cells with a lysosome cluster, one-way ANOVA, p values <0.0001 when significant. **F**) Rapalog induces lysosome clustering around the microtubule organizing center, labelled by cenexin-GFP. **G**) Cargo delivery to lysosomes is not perturbed by Rapalog treatment. WM983B (FKBP-FRB) cells expressing LAMP1-mCherry (red) were cultured in medium or in medium with 25 nM Rapalog for 1 hour and then treated with PepstatinA-BODIPY (white) probe for 45 minutes. Cells were imaged live using spinning disc. **H**) Western blots against phosphorylated (upper) and total (lower) forms of mTORC1 targets p70S6K and 4EBP1 (arrow) on extracts from WM983B cells treated with Rapalog or Rapamycin (5nM each) for 1h or 24h and (**I**) graphs showing quantification of phosphorylated over total forms at 24h, n=4, one dot represents one experiment, One way Anova, p values (from top, left): 0.019, 0.3915, 0.0036 and 0.0070, 0.6541, 0.0084, respectively. * p<0.05; ** p<0.01; *** p<0.001; **** p<0.0001.

**Figure S5:**
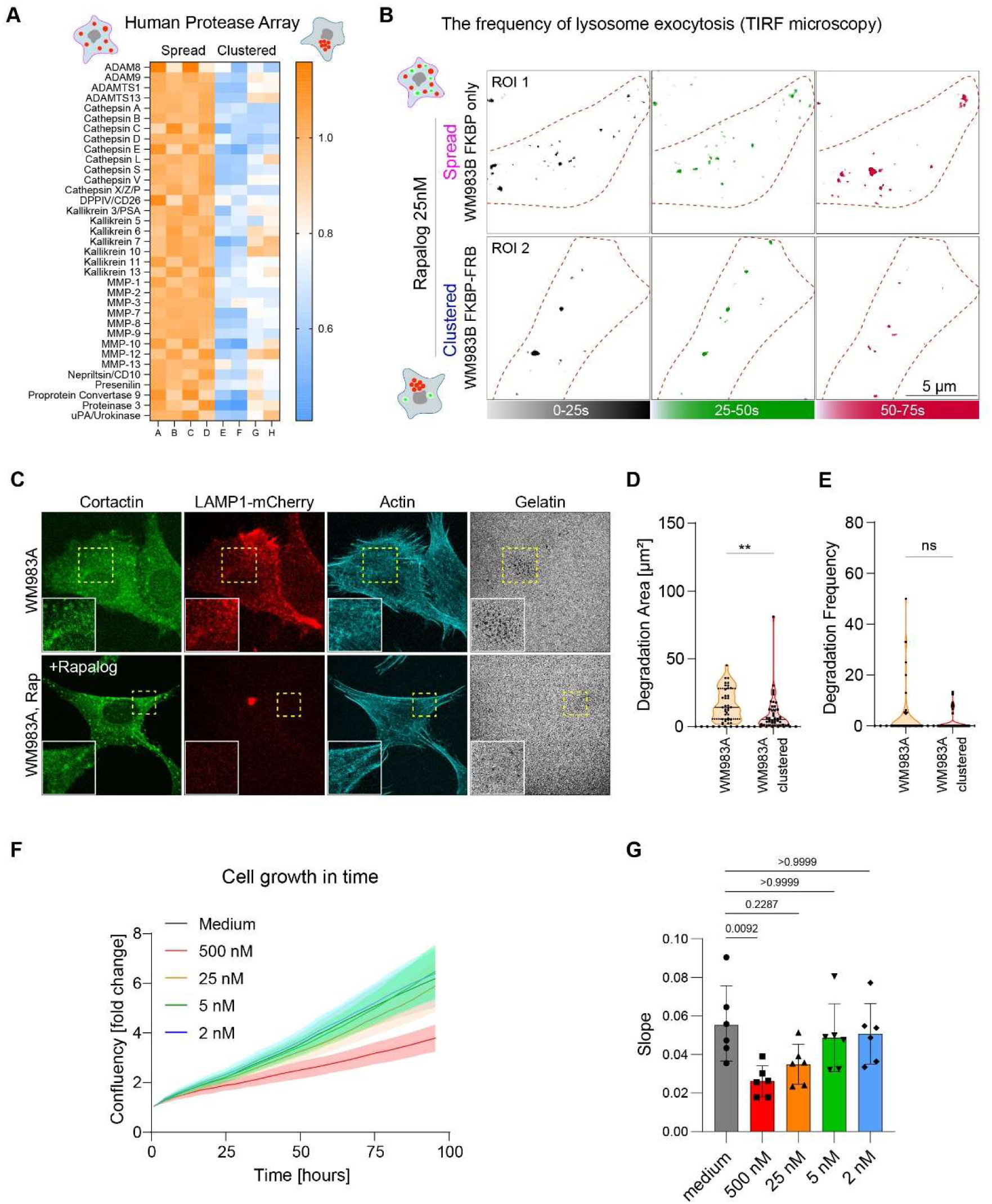
Peripheral lysosomes promote secretion of proteases. **A)** Lysosome clustering impairs secretion of proteases, quantified from conditioned medium by Human protease array. Color code indicates relative secretion levels. **B**) Quantification of cell secretion by TIRF microscopy showing the number of secretion events over time. **C-E**) Lysosome clustering impairs gelatin degradation. **C**) Representative immunofluorescence images of WM983A cells. **D**) Degradation Area (Mean ± SD) = 16.94 ± 11.71 n= 36 cells, 9.38 ± 13.47 n= 45 cells, in triplicate. **E**) Degradation frequency (Mean ± SD) = 3.62 ± 9.73 n= 57 cells, 1.49 ± 3.44 n= 58 cells, in triplicate. **F**) Proliferation rate of WM983B cells treated with increasing concentrations of Rapalog was analyzed using incucyte. Cell confluency was quantified for each time-point, all values were normalized to time 0 and the growth curves were plotted. Colored lines depict different culture conditions: medium (black), 2 nM (blue), 5 nM (green), 25 nM (orange), 500 nM Rapalog (red) and are representing average value of three experiments ± SD. **G**) Slopes were calculated for the different growth conditions. Slope (Mean ± SD) = 0.0552 ± 0.02, 0.0261 ± 0.01, 0.0348 ± 0.01, 0.0486 ± 0.02, 0.0507 ± 0.02, respectively. Kruskal-Wallis test with Dunn’s multiple comparison post-hoc test, respective p-values are shown in the graph. One dot represents one experiment (performed in technical triplicate). *p<0.05; ** p<0.01; *** p<0.001; **** p<0.0001.

**Figure S6:**
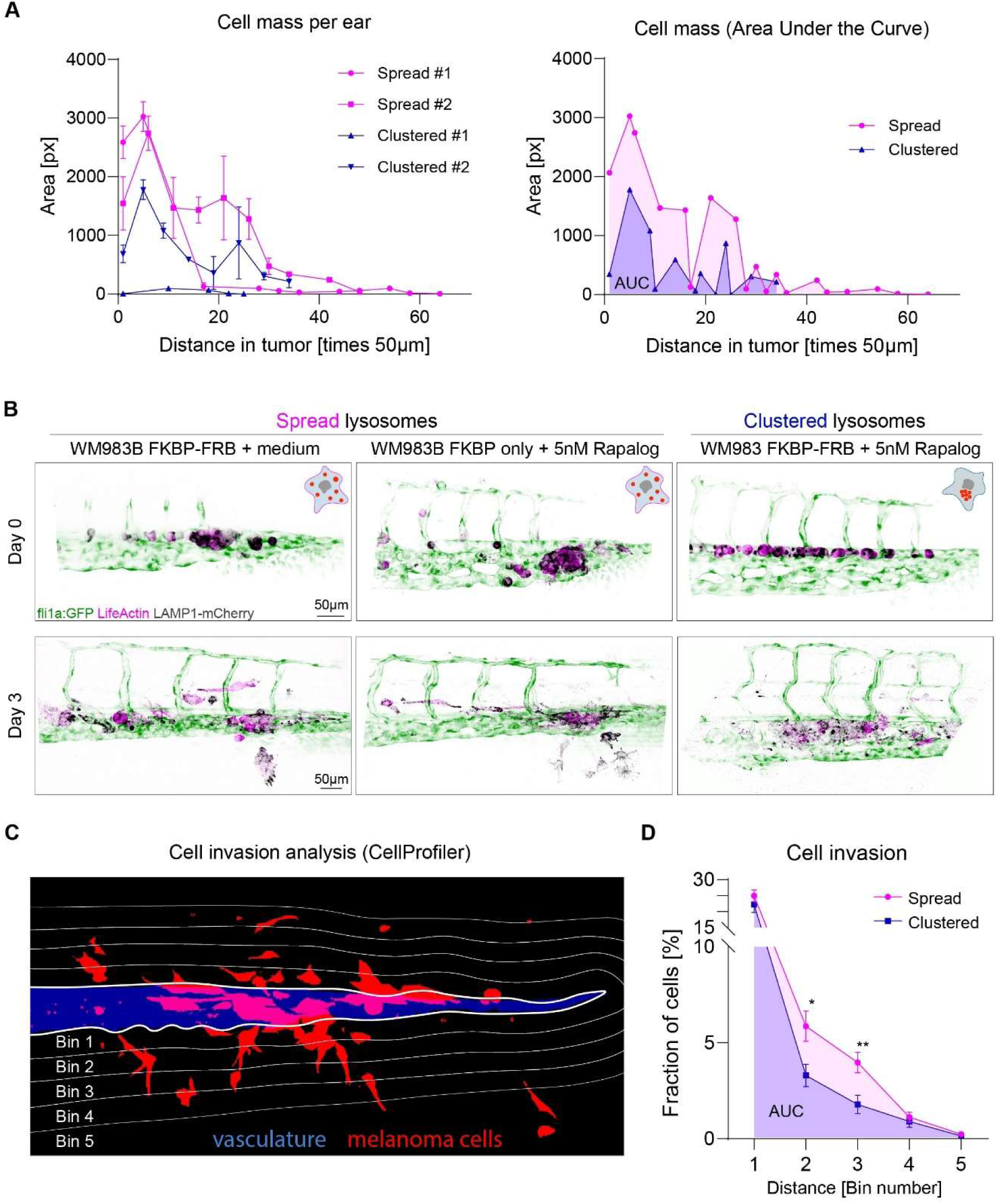
Monitoring cell invasion *in vivo*. **A**) Lysosome clustering prevents tumor progression in mice, magenta = spread lysosomes n= 2 (FKBP only, 5nM Rap), blue = clustered lysosomes n=2 (FKBP-FRB, 5nM Rap). (*left*) Cell mass in the individual ears, AUC ± SD = 33 046 ± 1 789, 51 713 ± 3 766, 1 264 ± 159, 24 537 ± 2 517, respectively, and (*right*) mean cell mass per condition, AUC (Mean ± SD) = 42 380 ± 13 200, 12 901 ± 16 456, respectively. **B**) Representative immunofluorescence images at day 0 and day 3 post-injection in zebrafish embryos, green= Fli1a:GFP, black= LAMP1-mCherry, magenta= LifeAct-miRFP670. **C**) Quantification of the post-extravasation invasion potential at 72 hours (day 3) post injection in zebrafish embryos. Representative example of the segmented image. Z-projections of vasculature (blue) and WM983B melanoma cells (red) were segmented using ImageJ. Segmented images were further analyzed using CellProfiler: Image was divided into five regions (bins) with increasing distance from the vasculature region and the total cell area in each region was quantified. **D**) Percentage of tumor cells in zebrafish embryos (cell area per bin / total cell area) is plotted for both conditions. Magenta = cells with spread lysosomes, blue= cells with clustered lysosomes. Each line represents mean value from three independent experiments ± SD. Percentage (scattered) = 24.9 ± 10.9, 5.9 ± 4.7, 3.9 ± 3.2, 1.1 ± 1.6, 0.2 ± 0.6, respectively, n= 35 embryos. Percentage (clustered) = 22.2 ± 9.5, 3.3 ± 2.3, 1.8 ± 1.9, 0.9 ± 1.2, 0.1 ± 0.3, respectively, n = 15 embryos, p values (from left) = 0.3760, 0.0121, 0.0043, 0.5745, 0.5363, Multiple unpaired t-tests. *p<0.05; ** p<0.01; *** p<0.001; **** p<0.0001.

## Methods

### Antibodies

Anti-Cortactin (p80/85) clone 4F11 (ref. n°05-180-I) and Anti-LAMP1 (ref. n°L1418) are from Merck (Sigma-Aldrich) and anti-SOX10 (ref. 383R-10) is from Cell Marque. Alexa Fluor^TM^ 488 Phalloidin (ref. n° A12379), Alexa Fluor^TM^ 568 Phalloidin (ref. n°A12380), Alexa Fluor^TM^ 647 Phalloidin (ref. n°A22287) and Alexa Fluor–conjugated secondary antibodies, are all from Thermo Fisher Scientific. Secondary antibodies include Alexa Fluor^TM^ 405 conjugated Goat anti-Rabbit (ref. n°A-31556), Alexa Fluor^TM^ 488 conjugated Goat anti-Rabbit (ref. n°A-11034), Alexa Fluor^TM^ 555 conjugated Goat anti-Rabbit (ref. n°A21429) & Alexa Fluor^TM^ 647 conjugated Goat anti-Rabbit (ref. n°A-21244).

### Cell Culture

Primary human melanoma cell lines were purchased from Rockland and cultured in MCDB153 (ref. P04-80062, Dutscher) and Leibovitz’s L-15 medium (ref. n°11415064, Thermo Fisher Scientific) in a 4 to 1 ratio, supplemented with 2% foetal bovine serum, 1.68 mM CaCl_2_ and 1% penicillin/streptomycin. WM1552C and WM1862 are non-tumorigenic (RGP) primary human melanoma cell line without metastatic capability. WM115 and WM983A are tumorigenic (VGP) human primary melanoma cell line. WM983B is a human metastatic melanoma cell line derived from the same patient as WM983A. All cell lines are mutant for BRAF ^V600^.

### Western blot

Cell extracts were prepared in presence of phosphatase inhibitors (Ref.A32961, Thermo Fisher Scientific) when needed and denatured in Laemmli buffer and incubated at 95°C for 10 min. 40 µg of protein extract were loaded on 4–20% polyacrylamide gels (Bio-Rad Laboratories, Inc). The following antibodies were used: Rabbit anti-KIF1B (Ref. A301-055A, Thermo Fisher Scientific), rabbit anti-KIF5B (Ref. ab167429, Abcam), mouse anti-GAPDH (Ref. MAB374, Millipore), rabbit anti-4EBP1 (Ref. 9644T, Cell Signaling), rabbit anti-Phospho-4EBP1 (Ref. 2855, Cell Signaling), rabbit anti-p70S6K (Ref. 2708T, Cell Signaling), rabbit anti Phospho-p70S6K (Ref. 9234, Cell Signaling) and Secondary horseradish peroxidase-conjugated antibodies: anti-Mouse (Ref. 715-035-151, Interchim) and anti-rabbit (Ref. 111-035-003, Jackson ImmunoResearch). Acquisitions were performed using iBright 1500 (Thermo Fisher) imager. Intensities were measured using the Fiji software.

### Lentivirus transduction and plasmid transfection

#### Transduction

pLSFFV-LAMP1-mCherry-FKBP (FK506 binding protein), pLSFFV-BicD-FRB (FKBP-rapamycin binding domain) and pLSFFV-LifeActin-miRFP lentivirus were produced in HEK293T cells using JetPRIME^®^ transfection reagent (Polyplus). WM983B cells were infected by lentivirus in the presence of 5μg/mL polybrene (ref. n°TR-1003, Merck/Sigma-Aldrich) followed by antibiotic selection (puromycin 1μg/mL, blasticidin 5μg/mL, hygromycin 200μg/ml). *<u>Transfection</u>:* Cenexin-GFP plasmid was transfected using JetOPTIMUS^®^ transfection reagent (Polyplus) in a live-video dish, 46h prior experiment.

#### siRNA silencing

Cells were reverse-transfected with siRNA from Dharmacon (30nM; ON-Targetplus human KIF1B (L-009317-00), human KIF5B (L-008867-00) and Non-targeting Pool (D-001810-10) using Lipofectamin RNAiMAX (ThermoFisher Scientific, Ref. n°13778150) and analyzed after 72h for knockdown efficiency by western blot and for lysosome positioning with Lysotracker by confocal imaging as described below.

#### Invasion assay

The 3D collagen invasion assay was adapted from (Sadok et al., 2015). Briefly, melanoma cell lines at 3×10^5^ cells/ml were labelled with Hoechst 33342 and suspended in 3mg/mL of serum free solution of neutralized Type I Bovine Collagen (PureCol^®^ 5005-B, Advanced Biomatrix). Then, 600μL were distributed into black 24 well plates (ref. n°058062, Dutscher) coated with bovine serum albumin or in 4-chamber glass bottom dishes (Cellvis). The plates were centrifuged at 1500 rpm at 4°C for 5min and incubated at 37°C for 2 hours. Once collagen had polymerized, medium supplemented with 10% foetal bovine serum, 100ng/mL EGF was added on top of the collagen. 24h after, cells were observed using a Leica TSC SPE confocal microscope (x20 HCX Pl Apo 0.7 NA objective, Wetzlar, Germany) and z-stacks were acquired. Four fields per sample were imaged. Nuclear localization was quantified by IMARIS (interactive microscopy image analysis software) at each plane. The invasion index was calculated as the number of nuclei above 10μm (20μm for experiment in Fig 2K-N) divided by the total number of nuclei by field.

#### Matrix degradation assay and immunofluorescence

Glass coverslips were coated with fluorescent-labelled-FITC gelatine or Cy3 gelatine as described previously (Kolli-Bouhafs et al., 2014). Then, previously starved melanoma cells expressing or not LAMP1-mCherry-FKBP and BicD2-FRB were plated on fluorescent gelatine and incubated at 37°C for 6 to 24h in medium supplemented with 10% foetal bovine serum with or without Rapalog. Cells were fixed using 4% paraformaldehyde permeabilized using triton-X-100 at 0,1% and incubated in 2% of bovine serum albumin at room temperature. Cells were then labelled for 1h with Anti-Cortactin (1/250). After three washes with PBS, cells were incubated with either Alexa Fluor^TM^ 405 conjugated Goat anti-Rabbit (1/1000), Alexa Fluor^TM^ 488 conjugated Goat anti-Rabbit (1/1000) or Alexa Fluor^TM^ 647 conjugated Goat anti-Rabbit (1/1000) and Alexa Fluor^TM^ 568 Phalloidin (1/250) or Alexa Fluor^TM^ 647 Phalloidin (1/250) for 1h, washed and mounted in ProLong^TM^ Gold Antifade Mountant (ref. n°P10144, Thermo Fisher Scientific). Cells were imaged using Leica TSC SPE or SP8 confocal microscope (x63 HCX Pl Apo 1.40 NA or x20 HCX Pl Apo 0.7 NA objective, Wetzlar, Germany). Invadopodia were identified as actin and cortactin rich punctate structure. Areas of degradation were identified as “black holes” within the fluorescent gelatine. Invadopodia and areas of degradations were quantified using ImageJ software. Degradation areas measurements were based on cells displaying degradation activity, and the frequency of degradation was based on randomly selected cells. Maximum filter, background subtraction and gaussian blur filters were then applied to extract the gel degradation areas by thresholding. Then, M1 Manders coefficients between LAMP1 and inverted Gelatin intensity pictures for these selections were calculated using the colocalization finder plugin.

### Clustering dynamics, washout experiments & proliferation assay

#### Mechanism of lysosome clustering

clustering was performed by heterodimerization between LAMP1-mCherry-FKBP (lysosomes) and BicD-FRB (dynein adaptor) by the use of Rapalog (A/C Heterodimerizer ref. n°635057, from Takara Bio Inc.). Cells were cultured in glass bottom dishes and imaged at different time points (24, 48 and 72 hours) using Olympus Spinning Disk (60X objective, N.A. 1.2). To establish time needed for lysosome clustering, WM983A and WM983B cells expressing both heterodimerizing domains (FKBP-FRB) were treated with 5nM or 25nM Rapalog and followed in time. Time to appearance of lysosome cluster was counted.

#### Washout experiments

WM983B cells expressing either one (FKBP only) or two (FKBP-FRB) heterodimerizing domains were cultured in medium supplemented with 5nM or 25nM Rapalog. To assess the stability of clustering, cells were treated for 1h with 5nM or 25nM Rapalog, washed 3x in PBS and cultured in normal medium for the duration of the experiment. Cells cultured in normal growth medium were used as a control.

#### Proliferation assay

proliferation rates of WM983B cells treated with increasing concentrations of Rapalog were analyzed using the Incucyte® Live-Cell Analysis System. Confluences were automatically calculated by the Incucyte® software based on bright field images, all values are normalized to time zero. Acquisition was performed for 96h.

### Micropatterning & Immunofluorescence

Micropatterns were prepared using photo-lithography methods as previously described (Duong et al., 2012; Schauer et al., 2010). Briefly, cover slides were cleaned in EtOH, dried, cleaned with UV for 10 minutes, coated in PLL-PEG and exposed to UV 10 minutes through a Photomask with 36μm crossbow micropatterns. Coverslips were coated in Fibronectin (40μg/ml – ref. n°F1141 Merck/Sigma-Aldrich). WM1862, WM983A, WM983B cells were trypsinized, resuspended in 5ml culture medium and seeded an micropatterned coverslips. Cells were let to spread for 4 hours before fixing them in 4% PFA. For micropatterning, cells were fixed in 4% PFA, washed 3x in PBS, permeabilized in saponin 0.5% / 2% BSA in PBS for 20 minutes, blocked 30 minutes in 2% BSA, stained with Anti-LAMP1 (ref. n°L1418) primary antibody for 1 hour, washed 3x in PBS, stained with Alexa Fluor^TM^ 555 secondary Goat anti-Rabbit (1:500) and Alexa Fluor™ 488 Phalloidin (1:200) 45 minutes. Mounted in Fluoromount-G^TM^ mounting medium (ref. n°00-4958-02, Thermo Fisher Scientific with DAPI. High resolution, volume imaging was achieved using Olympus Spinning Disk (60X objective, N.A. 1.2) / Delta Vision – Advanced Precision (100x oil objective, U PL S APO, NA 1.4) with CoolSNAP HQ2 CCD camera (1392*1040 pixels). Large z-stacks were acquired for the full cell volume (0.2μm step between layers). Image analysis and processing were performed using the Fiji (1.51q) (Schindelin et al., 2012), Metamorph and R software, allowing for precise calculation of lysosome volume and number.

### Patient samples

#### Samples from tumor biopsies

Samples were provided by the Laboratoire d’Histopathologie Cutanée de la Clinique Dermatologique des Hôpitaux Universitaires de Strasbourg. Patients consent was collected and the project was approved by the Ethic committee from the Hôpitaux Universitaires de Strasbourg (CE-2024-79). There was no selection based on gender, race, ethnicity or age.

#### TCGA data analysis

The Cancer Genome Atlas Program (TCGA). We analyzed data set from TCGA SKCM database containing 333 primary and/or metastatic melanomas from 331 patients (Akbani et al., 2015) using UCSC Xena platform (Goldman et al., 2020).

### Patient Biopsies: Immunohistochemistry, Imaging, Quantification

Paraffin-embedded 4μm sections were rehydrated (toluene - ethanol - PBS), followed by antigen retrieval (5min boiling in 10mM sodium citrate #C9999 Sigma-Aldrich), permeabilized 2× 10 min in PBS (0.3% Triton X100), blocked 1h (3% BSA, 20mM MgCl2, 0.3% Tween 20, 5% FBS) and incubated with primary antibodies (LAMP1 #L1418, SOX10 #383R-14) overnight at 4°C, followed by 1hr staining with secondary antibodies (1:250 dilution) and mounted in Fluoromount DAPI. Full tissue section was imaged using Slide Scanner (Olympus, 20x objective).

Quantification was done in a blinded setup, analyzing 23 samples of patient biopsies. Ten regions of interest were randomly selected per each patient sample, reaching 240 individual regions, which were randomized and then a score between 1 and 4 was given to each region based on the level of lysosome spreading (1 = 0 – 25% spread, 2 = 25 – 50% spread, 3 = 50 – 75% spread, 4 = 75 – 100% spread). Average score per patient sample was calculated and after that samples were annotated and plotted in the graph showing Healthy skin: 4 patients, Nevus: 7 patients, Primary melanoma: 6 patients, Metastatic site: 7 patients.

### Live cell imaging

For live cell imaging, wild-type patient-derived cells treated with 50nM Green Lysotracker (Molecular Probes #L7526) or WM983A and WM983B cells, stably expressing LAMP1-mCherry and LifeAct-miRFP were plated on fibronectin-coated (Sigma-Aldrich F1141, 10μg/ml in water) 4-chamber glass bottom dishes (35 mm, #1.5, Cellvis) or on fluorescent gelatin-coated coverslips mounted in a Ludin Chamber (Life Imaging Services), respectively.

#### Gelatin degradation

The cells were placed at 37°C, 5% CO_2_ on an iMIC microscope equipped with a multi-LED Lumencor Spectra X. Images were acquired with an Olympus 60x TIRFM (1.45 NA) objective every 2.5min during 4h and a Hamamatsu Flash 4 V2+ camera (Iwata) piloted by the Live Acquisition software (Till Photonics). Expressing cells were initially located via both the mCherry and LifeAct-miRFP signals, and were subsequently followed via dual phase contrast/fluorescent signal together with the FITC-coupled gelatin substrate. 10 to 20 different fields were sequentially recorded during each experiment using a Marzhauser Motorized Stage piloted by the iMIC software. JeasyTFM software were then used for automatic selection and repositioning of the best focused images in all channels and time-points. Mean fluorescence intensity of actin, Lamp1 and the underlying substrate at each time points were calculated at invadopodia identified using gelatin degradation.

#### Lysosome radial distribution

The cells were placed at 37°C, 5% CO_2_ chamber on an Olympus Spinning Disk (60X objective, N.A. 1.2) and full cell volume z-stacks (0.5μm step) were acquired and then processed in ImageJ (1.53t) and CellProfiler (4.2.1) (Stirling et al., 2021). Analysis was done on Maximum intensity projections, cell outline was manually segmented in ImageJ, nuclei were segmented based on thresholding of DAPI channel in ImageJ. Binary masks were processed in CellProfiler and radial distribution was calculated using green lysotracker signal.

### TIRF microscopy

WM983B cells expressing FKBP only (spread) and WM983B cells expressing both FKBP and FRB (clustered) were mixed in a 1:1 ratio, seeded in fibronectin-coated low glass bottom μ-Dish 35mm (ref. n°80137, Ibidi) 48h prior to imaging, treated with 25nM Rapalog for 1h. Imaging was performed in culture medium using an inverted Leica DMI8 microscope (objective 100X HC PL APO 1,47 oil). Recording was done with an Evolve^®^ 512 camera (for TIRF-HILO), at 512X512 pixels resolution, at an acquisition rate of 250ms between frames, for a total duration of 90 seconds, with AFC (Adaptive Focus Control). Exocytosis events were identified based on VAMP7-pHluorin signal, marking lysosome exocytosis (Lachuer et al., 2023). Secretion events were detected and counted manually.

### Human protease array

Cells were seeded on collagen, cultured for 48h in serum-free medium and conditioned medium was analyzed using a human protease array (RnDSystems, ARY021B), following the manufacturer’s instructions.

### Flow Cytometry

Cells were seeded at a density of 1*10^5^ cells per well in 96-well plates (Falcon® 353077) and labeled 30 minutes at 37°C 5% CO_2_ with either 50nM LysoTracker™ Green DND-26 (Invitrogen™ L7526), 1µM LysoSensor™ Green DND-189 (Invitrogen™ L7535) or Magic Red® Cathepsin B (1:26 dilution, IMC 937) diluted in 100µL of culture medium. Cells were washed with PBS, resuspended in 300µL of BSA 2% in PBS and analyzed by flow cytometry (Attune NxT, Invitrogen™). Geographic mean fluorescent intensities were determined with FlowJo V10 (LLC).

### Experimental metastasis assay in mouse

All animals were housed and handled according to the guidelines of INSERM and the ethical committee of Alsace (CREMEAS), following French and European Union animal welfare guidelines (Directive 2010/63/EU on the protection of animals used for scientific purposes. Mouse facility agreement number: #C67-482-33; APAFIS 43901-2023062112024760. Six to eight week-old immunodeficient nude mice (Crl:NU(NCr)-Foxn1nu; Charles River) were used in all experiments. Mice were injected subcutaneously with 250,000 WM983B cells of different lysosome clustering status (FKBP only spread; FKBP-FRB clustered) diluted in 20μl PBS and previously labelled with eBioscience eFluor 670 (ThermoFisher Scientific, ref. 65-0840-85) and treated with 5nM Rapalog 1h before injection. Subsequently, mice were treated every day with local application of Rapalog (5nM) on the ear. At day 11, ears were harvested, imaged by fluorescence, embedded in OCT (Cellpath) and frozen at -80°C. 7μm transversal ear sections were performed with a cryostat, mounted with Fluoromount-GTM with DAPI (ThermoFisher Scientific, ref. 00-4959-52). Imaging was performed using a slide scanner (Olympus, 20X objective) and area covered by tumor cells (Cell Trace fluorescence) was measured in ImageJ and plotted over the section number (=distance in tumor) using GraphPad Prism.

### Experimental metastasis assay in zebrafish

All zebrafish (ZF) procedures were performed in accordance with French and European Union animal welfare guidelines and supervised by local ethics committee (ZF facility A6748233; APAFIS 2018092515234191). Tg(fli1a:eGFP) (Lawson and Weinstein, 2002) embryos were maintained in Danieau 0.3X medium (17.4mM NaCl, 0.2mM KCl, 0.1mM MgSO_4_, 0.2mM Ca(NO_3_)_2_) buffered with HEPES 0.15mM (pH=7.6), supplemented with 200μM of 1-phenyl-2-thiourea (PTU, ref. n°P7629, Merck/Sigma-Aldrich) to avoid pigmentation. Two days post-fertilization (2dpf) embryos were mounted in 0.8% ultrapure low melting point agarose (Invitrogen) containing 0.17mg/ml tricaine (ethyl-3-aminobenzoate-methanesulfonate, ref. n°E10521, Merck/Sigma-Aldrich). WM983B cells of different lysosome clustering status (spread, clustered) were injected with a Nanoject II Auto-Nanoliter Injector (Drummond Scientific Company) and microforged glass capillaries (25 to 30μm inner diameter) filled with mineral oil (ref. n°M5904, Merck/Sigma-Alrich). 13.8nL of cell suspension from confluent T25 flasks (50×10^6^ cells per ml approx.) were injected in the duct of Cuvier under a M205 FA Fluorescence stereomicroscope (Leica), as previously described (Stoletov et al., 2010). Embryos were injected with WM983B cells or with WM983B cells treated with 5nM rapalog 1h prior to injection, and then kept in Danieau with PTU. Caudal plexus was recorded at day 0 (injection day) and 3dpi using the inverted spinning-disk Olympus IXplore Spin, 30x / 1.05 NA (silicone) objective. Z-stacks of the caudal plexus were acquired for each embryo (3μm or 5μm between layers, at day 0 and 3dpi respectively) with the following settings: 488nm laser at 2% for 100ms / 561nm laser at 15% for 300ms / 640nm laser at 15% for 300ms). Detailed lysosome status (Fig.4b) was imaged at 3 hpi using a 60x/1.2 NA (water) objective.

### Correlative Light and Electron Microscopy (CLEM)

WM983B cells expressing LAMP1-mCherry-FKBP and BicD2-FRB were cultured in control medium, or in medium supplemented with 25nM Rapalog for one hour and imaged with an Olympus Spinning Disk (60X objective, N.A. 1.2). The samples were then chemically fixed right after the photonic acquisition with 0,05% malachite green, 2.5% glutaraldehyde in 0.1M sodium cacodylate buffer (NaCac), pH7.4 during 30min in an ice bath. Subsequently the samples were post-fixed in 1% OsO_4_ - 0.8% K_3_[Fe(CN)_6_] - 0.1M NaCac buffer pH7.4 (under a fume hood) kept in an ice bath for 50 min, and then washed 2 times with in ice-cold 0.1M NaCac. Then the samples were incubated in 1% aqueous tannic acid solution for 25 min in an ice bath and finally washed 5 times with distilled water. Samples were then kept in 1% uranyl acetate aqueous solution overnight at 4°C sheltered from the light. The samples were serial dehydrated with ethanol solutions (25%, 50%, 70% 95% and 100%). Subsequently the samples were incubated in a serial resin-ethanol 100% mix (1:3; 1:1; 3:1), ending with an incubation in 100% Epon resin 3 times 1h at room temperature. The samples were allowed to polymerize in an oven at 60°C for 48h. The resin blocks were trimmed by ultramicrotomy, 90nm thin sections were collected and placed in copper/formvar slot grids. The transmission electron microscopy (TEM) data sets were acquired with a Hitachi 7500 TEM, with 80 kV beam voltage, and the 8-bit images were obtained with a Hamamatsu camera C4742-51-12NR. Correlative light and electron images were obtained/combined using Adobe Photoshop v.24.4.

### Transmission electron microscopy

WM1862, WM983A, and WM983B cells were seeded on ACLAR^®^ films (EMS) and were fixed in primary fixation buffer containing 2.5% glutaraldehyde (EMS) and with 0,05% malachite green (Sigma-Aldrich) in 0.1M cacodylate buffer (EMS) for 30 minutes on ice. Following post-fixation were processed in 1% OsO_4_ and 0.8% K_3_[Fe(CN)_6_] in 0.1M cacodylate buffer on ice for 50 min, and then incubated in 1% aqueous tannic acid solution for 25 min. Cells were stained with 1% uranyl acetate, and then dehydrated in sequential gradient alcohol baths and infiltrated with Epon resin. The resin polymerization is cured in an oven at 60°C for 48 h. Ultrathin sections (90 nm) were obtained by Leica ultramicrotome, and collected serial sections on a copper grid for TEM observation. Finally, the ultra-sections were post-stained by UranyLess (EMS) for 10 minutes, and rinsed several times with H_2_O followed by 3% Reynolds lead citrate (EMS) for 10 minutes. Micrographs were obtained at 80 kV in Hitachi 7500 TEM with Hamamatsu camera C4742-51-12NR digital camera. Area of protrusions were analysis by imageJ (Fiji), and manually calculate the containing number of endo-lysosome.

### Image analysis

Organelles segmentation was performed using MetaMorph Microscopy Automation and Image Analysis Software (Molecular Devices) and the ImageJ Modular Image Analysis (MIA) plugin. Segmentation on LAMP1 images were performed to get coordinates for individual LAMP1+ objects and their number. 3D Density maps and inter-organelle distance and distance to barycenter were obtained through the use of R software. Codes are available on GitHub (https://github.com/KJerabkovaRoda/Lysosome_positioning).

Cell invasion in zebrafish was performed by ImageJ & Cell Profiler (Molecular Devices) software. Zebrafish Z-projection images were divided into 6 regions – plexus and 5 regions outside (bins). Area of cells was quantified for each region using Cell Profiler and percentage of cells (from total) was calculated per each region and plotted in a graph. Invasion potential was calculated as area under the curve for all cells that extravasated and invaded outside of the vasculature region.

### Mass Spectrometry - Quantitative Proteomics

Conditioned medium was produced by WM983B FKBP-FRB +/- Rapalog (5nM) in serum free medium. After 24h, supernatant was collected, centrifuged at 300g for 10min and concentrated using a 3kDa filter. A Pierce^TM^ 660nm protein assay quantification (ref. n°22660, Thermo Fisher Scientific), 2µg of each protein extract were digested using the automated Single Pot Solid Phase enhanced Sample Preparation (SP3) protocol as described in (Hughes et al., 2019) on the Bravo AssayMAP platform (Agilent Technologies). Extracted peptides were cleaned-up using automated C18 solid phase extraction on the same platform and analysed by nanoLC-MS/MS on a nanoUPLC system (nanoAcquityUPLC, Waters) coupled to a quadrupole-Orbitrap mass spectrometer (Q-Exactive HF-X, Thermo Scientific). Chromatographic separation was conducted over a 60 minutes linear gradient from 2 to 40% of solvent B (0.1 % formic acid in acetonitrile) at a flow rate of 350 nL/min. A Top 10 method was used with automatic switching between MS and MS/MS modes to acquire high resolution MS/MS spectra. To minimize carry-over, a solvent blank injection was performed after each sample.

NanoLC-MS/MS data was interpreted to do label-free extracted ion chromatogram-based differential analysis. Searches were done using Mascot software (version 2.5.1, MatrixScience) against a composite database including *Homo sapiens* and *Bos taurus* protein sequences, which were downloaded from UniProtKB-SwissProt (28-07-2021; 26.031 sequences, Taxonomy ID: 9913 and 9606 respectively) to which common contaminants and decoy sequences were added. One trypsin missed cleavage was tolerated. Carbamidomethylation of cysteine residues was set as a fixed modification. Oxidation of methionine residues and acetylation of proteins n-termini were set as variable modifications. Identification results were imported into the Proline software (version 2.2.0) (Bouyssié et al., 2020) and validated. The maximum false discovery rate was set at 1% at peptide and protein levels with the use of a decoy strategy. Peptides abundances were extracted with cross assignment between all samples. Protein abundances were computed using the best ion of the unique peptide abundances normalized at the peptide level using the median. To be considered, proteins must be identified in at least three out of the four replicates in at least one condition. The imputation of the missing values and differential data analysis were performed using the open-source ProStaR software (version 1.30.7) (Wieczorek et al., 2017). Imputation of missing values was done using the approximation of the lower limit of quantification by the 2.5% lower quantile of each replicate intensity distribution (“det quantile”). A Limma moderated t-test was applied on the dataset to perform differential analysis. The adaptive Benjamini-Hochberg procedure was applied to adjust the p-values and False Discovery Rate. The mass spectrometry proteomics data have been deposited to the ProteomeXchange Consortium via the PRIDE partner repository (Perez-Riverol et al., 2022) with the dataset identifier PXD042007.

### Transcriptomic analysis

#### Patient-derived cell lines

WM1862, WM983A, WM983B. RNA integrity was assessed by Bioanalyzer (total RNA Pico Kit, 2100 Instrument, Agilent Technologies, Paolo Alto, CA, USA). All samples had RNA integrity numbers above 9.5. Sequencing libraries were prepared using “NEBNext Ultra II Directional RNA Library Prep Kit for Illumina” combined with “NEB Ultra II polyA mRNA magnetic isolation” for mRNA enrichment (New England Biolabs, Ipswich, MA, USA). Libraries were pooled and sequenced (single-end, 100bp) on a NextSeq2000 according to the manufacturer’s instructions (Illumina Inc., San Diego, CA, USA). For each sample, quality control was carried out and assessed with the NGS Core Tool FastQC (Andrews S, 2010). Sequence reads (minimum 33 Million per sample) were mapped to Homo Sapiens genome version GRCh38 using STAR (Dobin et al., 2013) to obtain a BAM (Binary Alignment Map) file. An abundance matrix was generated based on read counts identified by Featurecounts (Liao et al., 2014) using default parameters. At last, differential expression analyses were performed using the DEseq2 (Love et al., 2014) package of the Bioconductor framework for RNASeq data (Gentleman et al., 2004). Up- and down-regulated genes were selected based on their adjusted p-value (< 0.01). Functional enrichment analyses were performed using STRING v11 (Szklarczyk et al., 2019) and Gene Ontology (Carbon et a1., 2021). Bubble plots and heat maps (Fig 1g) were generated using GraphPad Prism 9 (version 9.5.1 for Windows). Raw data (FASTQ files) are available at the EMBL-EBI ArrayExpress archive (Accession number E-MTAB-13165).

### Statistical analysis

Statistical analysis of the results was done using GraphPad Prism 9 (version 9.5.1 for Windows). Mann-Whitney (two groups) or Kruskal-Wallis (>2 groups) with Dunn’s multiple comparison *post-hoc* test statistical tests with Bonferroni multiple comparison correction were performed as specified in the figure legends, to the exception of quantitative proteomics and transcriptomic analysis which have dedicated statistical methodologies specified above. Illustrations of the statistical analyses are displayed in the figures as the mean +/- standard deviation (SD). p-Values smaller than 0.05 were considered as statistically significant. * p<0.05; ** p<0.01; *** p<0.001; **** p<0.0001.

## References

Akbani, R., Akdemir, K.C., Aksoy, B.A., Albert, M., Ally, A., Amin, S.B., Arachchi, H., Arora, A., Auman, J.T., Ayala, B., Baboud, J., Balasundaram, M., Balu, S., Barnabas, N., Bartlett, J., Bartlett, P., Bastian, B.C., Baylin, S.B., Behera, M., Belyaev, D., Benz, C., Bernard, B., Beroukhim, R., Bir, N., Black, A.D., Bodenheimer, T., Boice, L., Boland, G.M., Bono, R., Bootwalla, M.S., Bosenberg, M., Bowen, J., Bowlby, R., Bristow, C.A., Brockway-Lunardi, L., Brooks, D., Brzezinski, J., Bshara, W., Buda, E., Burns, W.R., Butterfield, Y.S.N., Button, M., Calderone, T., Cappellini, G.A., Carter, C., Carter, S.L., Cherney, L., Cherniack, A.D., Chevalier, A., Chin, L., Cho, J., Cho, R.J., Choi, Y.-L., Chu, A., Chudamani, S., Cibulskis, K., Ciriello, G., Clarke, A., Coons, S., Cope, L., Crain, D., Curley, E., Danilova, L., D’Atri, S., Davidsen, T., Davies, M.A., Delman, K.A., Demchok, J.A., Deng, Q.A., Deribe, Y.L., Dhalla, N., Dhir, R., DiCara, D., Dinikin, M., Dubina, M., Ebrom, J.S., Egea, S., Eley, G., Engel, J., Eschbacher, J.M., Fedosenko, K.V., Felau, I., Fennell, T., Ferguson, M.L., Fisher, S., Flaherty, K.T., Frazer, S., Frick, J., Fulidou, V., Gabriel, S.B., Gao, J., Gardner, J., Garraway, L.A., Gastier-Foster, J.M., Gaudioso, C., Gehlenborg, N., Genovese, G., Gerken, M., Gershenwald, J.E., Getz, G., Gomez-Fernandez, C., Gribbin, T., Grimsby, J., Gross, B., Guin, R., Gutschner, T., Hadjipanayis, A., Halaban, R., Hanf, B., Haussler, D., Haydu, L.E., Hayes, D.N., Hayward, N.K., Heiman, D.I., Herbert, L., Herman, J.G., Hersey, P., Hoadley, K.A., Hodis, E., Holt, R.A., Hoon, D.S., Hoppough, S., Hoyle, A.P., Huang, F.W., Huang, M., Huang, S., Hutter, C.M., Ibbs, M., Iype, L., Jacobsen, A., Jakrot, V., Janning, A., Jeck, W.R., Jefferys, S.R., Jensen, M.A., Jones, C.D., Jones, S.J.M., Ju, Z., Kakavand, H., Kang, H., Kefford, R.F., Khuri, F.R., Kim, J., Kirkwood, J.M., Klode, J., Korkut, A., Korski, K., Krauthammer, M., Kucherlapati, R., Kwong, L.N., Kycler, W., Ladanyi, M., Lai, P.H., Laird, P.W., Lander, E., Lawrence, M.S., Lazar, A.J., Łaźniak, R., Lee, D., Lee, J.E., Lee, J., Lee, K., Lee, S., Lee, W., Leporowska, E., Leraas, K.M., Li, H.I., Lichtenberg, T.M., Lichtenstein, L., Lin, P., Ling, S., Liu, J., Liu, O., Liu, W., Long, G.V., Lu, Y., Ma, S., Ma, Y., Mackiewicz, A., Mahadeshwar, H.S., Malke, J., Mallery, D., Manikhas, G.M., Mann, G.J., Marra, M.A., Matejka, B., Mayo, M., Mehrabi, S., Meng, S., Meyerson, M., Mieczkowski, P.A., Miller, J.P., Miller, M.L., Mills, G.B., Moiseenko, F., Moore, R.A., Morris, S., Morrison, C., Morton, D., Moschos, S., Mose, L.E., Muller, F.L., Mungall, A.J., Murawa, D., Murawa, P., Murray, B.A., Nezi, L., Ng, S., Nicholson, D., Noble, M.S., Osunkoya, A., Owonikoko, T.K., Ozenberger, B.A., Pagani, E., Paklina, O.V., Pantazi, A., Parfenov, M., Parfitt, J., Park, P.J., Park, W.-Y., Parker, J.S., Passarelli, F., Penny, R., Perou, C.M., Pihl, T.D., Potapova, O., Prieto, V.G., Protopopov, A., Quinn, M.J., Radenbaugh, A., Rai, K., Ramalingam, S.S., Raman, A.T., Ramirez, N.C., Ramirez, R., Rao, U., Rathmell, W.K., Ren, X., Reynolds, S.M., Roach, J., Robertson, A.G., Ross, M.I., Roszik, J., Russo, G., Saksena, G., Saller, C., Samuels, Y., Sander, Chris, Sander, Cindy, Sandusky, G., Santoso, N., Saul, M., Saw, R.P., Schadendorf, D., Schein, J.E., Schultz, N., Schumacher, S.E., Schwallier, C., Scolyer, R.A., Seidman, J., Sekhar, P.C., Sekhon, H.S., Senbabaoglu, Y., Seth, S., Shannon, K.F., Sharpe, S., Sharpless, N.E., Shaw, K.R.M., Shelton, C., Shelton, T., Shen, R., Sheth, M., Shi, Y., Shiau, C.J., Shmulevich, I., Sica, G.L., Simons, J.V., Sinha, R., Sipahimalani, P., Sofia, H.J., Soloway, M.G., Song, X., Sougnez, C., Spillane, A.J., Spychała, A., Stretch, J.R., Stuart, J., Suchorska, W.M., Sucker, A., Sumer, S.O., Sun, Y., Synott, M., Tabak, B., Tabler, T.R., Tam, A., Tan, D., Tang, J., Tarnuzzer, R., Tarvin, K., Tatka, H., Taylor, B.S., Teresiak, M., Thiessen, N., Thompson, J.F., Thorne, L., Thorsson, V., Trent, J.M., Triche, T.J., Tsai, K.Y., Tsou, P., Van Den Berg, D.J., Van Allen, E.M., Veluvolu, U., Verhaak, R.G., Voet, D., Voronina, O., Walter, V., Walton, J.S., Wan, Y., Wang, Y., Wang, Z., Waring, S., Watson, I.R., Weinhold, N., Weinstein, J.N., Weisenberger, D.J., White, P., Wilkerson, M.D., Wilmott, J.S., Wise, L., Wiznerowicz, M., Woodman, S.E., Wu, C.-J., Wu, C.-C., Wu, J., Wu, Y., Xi, R., Xu, A.W., Yang, D., Yang, Liming, Yang, Lixing, Zack, T.I., Zenklusen, J.C., Zhang, H., Zhang, J., Zhang, W., Zhao, X., Zhu, J., Zhu, K., Zimmer, L., Zmuda, E., Zou, L., 2015. Genomic Classification of Cutaneous Melanoma. Cell 161, 1681–1696. 10.1016/j.cell.2015.05.044

Andrews S, 2010. FastQC: a quality control tool for high throughput sequence data.

Ballabio, A., Bonifacino, J.S., 2020. Lysosomes as dynamic regulators of cell and organismal homeostasis. Nat. Rev. Mol. Cell Biol. 21, 101–118. 10.1038/s41580-019-0185-4

Bayle, J.H., Grimley, J.S., Stankunas, K., Gestwicki, J.E., Wandless, T.J., Crabtree, G.R., 2006. Rapamycin Analogs with Differential Binding Specificity Permit Orthogonal Control of Protein Activity. Chem. Biol. 13, 99–107. 10.1016/j.chembiol.2005.10.017

Bian, B., Mongrain, S., Cagnol, S., Langlois, M.-J., Boulanger, J., Bernatchez, G., Carrier, J.C., Boudreau, F., Rivard, N., 2016. Cathepsin B promotes colorectal tumorigenesis, cell invasion, and metastasis. Mol. Carcinog. 55, 671–687. 10.1002/mc.22312

Bouyssié, D., Hesse, A.-M., Mouton-Barbosa, E., Rompais, M., Macron, C., Carapito, C., Gonzalez de Peredo, A., Couté, Y., Dupierris, V., Burel, A., Menetrey, J.-P., Kalaitzakis, A., Poisat, J., Romdhani, A., Burlet-Schiltz, O., Cianférani, S., Garin, J., Bruley, C., 2020. Proline: an efficient and user-friendly software suite for large-scale proteomics. Bioinformatics 36, 3148–3155. 10.1093/bioinformatics/btaa118

Braeuer, R.R., Zigler, M., Villares, G.J., Dobroff, A.S., Bar-Eli, M., 2011. Transcriptional Control of Melanoma Metastasis: The Importance of the Tumor Microenvironment. Semin. Cancer Biol. 21, 83–88. 10.1016/j.semcancer.2010.12.007

Chaineau, M., Danglot, L., Galli, T., 2009. Multiple roles of the vesicular-SNARE TI-VAMP in post-Golgi and endosomal trafficking. FEBS Lett. 583, 3817–3826. 10.1016/j.febslet.2009.10.026

Chen, C.-S., Chen, W.-N.U., Zhou, M., Arttamangkul, S., Haugland, R.P., 2000. Probing the cathepsin D using a BODIPY FL–pepstatin A: applications in fluorescence polarization and microscopy. J. Biochem. Biophys. Methods 42, 137–151. 10.1016/S0165-022X(00)00048-8

Circu, M.L., Dykes, S.S., Carroll, J., Kelly, K., Galiano, F., Greer, A., Cardelli, J., El-Osta, H., 2016. A Novel High Content Imaging-Based Screen Identifies the Anti-Helminthic Niclosamide as an Inhibitor of Lysosome Anterograde Trafficking and Prostate Cancer Cell Invasion. PLOS ONE 11, e0146931. 10.1371/journal.pone.0146931

Corrotte, M., Castro-Gomes, T., 2019. Chapter One - Lysosomes and plasma membrane repair, in: Andrade, L.O. (Ed.), Current Topics in Membranes, Plasma Membrane Repair. Academic Press, pp. 1–16. 10.1016/bs.ctm.2019.08.001

da Costa, A.C., Santa-Cruz, F., Mattos, L.A.R., Rêgo Aquino, M.A., Martins, C.R., Bandeira Ferraz, Á.A., Figueiredo, J.L., 2020. Cathepsin S as a target in gastric cancer. Mol. Clin. Oncol. 12, 99–103. 10.3892/mco.2019.1958

Dillekås, H., Rogers, M.S., Straume, O., 2019. Are 90% of deaths from cancer caused by metastases? Cancer Med. 8, 5574–5576. 10.1002/cam4.2474

Dobin, A., Davis, C.A., Schlesinger, F., Drenkow, J., Zaleski, C., Jha, S., Batut, P., Chaisson, M., Gingeras, T.R., 2013. STAR: ultrafast universal RNA-seq aligner. Bioinformatics 29, 15–21. 10.1093/bioinformatics/bts635

Duong, T., Goud, B., Schauer, K., 2012. Closed-form density-based framework for automatic detection of cellular morphology changes. Proc. Natl. Acad. Sci. 109, 8382–8387. 10.1073/pnas.1117796109

Dykes, S.S., Gray, A.L., Coleman, D.T., Saxena, M., Stephens, C.A., Carroll, J.L., Pruitt, K., Cardelli, J.A., 2016. The Arf-like GTPase Arl8b is essential for three-dimensional invasive growth of prostate cancer in vitro and xenograft formation and growth in vivo. Oncotarget 7, 31037–31052. 10.18632/oncotarget.8832

Follain, G., Osmani, N., Azevedo, A.S., Allio, G., Mercier, L., Karreman, M.A., Solecki, G., Garcia Leòn, M.J., Lefebvre, O., Fekonja, N., Hille, C., Chabannes, V., Dollé, G., Metivet, T., Hovsepian, F.D., Prudhomme, C., Pichot, A., Paul, N., Carapito, R., Bahram, S., Ruthensteiner, B., Kemmling, A., Siemonsen, S., Schneider, T., Fiehler, J., Glatzel, M., Winkler, F., Schwab, Y., Pantel, K., Harlepp, S., Goetz, J.G., 2018. Hemodynamic Forces Tune the Arrest, Adhesion, and Extravasation of Circulating Tumor Cells. Dev. Cell 45, 33–52.e12. 10.1016/j.devcel.2018.02.015

Fourriere, L., Kasri, A., Gareil, N., Bardin, S., Bousquet, H., Pereira, D., Perez, F., Goud, B., Boncompain, G., Miserey-Lenkei, S., 2019. RAB6 and microtubules restrict protein secretion to focal adhesions. J. Cell Biol. 218, 2215–2231. 10.1083/jcb.201805002

Gentleman, R.C., Carey, V.J., Bates, D.M., Bolstad, B., Dettling, M., Dudoit, S., Ellis, B., Gautier, L., Ge, Y., Gentry, J., Hornik, K., Hothorn, T., Huber, W., Iacus, S., Irizarry, R., Leisch, F., Li, C., Maechler, M., Rossini, A.J., Sawitzki, G., Smith, C., Smyth, G., Tierney, L., Yang, J.Y., Zhang, J., 2004. Bioconductor: open software development for computational biology and bioinformatics. Genome Biol. 5, R80. 10.1186/gb-2004-5-10-r80

Gocheva, V., Zeng, W., Ke, D., Klimstra, D., Reinheckel, T., Peters, C., Hanahan, D., Joyce, J.A., 2006. Distinct roles for cysteine cathepsin genes in multistage tumorigenesis. Genes Dev. 20, 543–556. 10.1101/gad.1407406

Goldman, M.J., Craft, B., Hastie, M., Repečka, K., McDade, F., Kamath, A., Banerjee, A., Luo, Y., Rogers, D., Brooks, A.N., Zhu, J., Haussler, D., 2020. Visualizing and interpreting cancer genomics data via the Xena platform. Nat. Biotechnol. 38, 675–678. 10.1038/s41587-020-0546-8

Gutierrez-Ruiz, O.L., Johnson, K.M., Krueger, E.W., Nooren, R.E., Cruz-Reyes, N., Heppelmann, C.J., Hogenson, T.L., Fernandez-Zapico, M.E., McNiven, M.A., Razidlo, G.L., 2023. Ectopic expression of DOCK8 regulates lysosome-mediated pancreatic tumor cell invasion. Cell Rep. 42, 113042. 10.1016/j.celrep.2023.113042

Hämälistö, S., Jäättelä, M., 2016. Lysosomes in cancer-living on the edge (of the cell). Curr. Opin. Cell Biol. 39, 69–76. 10.1016/j.ceb.2016.02.009

Heinrich, L., Bennett, D., Ackerman, D., Park, W., Bogovic, J., Eckstein, N., Petruncio, A., Clements, J., Pang, S., Xu, C.S., Funke, J., Korff, W., Hess, H.F., Lippincott-Schwartz, J., Saalfeld, S., Weigel, A.V., 2021. Whole-cell organelle segmentation in volume electron microscopy. Nature 599, 141–146. 10.1038/s41586-021-03977-3

Hooikaas, P.J., Martin, M., Mühlethaler, T., Kuijntjes, G.-J., Peeters, C.A.E., Katrukha, E.A., Ferrari, L., Stucchi, R., Verhagen, D.G.F., van Riel, W.E., Grigoriev, I., Altelaar, A.F.M., Hoogenraad, C.C., Rüdiger, S.G.D., Steinmetz, M.O., Kapitein, L.C., Akhmanova, A., 2019. MAP7 family proteins regulate kinesin-1 recruitment and activation. J. Cell Biol. 218, 1298–1318. 10.1083/jcb.201808065

Hoshino, D., Kirkbride, K.C., Costello, K., Clark, E.S., Sinha, S., Grega-Larson, N., Tyska, M.J., Weaver, A.M., 2013. Exosome secretion is enhanced by invadopodia and drives invasive behavior. Cell Rep. 5, 10.1016/j.celrep.2013.10.050. 10.1016/j.celrep.2013.10.050

Hua, H., Li, M., Luo, T., Yin, Y., Jiang, Y., 2011. Matrix metalloproteinases in tumorigenesis: an evolving paradigm. Cell. Mol. Life Sci. CMLS 68, 3853–3868. 10.1007/s00018-011-0763-x

Hughes, C.S., Moggridge, S., Müller, T., Sorensen, P.H., Morin, G.B., Krijgsveld, J., 2019. Single-pot, solid-phase-enhanced sample preparation for proteomics experiments. Nat. Protoc. 14, 68–85. 10.1038/s41596-018-0082-x

Jacob, A., Prekeris, R., 2015. The regulation of MMP targeting to invadopodia during cancer metastasis. Front. Cell Dev. Biol. 3, 4. 10.3389/fcell.2015.00004

Jerabkova-Roda, K., Marwaha, R., Das, T., Goetz, J.G., 2024. Organelle morphology and positioning orchestrate physiological and disease-associated processes. Curr. Opin. Cell Biol. 86, 102293. 10.1016/j.ceb.2023.102293

Jia, R., Bonifacino, J.S., 2019. Lysosome Positioning Influences mTORC2 and AKT Signaling. Mol. Cell 75, 26–38.e3. 10.1016/j.molcel.2019.05.009

Johnson, D.E., Ostrowski, P., Jaumouillé, V., Grinstein, S., 2016. The position of lysosomes within the cell determines their luminal pH. J. Cell Biol. 212, 677–692. 10.1083/jcb.201507112

Kalluri, R., 2016. The biology and function of fibroblasts in cancer. Nat. Rev. Cancer 16, 582–598. 10.1038/nrc.2016.73

Kapitein, L.C., Schlager, M.A., van der Zwan, W.A., Wulf, P.S., Keijzer, N., Hoogenraad, C.C., 2010. Probing Intracellular Motor Protein Activity Using an Inducible Cargo Trafficking Assay. Biophys. J. 99, 2143–2152. 10.1016/j.bpj.2010.07.055

Kolli-Bouhafs, K., Sick, E., Noulet, F., Gies, J.-P., De Mey, J., Rondé, P., 2014. FAK competes for Src to promote migration against invasion in melanoma cells. Cell Death Dis. 5, e1379. 10.1038/cddis.2014.329

Korolchuk, V.I., Saiki, S., Lichtenberg, M., Siddiqi, F.H., Roberts, E.A., Imarisio, S., Jahreiss, L., Sarkar, S., Futter, M., Menzies, F.M., O’Kane, C.J., Deretic, V., Rubinsztein, D.C., 2011. Lysosomal positioning coordinates cellular nutrient responses. Nat. Cell Biol. 13, 453–460. 10.1038/ncb2204

Kundu, S.T., Grzeskowiak, C.L., Fradette, J.J., Gibson, L.A., Rodriguez, L.B., Creighton, C.J., Scott, K.L., Gibbons, D.L., 2018. TMEM106B drives lung cancer metastasis by inducing TFEB-dependent lysosome synthesis and secretion of cathepsins. Nat. Commun. 9, 2731. 10.1038/s41467-018-05013-x

Lachuer, H., Le, L., Lévêque-Fort, S., Goud, B., Schauer, K., 2023. Spatial organization of lysosomal exocytosis relies on membrane tension gradients. Proc. Natl. Acad. Sci. 120, e2207425120. 10.1073/pnas.2207425120

Lawson, N.D., Weinstein, B.M., 2002. In vivo imaging of embryonic vascular development using transgenic zebrafish. Dev. Biol. 248, 307–318. 10.1006/dbio.2002.0711

Liao, Y., Smyth, G.K., Shi, W., 2014. featureCounts: an efficient general purpose program for assigning sequence reads to genomic features. Bioinformatics 30, 923–930. 10.1093/bioinformatics/btt656

Love, M.I., Huber, W., Anders, S., 2014. Moderated estimation of fold change and dispersion for RNA-seq data with DESeq2. Genome Biol. 15, 550. 10.1186/s13059-014-0550-8

Machado, E., White-Gilbertson, S., van de Vlekkert, D., Janke, L., Moshiach, S., Campos, Y., Finkelstein, D., Gomero, E., Mosca, R., Qiu, X., Morton, C.L., Annunziata, I., d’Azzo, A., 2015. Regulated lysosomal exocytosis mediates cancer progression. Sci. Adv. 1, e1500603. 10.1126/sciadv.1500603

Marwaha, R., Rawal, S., Khuntia, P., Banerjee, S., Manoj, D., Jaiswal, M., Das, T., 2023. Mechanosensitive dynamics of lysosomes along microtubules regulate leader cell emergence in collective cell migration. 10.1101/2022.08.03.502740

Mathur, P., De Barros Santos, C., Lachuer, H., Patat, J., Latgé, B., Radvanyi, F., Goud, B., Schauer, K., 2023. Transcription factor EB regulates phosphatidylinositol-3-phosphate levels that control lysosome positioning in the bladder cancer model. Commun. Biol. 6, 1–14. 10.1038/s42003-023-04501-1

Medina, D.L., Fraldi, A., Bouche, V., Annunziata, F., Mansueto, G., Spampanato, C., Puri, C., Pignata, A., Martina, J.A., Sardiello, M., Palmieri, M., Polishchuk, R., Puertollano, R., Ballabio, A., 2011. Transcriptional Activation of Lysosomal Exocytosis Promotes Cellular Clearance. Dev. Cell 21, 421–430. 10.1016/j.devcel.2011.07.016

Moamer, A., Hachim, I.Y., Binothman, N., Wang, N., Lebrun, J.-J., Ali, S., 2019. A role for kinesin-1 subunits KIF5B/KLC1 in regulating epithelial mesenchymal plasticity in breast tumorigenesis. EBioMedicine 45, 92–107. 10.1016/j.ebiom.2019.06.009

Monteiro, P., Rossé, C., Castro-Castro, A., Irondelle, M., Lagoutte, E., Paul-Gilloteaux, P., Desnos, C., Formstecher, E., Darchen, F., Perrais, D., Gautreau, A., Hertzog, M., Chavrier, P., 2013. Endosomal WASH and exocyst complexes control exocytosis of MT1-MMP at invadopodia. J. Cell Biol. 203, 1063–1079. 10.1083/jcb.201306162

Mulligan, R.J., Magaj, M.M., Digilio, L., Redemann, S., Yap, C.C., Winckler, B., 2024. Collapse of late endosomal pH elicits a rapid Rab7 response via the V-ATPase and RILP. J. Cell Sci. 137, jcs261765. 10.1242/jcs.261765

Naegeli, K.M., Hastie, E., Garde, A., Wang, Z., Keeley, D.P., Gordon, K.L., Pani, A.M., Kelley, L.C., Morrissey, M.A., Chi, Q., Goldstein, B., Sherwood, D.R., 2017. Cell invasion in vivo via rapid exocytosis of a transient lysosome-derived membrane domain. Dev. Cell 43, 403–417.e10. 10.1016/j.devcel.2017.10.024

Nair, S.V., Narendradev, N.D., Nambiar, R.P., Kumar, R., Srinivasula, S.M., 2020. Naturally occurring and tumor-associated variants of RNF167 promote lysosomal exocytosis and plasma membrane resealing. J. Cell Sci. 133, jcs239335. 10.1242/jcs.239335

Nakahara, H., Howard, L., Thompson, E.W., Sato, H., Seiki, M., Yeh, Y., Chen, W.-T., 1997. Transmembrane/cytoplasmic domain-mediated membrane type 1-matrix metalloprotease docking to invadopodia is required for cell invasion. Proc. Natl. Acad. Sci. 94, 7959–7964. 10.1073/pnas.94.15.7959

Perez-Riverol, Y., Bai, J., Bandla, C., García-Seisdedos, D., Hewapathirana, S., Kamatchinathan, S., Kundu, D.J., Prakash, A., Frericks-Zipper, A., Eisenacher, M., Walzer, M., Wang, S., Brazma, A., Vizcaíno, J.A., 2022. The PRIDE database resources in 2022: a hub for mass spectrometry-based proteomics evidences. Nucleic Acids Res. 50, D543–D552. 10.1093/nar/gkab1038

Pu, J., Guardia, C.M., Keren-Kaplan, T., Bonifacino, J.S., 2016a. Mechanisms and functions of lysosome positioning. J. Cell Sci. 129, 4329–4339. 10.1242/jcs.196287

Pu, J., Guardia, C.M., Keren-Kaplan, T., Bonifacino, J.S., 2016b. Mechanisms and functions of lysosome positioning. J. Cell Sci. 129, 4329–4339. 10.1242/jcs.196287

Sadok, A., McCarthy, A., Caldwell, J., Collins, I., Garrett, M.D., Yeo, M., Hooper, S., Sahai, E., Kuemper, S., Mardakheh, F.K., Marshall, C.J., 2015. Rho Kinase Inhibitors Block Melanoma Cell Migration and Inhibit Metastasis. Cancer Res. 75, 2272–2284. 10.1158/0008-5472.CAN-14-2156

Sahai, E., Astsaturov, I., Cukierman, E., DeNardo, D.G., Egeblad, M., Evans, R.M., Fearon, D., Greten, F.R., Hingorani, S.R., Hunter, T., Hynes, R.O., Jain, R.K., Janowitz, T., Jorgensen, C., Kimmelman, A.C., Kolonin, M.G., Maki, R.G., Powers, R.S., Puré, E., Ramirez, D.C., Scherz-Shouval, R., Sherman, M.H., Stewart, S., Tlsty, T.D., Tuveson, D.A., Watt, F.M., Weaver, V., Weeraratna, A.T., Werb, Z., 2020. A framework for advancing our understanding of cancer-associated fibroblasts. Nat. Rev. Cancer 20, 174–186. 10.1038/s41568-019-0238-1

Schauer, K., Duong, T., Bleakley, K., Bardin, S., Bornens, M., Goud, B., 2010. Probabilistic density maps to study global endomembrane organization. Nat. Methods 7, 560–566. 10.1038/nmeth.1462

Schindelin, J., Arganda-Carreras, I., Frise, E., Kaynig, V., Longair, M., Pietzsch, T., Preibisch, S., Rueden, C., Saalfeld, S., Schmid, B., Tinevez, J.-Y., White, D.J., Hartenstein, V., Eliceiri, K., Tomancak, P., Cardona, A., 2012. Fiji: an open-source platform for biological-image analysis. Nat. Methods 9, 676–682. 10.1038/nmeth.2019

Sero, J.E., Sailem, H.Z., Ardy, R.C., Almuttaqi, H., Zhang, T., Bakal, C., 2015. Cell shape and the microenvironment regulate nuclear translocation of NF-κB in breast epithelial and tumor cells. Mol. Syst. Biol. 11, 790. 10.15252/msb.20145644

Serra-Marques, A., Martin, M., Katrukha, E.A., Grigoriev, I., Peeters, C.A., Liu, Q., Hooikaas, P.J., Yao, Y., Solianova, V., Smal, I., Pedersen, L.B., Meijering, E., Kapitein, L.C., Akhmanova, A., 2020. Concerted action of kinesins KIF5B and KIF13B promotes efficient secretory vesicle transport to microtubule plus ends. eLife 9, e61302. 10.7554/eLife.61302

Steffan, J.J., Snider, J.L., Skalli, O., Welbourne, T., Cardelli, J.A., 2009. Na+/H+ exchangers and RhoA regulate acidic extracellular pH-induced lysosome trafficking in prostate cancer cells. Traffic Cph. Den. 10, 737–753. 10.1111/j.1600-0854.2009.00904.x

Stirling, D.R., Swain-Bowden, M.J., Lucas, A.M., Carpenter, A.E., Cimini, B.A., Goodman, A., 2021. CellProfiler 4: improvements in speed, utility and usability. BMC Bioinformatics 22, 433. 10.1186/s12859-021-04344-9

Stoletov, K., Kato, H., Zardouzian, E., Kelber, J., Yang, J., Shattil, S., Klemke, R., 2010. Visualizing extravasation dynamics of metastatic tumor cells. J. Cell Sci. 123, 2332–2341. 10.1242/jcs.069443

van Bergeijk, P., Hoogenraad, C.C., Kapitein, L.C., 2016. Right Time, Right Place: Probing the Functions of Organelle Positioning. Trends Cell Biol. 26, 121–134. 10.1016/j.tcb.2015.10.001

Vidak, E., Javoršek, U., Vizovišek, M., Turk, B., 2019. Cysteine Cathepsins and Their Extracellular Roles: Shaping the Microenvironment. Cells 8, 264. 10.3390/cells8030264

Wang, L., Goldwag, J., Bouyea, M., Barra, J., Matteson, K., Maharjan, N., Eladdadi, A., Embrechts, M.J., Intes, X., Kruger, U., Barroso, M., 2023. Spatial topology of organelle is a new breast cancer cell classifier. iScience 107229. 10.1016/j.isci.2023.107229

Wang, Q., Yao, J., Jin, Q., Wang, X., Zhu, H., Huang, F., Wang, W., Qiang, J., Ni, Q., 2017. LAMP1 expression is associated with poor prognosis in breast cancer. Oncol. Lett. 14, 4729–4735. 10.3892/ol.2017.6757

Wang, Y., McNiven, M.A., 2012. Invasive matrix degradation at focal adhesions occurs via protease recruitment by a FAK–p130Cas complex. J. Cell Biol. 196, 375–385. 10.1083/jcb.201105153

Wieczorek, S., Combes, F., Lazar, C., Giai Gianetto, Q., Gatto, L., Dorffer, A., Hesse, A.-M., Couté, Y., Ferro, M., Bruley, C., Burger, T., 2017. DAPAR & ProStaR: software to perform statistical analyses in quantitative discovery proteomics. Bioinformatics 33, 135–136. 10.1093/bioinformatics/btw580

Winkler, J., Abisoye-Ogunniyan, A., Metcalf, K.J., Werb, Z., 2020. Concepts of extracellular matrix remodelling in tumour progression and metastasis. Nat. Commun. 11, 5120. 10.1038/s41467-020-18794-x

Wu, Pei-Hsun, Gilkes, D.M., Phillip, J.M., Narkar, A., Cheng, T.W.-T., Marchand, J., Lee, M.-H., Li, R., Wirtz, D., 2020. Single-cell morphology encodes metastatic potential. Sci. Adv. 6, eaaw6938. 10.1126/sciadv.aaw6938

Wu, Ping-Hsiu, Onodera, Y., Giaccia, A.J., Le, Q.-T., Shimizu, S., Shirato, H., Nam, J.-M., 2020. Lysosomal trafficking mediated by Arl8b and BORC promotes invasion of cancer cells that survive radiation. Commun. Biol. 3, 620. 10.1038/s42003-020-01339-9

