## Supplementary material for "Peripheral positioning of lysosomes supports melanoma aggressiveness": Table 1

### Table 1: Lysosome genes differentially expressed (refers to Figure 1A)

| Gene | GeneSymbol | Average | WM1862 | WM1862 | WM1862 | WM1862 | WM983A | WM983A | WM983A | WM983A | WM983B | WM983B | WM983B | WM983B |
|---|---|---|---|---|---|---|---|---|---|---|---|---|---|---|
| ENSG00000101986 | ABCD1 | 2606.597931 | 1.70984909 | 1.698634074 | 2.165143828 | 1.865820678 | 0.470239199 | 0.619388744 | 0.661886886 | 0.412210243 | 0.621293358 | 0.676469725 | 0.561573056 | 0.537491118 |
| ENSG00000157766 | ACAN | 79.82246872 | 1.362753019 | 0.851552102 | 2.71846108 | 1.807085612 | 0.540691298 | 1.069125149 | 1.166533055 | 1.793663063 | 0.132210469 | 0.20820055 | 0.160860829 | 0.188863776 |
| ENSG00000102575 | ACP5 | 1927.866114 | 2.197409265 | 2.155628998 | 2.285839264 | 1.642360738 | 0.555199858 | 0.545654435 | 0.645212426 | 0.393813401 | 0.300081179 | 0.435763968 | 0.276881243 | 0.566155225 |
| ENSG00000196839 | ADA | 450.2645654 | 1.478605243 | 1.284341113 | 1.37148154 | 1.145110869 | 0.832553793 | 0.928327236 | 0.659424789 | 0.942267634 | 0.705275078 | 0.943040102 | 0.794409187 | 0.915163416 |
| ENSG00000038002 | AGA | 478.6861845 | 1.731679416 | 1.350083426 | 1.486486167 | 1.510961127 | 0.499755206 | 0.754961541 | 0.86617859 | 0.532341542 | 0.825742499 | 0.725608597 | 0.823881942 | 0.892319946 |
| ENSG00000151150 | ANK3 | 60.30011197 | 1.102413111 | 1.370036056 | 1.859262028 | 2.069790946 | 1.083837671 | 0.563214871 | 0.39333445 | 2.613982342 | 0.143193302 | 0.316947075 | 0.167310034 | 0.316678112 |
| ENSG00000106367 | AP1S1 | 2971.060627 | 1.672654271 | 1.380444973 | 1.664611327 | 1.320844996 | 0.543293643 | 0.717216183 | 0.702212312 | 0.349707134 | 0.886721693 | 0.940013368 | 0.868680356 | 0.953599745 |
| ENSG00000182287 | AP1S2 | 4022.719991 | 1.615698843 | 1.466943107 | 1.43553619 | 1.324196832 | 0.696764672 | 0.706574519 | 0.72062691 | 0.587749439 | 0.837592959 | 0.829979107 | 0.821698322 | 0.956639099 |
| ENSG00000006125 | AP2B1 | 4857.403923 | 1.426394459 | 1.098181513 | 1.371271607 | 1.214592242 | 0.710059493 | 0.792043652 | 0.899715506 | 0.775828718 | 0.859178404 | 1.021285447 | 0.925396611 | 0.906052348 |
| ENSG00000053770 | AP5M1 | 1034.824471 | 1.440501045 | 1.15303364 | 1.432894418 | 1.481902782 | 0.76025709 | 0.80616965 | 0.921890786 | 0.780634871 | 0.772283268 | 0.76605282 | 0.818054866 | 0.866324763 |
| ENSG00000068650 | ATP11A | 3358.66902 | 1.73512103 | 1.124925541 | 1.580992558 | 1.694441667 | 0.844436736 | 0.605666085 | 0.483338175 | 1.212366723 | 0.696980989 | 0.743454337 | 0.597212647 | 0.681063512 |
| ENSG00000033627 | ATP6V0A1 | 3201.369895 | 1.086373778 | 1.079262795 | 0.948096444 | 1.025779337 | 1.402462679 | 0.947153864 | 0.703830127 | 1.826254335 | 0.767788465 | 0.7158721 | 0.652627372 | 0.844498704 |
| ENSG00000147614 | ATP6V0D2 | 513.3890894 | 2.607335387 | 2.446354534 | 2.182034962 | 2.259702107 | 0.369896802 | 0.522434329 | 0.586899615 | 0.319817998 | 0.175662692 | 0.187753649 | 0.107189449 | 0.234918476 |
| ENSG00000100554 | ATP6V1D | 1811.678397 | 1.248668624 | 1.337997648 | 1.262631003 | 1.194866344 | 0.95563598 | 0.719562165 | 0.839329141 | 0.780136662 | 0.890724094 | 0.889810765 | 0.891508877 | 0.989128697 |
| ENSG00000186318 | BACE1 | 282.044706 | 1.689124811 | 1.145680579 | 1.557956581 | 1.432728546 | 0.786975194 | 0.725567022 | 1.308120277 | 0.419144797 | 0.734740726 | 0.945723349 | 0.744671875 | 0.509566243 |
| ENSG00000105327 | BBC3 | 867.9969728 | 1.745211952 | 1.838481852 | 1.456216656 | 1.388391195 | 1.324049411 | 0.615995849 | 0.384575599 | 1.607110832 | 0.553755216 | 0.296770354 | 0.352919518 | 0.436521565 |
| ENSG00000132692 | BCAN | 700.9108321 | 2.738915775 | 1.960449311 | 2.554109143 | 2.152851229 | 0.418717079 | 0.365268469 | 0.448679879 | 0.260490297 | 0.265544477 | 0.323650965 | 0.155715416 | 0.35560796 |
| ENSG00000196072 | BLOC1S2 | 1209.945552 | 1.153761269 | 1.221235327 | 1.153768033 | 0.988401634 | 1.29534727 | 0.858622454 | 0.779748941 | 1.286447435 | 0.923758425 | 0.623587 | 0.90128771 | 0.814034503 |
| ENSG00000165714 | BORCS5 | 221.1130456 | 2.004272938 | 1.962712798 | 1.96274511 | 1.952463191 | 0.685957542 | 0.535614304 | 0.397284822 | 0.427718123 | 0.551045765 | 0.608804255 | 0.497753475 | 0.413627676 |
| ENSG00000164713 | BRI3 | 5269.460744 | 1.153149738 | 1.328044975 | 1.044417551 | 1.022349649 | 1.325918675 | 0.87454324 | 0.628313373 | 1.815447351 | 0.793449639 | 0.58945432 | 0.681068973 | 0.743842516 |
| ENSG00000185127 | C6orf120 | 597.0031538 | 2.014402511 | 1.334754428 | 1.599279337 | 1.542235355 | 0.681622323 | 0.600963666 | 0.551786387 | 0.836077418 | 0.682984141 | 0.737694042 | 0.58993209 | 0.828268303 |
| ENSG00000121691 | CAT | 1862.731881 | 1.6145742 | 1.386097456 | 1.449360726 | 1.358183779 | 0.980417832 | 0.708722921 | 0.755489623 | 1.334872507 | 0.636084663 | 0.583937452 | 0.553430876 | 0.638827964 |
| ENSG00000019582 | CD74 | 8498.83459 | 0.895114641 | 1.06138538 | 1.053843158 | 0.89677123 | 1.83397828 | 1.39790651 | 1.214492478 | 2.337319581 | 0.446914494 | 0.159271745 | 0.282957189 | 0.420045314 |
| ENSG00000089486 | CDIP1 | 857.1083793 | 1.79088533 | 1.726409126 | 1.760587561 | 1.577903665 | 0.517932604 | 0.517141828 | 0.544220779 | 0.482741166 | 0.737645994 | 0.902590943 | 0.664512693 | 0.777428311 |
| ENSG00000255112 | CHMP1B | 2530.874789 | 0.910948208 | 1.17801164 | 0.809160324 | 0.948696286 | 1.674616489 | 0.830948485 | 0.494607325 | 2.487056616 | 0.732377134 | 0.511204872 | 0.680569293 | 0.741803326 |
| ENSG00000130724 | CHMP2A | 2969.459463 | 1.141708576 | 1.372737374 | 0.993824691 | 1.03767735 | 1.716720377 | 0.882707303 | 0.707028368 | 1.427892943 | 0.709824476 | 0.583732582 | 0.647691878 | 0.77845408 |
| ENSG00000176108 | CHMP6 | 436.0028738 | 1.051091678 | 1.004957499 | 0.95597956 | 1.076981156 | 1.145439518 | 1.310213629 | 1.629954605 | 0.979114611 | 0.607320645 | 0.850007071 | 0.660522655 | 0.728417373 |
| ENSG00000073464 | CLCN4 | 265.2627459 | 1.541588015 | 1.182682084 | 1.526999118 | 1.615929959 | 1.032008333 | 1.43132367 | 1.311400257 | 1.629142821 | 0.256791357 | 0.103374807 | 0.145218 | 0.223541579 |
| ENSG00000091317 | CMTM6 | 5773.925662 | 0.94529316 | 1.108960546 | 0.943145525 | 1.160348842 | 1.692095843 | 0.8687182 | 0.472853014 | 2.148217863 | 0.807040314 | 0.45592249 | 0.646661992 | 0.75074221 |
| ENSG00000142156 | COL6A1 | 1379.402102 | 1.377112302 | 1.07954843 | 1.217844557 | 1.010857047 | 1.283719231 | 1.152756328 | 1.376832617 | 1.008427311 | 0.584929368 | 0.650593775 | 0.610378489 | 0.647000545 |
| ENSG00000143162 | CREG1 | 1558.884942 | 1.420143331 | 1.318841108 | 1.434323413 | 1.426035822 | 0.952398967 | 0.756925015 | 0.793423268 | 0.803005206 | 0.757604081 | 0.793702352 | 0.684835618 | 0.858761819 |
| ENSG00000117984 | CTSD | 12827.1489 | 1.15340186 | 1.375577914 | 1.070833252 | 0.905052894 | 1.643696639 | 1.162526089 | 0.966508204 | 1.174655903 | 0.662751798 | 0.611337238 | 0.550778759 | 0.722879449 |
| ENSG00000103811 | CTSH | 1595.141442 | 2.809825604 | 2.39679016 | 2.482629695 | 2.342800159 | 0.398120482 | 0.286607619 | 0.268192175 | 0.437255061 | 0.178029031 | 0.104185632 | 0.137418791 | 0.158145591 |
| ENSG00000143387 | CTSK | 4572.732114 | 1.949773656 | 1.483969717 | 1.701620133 | 1.345465093 | 0.429852095 | 0.440481323 | 0.463935617 | 0.200214967 | 0.852870523 | 0.98092127 | 0.82776323 | 1.323132374 |
| ENSG00000135047 | CTSL | 6074.285076 | 2.106677476 | 2.099814246 | 1.798227965 | 1.452103657 | 0.551367402 | 0.484131695 | 0.489096745 | 0.337989594 | 0.640936769 | 0.692885241 | 0.648206139 | 0.698563071 |
| ENSG00000102580 | DNAJC3 | 1888.224704 | 1.170311576 | 0.990785608 | 1.171223589 | 1.265621762 | 0.955426189 | 0.79877017 | 0.709932728 | 1.290416064 | 0.944547991 | 0.868261294 | 0.922398761 | 0.912304268 |
| ENSG00000137976 | DNASE2B | 30.21236038 | 3.067056127 | 1.488356003 | 2.184616638 | 3.724712869 | 0.285706338 | 0.115293005 | 0.581516014 | 0.130429587 | 0 | 0.110015117 | 0.212500915 | 0.099797385 |
| ENSG00000149927 | DOC2A | 107.023 | 3.538582105 | 2.68706763 | 2.931496549 | 2.447076888 | 0.02304408 | 0.06509393 | 0.065664283 | 0.135006627 | 0.026893198 | 0.062114057 | 0.008569792 | 0.009390862 |
| ENSG00000176978 | DPP7 | 4219.542168 | 1.463711104 | 1.416360895 | 1.37028925 | 1.201574594 | 0.78437487 | 0.705191831 | 0.786109408 | 0.450757283 | 0.866506713 | 1.036048299 | 0.899875507 | 1.019200245 |
| ENSG00000136048 | DRAM1 | 847.86498 | 0.937278939 | 1.126074448 | 1.158084265 | 1.087133463 | 1.118423112 | 1.216053354 | 1.138642354 | 1.415986198 | 0.712873 | 0.663497059 | 0.693391883 | 0.732561924 |
| ENSG00000101210 | EEF1A2 | 220.1970516 | 3.627272978 | 2.664246389 | 2.808546608 | 2.778270037 | 0.01120018 | 0.007909447 | 0.011968124 | 0.047721931 | 0 | 0.007547368 | 0.012495602 | 0.022821336 |
| ENSG00000138798 | EGF | 107.8495662 | 1.326137312 | 1.212033531 | 1.643140507 | 1.489244554 | 0.651722844 | 1.243454629 | 1.645315945 | 1.778172852 | 0.329140736 | 0.246552044 | 0.127561694 | 0.307523353 |
| ENSG00000115363 | EVA1A | 684.3181733 | 1.287859637 | 1.016217328 | 1.297449453 | 1.02703072 | 1.414549254 | 1.30689289 | 1.648250779 | 0.602713746 | 0.556584038 | 0.699425467 | 0.455688706 | 0.687337982 |
| ENSG00000052795 | FNIP2 | 1359.824129 | 1.49322873 | 1.072020412 | 1.423544204 | 1.428653772 | 0.613920897 | 0.935610523 | 1.066548648 | 0.864529494 | 0.797954588 | 0.834116937 | 0.727082077 | 0.742789717 |
| ENSG00000167996 | FTH1 | 23002.64179 | 1.372883832 | 1.426450565 | 1.481366121 | 1.105003199 | 1.262037122 | 0.710165536 | 0.61549259 | 1.766493021 | 0.648644658 | 0.441691689 | 0.571886314 | 0.597885352 |
| ENSG00000087086 | FTL | 60070.32334 | 1.571951435 | 1.657887589 | 1.36910597 | 1.237071091 | 1.367020733 | 0.770004065 | 0.539408681 | 1.527725867 | 0.555399499 | 0.410079932 | 0.44491495 | 0.549430189 |
| ENSG00000179163 | FUCA1 | 753.2002226 | 1.937655202 | 1.525844501 | 1.687764758 | 1.574192239 | 0.703986785 | 0.702944059 | 0.720766138 | 0.664437382 | 0.545169617 | 0.646493567 | 0.524824683 | 0.76592107 |
| ENSG00000171298 | GAA | 1795.864962 | 1.310140023 | 1.216431861 | 1.117677042 | 1.03506118 | 1.311493615 | 0.946043742 | 1.017433416 | 1.359706691 | 0.631454574 | 0.659816056 | 0.658814963 | 0.735926836 |
| ENSG00000102393 | GLA | 1300.97157 | 1.283616447 | 1.205720034 | 1.289873734 | 1.218845513 | 0.787661664 | 0.730271866 | 0.869014615 | 0.657283461 | 1.028737843 | 0.991929214 | 0.875590844 | 1.061454765 |
| ENSG00000170266 | GLB1 | 2048.021645 | 2.11570161 | 1.684499155 | 1.941590847 | 1.607445541 | 0.576214107 | 0.507263398 | 0.535727865 | 0.59646907 | 0.55839308 | 0.643089999 | 0.569639361 | 0.663965966 |
| ENSG00000166105 | GLB1L3 | 16.6791658 | 1.811613222 | 1.630129273 | 2.873019522 | 1.104035276 | 1.18291121 | 0 | 0.210669665 | 2.047568993 | 0 | 0.548018716 | 0.10997731 | 0.482056813 |
| ENSG00000196743 | GM2A | 2744.568033 | 2.118953038 | 1.795752432 | 1.945942501 | 1.563661471 | 0.53511146 | 0.585079042 | 0.697428305 | 0.556602607 | 0.50267027 | 0.618847901 | 0.450467344 | 0.62948363 |
| ENSG00000127955 | GNAI1 | 571.8395963 | 1.48657483 | 1.238047959 | 1.658587153 | 1.339961635 | 0.75690153 | 0.977661311 | 0.941678359 | 0.633978281 | 0.661028694 | 0.850077797 | 0.69448216 | 0.761020291 |
| ENSG00000063660 | GPC1 | 740.7442442 | 1.808430855 | 1.269154314 | 1.563571974 | 1.187722371 | 0.875636687 | 0.960464674 | 1.265355256 | 0.368837077 | 0.546565833 | 0.796465731 | 0.683468044 | 0.674327184 |
| ENSG00000147257 | GPC3 | 5.248042646 | 3.454567655 | 1.793362888 | 1.37825528 | 1.949339195 | 0 | 0.49779614 | 0 | 2.00231386 | 0 | 0.158336002 | 0 | 0.76602898 |
| ENSG00000076716 | GPC4 | 65.55989931 | 3.164808929 | 1.850302499 | 3.075414222 | 2.730766343 | 0.357373062 | 0.225807451 | 0.120592576 | 0.220391037 | 0.058535594 | 0.076048386 | 0.027979448 | 0.091980451 |
| ENSG00000183098 | GPC6 | 114.5321166 | 3.042757662 | 2.017845967 | 2.676143791 | 3.19772057 | 0.269165388 | 0.083635956 | 0.199417102 | 0.355528098 | 0.075389964 | 0.007255206 | 0.040039638 | 0.035100659 |
| ENSG00000101850 | GPR143 | 3879.367647 | 1.560645776 | 1.542176327 | 1.467607042 | 1.179567313 | 0.947561763 | 1.046274856 | 1.193344993 | 0.552584549 | 0.550011991 | 0.753121348 | 0.442580552 | 0.764523491 |
| ENSG00000030582 | GRN | 10824.50595 | 1.07024654 | 1.169251603 | 1.010180345 | 0.9648502 | 1.29264557 | 0.811567441 | 0.644279122 | 1.70238645 | 0.864072735 | 0.783243563 | 0.775876418 | 0.911400013 |
| ENSG00000173805 | HAP1 | 13.65238416 | 2.50835482 | 1.991534673 | 3.11262206 | 1.948272133 | 0 | 0.191355249 | 0.128687936 | 1.731823096 | 0.070273214 | 0.243460507 | 0 | 0.073616313 |
| ENSG00000051620 | HEBP2 | 734.2981276 | 0.894321616 | 1.307354255 | 0.858217548 | 0.948766762 | 1.828782047 | 1.030564032 | 0.63165195 | 2.041045779 | 0.663727185 | 0.452652165 | 0.612027703 | 0.730888958 |
| ENSG00000204257 | HLA-DMA | 522.1778792 | 1.323196058 | 1.433895746 | 1.412891206 | 1.099079638 | 1.461768738 | 1.202387409 | 1.012731599 | 1.53947441 | 0.422578776 | 0.297577553 | 0.286296841 | 0.508122026 |
| ENSG00000242574 | HLA-DMB | 310.6269841 | 0.953293656 | 1.016695232 | 1.100245632 | 1.004489566 | 1.413244515 | 1.160616622 | 1.156646019 | 2.308835464 | 0.509616028 | 0.382537389 | 0.434035824 | 0.559744052 |
| ENSG00000204252 | HLA-DOA | 50.09525269 | 0.965079784 | 1.315124904 | 0.902423645 | 0.898547159 | 1.969245701 | 1.286360729 | 0.490995825 | 2.281193908 | 0.497937794 | 0.331749636 | 0.439402063 | 0.62193885 |
| ENSG00000231389 | HLA-DPA1 | 1323.902853 | 1.765023206 | 1.536337078 | 1.97642092 | 1.56555415 | 1.295619581 | 0.8583848 | 0.68608948 | 1.487253628 | 0.263056369 | 0.113605534 | 0.180121125 | 0.272534128 |
| ENSG00000223865 | HLA-DPB1 | 619.2602897 | 1.284909122 | 1.535018719 | 1.333013597 | 0.95485929 | 1.810077385 | 1.177009184 | 0.791548093 | 2.036280131 | 0.339288545 | 0.119424603 | 0.251781091 | 0.366790241 |
| ENSG00000232629 | HLA-DQB2 | 43.58897383 | 1.317093976 | 1.727344158 | 1.991276742 | 0.962259002 | 0.650665353 | 0.539404722 | 0.141070997 | 3.646265157 | 0.462211731 | 0.17157061 | 0.252494558 | 0.138342994 |
| ENSG00000204287 | HLA-DRA | 1537.799621 | 0.998823259 | 1.341004345 | 1.283485656 | 1.090345096 | 1.910066547 | 0.831858908 | 0.384442731 | 2.69658649 | 0.571470792 | 0.147516272 | 0.327431167 | 0.416968736 |
| ENSG00000196126 | HLA-DRB1 | 506.8780244 | 0.904120854 | 1.376085426 | 1.072032724 | 0.934463406 | 1.887838111 | 0.776537793 | 0.370874223 | 2.987895241 | 0.662464936 | 0.13606664 | 0.39988603 | 0.491734616 |
| ENSG00000198502 | HLA-DRB5 | 138.4792977 | 0.967353798 | 1.76707112 | 1.390694651 | 1.137681351 | 1.585045161 | 0.798631606 | 0.374268691 | 2.390315405 | 0.637384194 | 0.102009613 | 0.450372103 | 0.399172305 |
| ENSG00000204642 | HLA-F | 3247.719304 | 1.725858694 | 1.918425932 | 1.533111083 | 1.388191352 | 1.326633332 | 0.560932778 | 0.310242485 | 1.703528477 | 0.527891149 | 0.212873632 | 0.334929672 | 0.457381413 |
| ENSG00000080824 | HSP90AA1 | 66193.56163 | 1.270999222 | 1.27415545 | 1.185908193 | 0.954098996 | 1.384306181 | 1.164563283 | 1.040877011 | 0.626248943 | 0.746895019 | 0.791039792 | 0.779832254 | 0.781075656 |
| ENSG00000114378 | HYAL1 | 92.66574163 | 2.260802797 | 2.313437011 | 2.048977182 | 1.854705016 | 0.465752644 | 0.610831993 | 0.360230707 | 0.566996408 | 0.320952529 | 0.2690166 | 0.386004904 | 0.542292206 |
| ENSG00000127415 | IDUA | 325.517599 | 2.212331082 | 2.059238172 | 2.052612636 | 1.662516519 | 0.803096768 | 0.438729664 | 0.572107613 | 0.589139178 | 0.271151286 | 0.444172643 | 0.391640637 | 0.503263803 |
| ENSG00000142089 | IFITM3 | 3079.003858 | 0.883553606 | 1.270915201 | 0.943490487 | 0.943610745 | 1.831460089 | 0.874776912 | 0.473602727 | 2.316483006 | 0.721338441 | 0.425866813 | 0.578179546 | 0.736722426 |
| ENSG00000118162 | KPTN | 433.2480231 | 1.267005399 | 1.342026916 | 1.391958468 | 1.338847895 | 0.831098986 | 0.936649169 | 1.137476967 | 0.891354093 | 0.704188364 | 0.698138881 | 0.776921064 | 0.6843338 |
| ENSG00000109270 | LAMTOR3 | 1143.059193 | 1.114654715 | 1.1371701 | 1.149301377 | 1.204650624 | 1.176962215 | 0.83344362 | 0.658610182 | 1.419180905 | 0.977812352 | 0.697878058 | 0.787132253 | 0.843203598 |
| ENSG00000188186 | LAMTOR4 | 1905.781378 | 1.157416285 | 1.535315574 | 0.954535161 | 0.893770322 | 2.00583454 | 0.914784125 | 0.606595452 | 1.158601687 | 0.756631148 | 0.631353386 | 0.596754947 | 0.788407374 |
| ENSG00000113441 | LNPEP | 1350.884225 | 1.084834319 | 1.084533644 | 1.081584014 | 1.038255153 | 1.326337274 | 1.057190722 | 0.799840371 | 1.980671356 | 0.659775065 | 0.614503541 | 0.618511343 | 0.653963198 |
| ENSG00000198589 | LRBA | 1568.510609 | 1.75818486 | 1.045397897 | 1.60536863 | 1.616865287 | 0.408810813 | 0.804467586 | 1.137466991 | 0.687535645 | 0.606767801 | 0.893196764 | 0.809274864 | 0.626662863 |
| ENSG00000081479 | LRP2 | 86.95470165 | 1.308892342 | 0.853863214 | 1.029388136 | 0.635309663 | 1.077772291 | 1.351973952 | 2.060883479 | 2.462260835 | 0.187566021 | 0.506477121 | 0.063285702 | 0.462327244 |
| ENSG00000171017 | LRRC8E | 4.194490855 | 2.881512759 | 2.991748062 | 1.939993571 | 1.707275306 | 1.469931996 | 0.4152201 | 0 | 0 | 0 | 0.594318206 | 0 | 0 |
| ENSG00000003056 | M6PR | 3325.55619 | 1.750883427 | 1.435802288 | 1.963496932 | 1.824826684 | 0.555461566 | 0.531830522 | 0.537282874 | 0.732293133 | 0.673340715 | 0.702382345 | 0.623292025 | 0.669107489 |
| ENSG00000013288 | MAN2B2 | 890.9098718 | 1.279767887 | 1.076362574 | 1.137650533 | 1.088577458 | 1.211102143 | 0.807372404 | 0.709929241 | 1.234045603 | 0.885189718 | 0.943895189 | 0.807104397 | 0.819002851 |
| ENSG00000139266 | MARCHF9 | 753.4418149 | 1.11623416 | 1.133953802 | 1.03921452 | 0.939595971 | 1.510631278 | 1.322220634 | 1.202057619 | 0.672941183 | 0.699070442 | 0.88340495 | 0.640299918 | 0.840375523 |
| ENSG00000008394 | MGST1 | 190.2789927 | 3.160108836 | 3.16558999 | 2.717954727 | 2.21508881 | 0.129612129 | 0.073224559 | 0.013849904 | 0.400383961 | 0.030252322 | 0.021835151 | 0.019280424 | 0.052819188 |
| ENSG00000100427 | MLC1 | 26.69642217 | 1.207300748 | 1.488516178 | 1.625643115 | 1.264578079 | 1.247145699 | 0.717624479 | 0.032905105 | 2.902942758 | 0.431247409 | 0.311260471 | 0.206132092 | 0.564703867 |
| ENSG00000108960 | MMD | 927.1151132 | 0.836518629 | 0.651954372 | 0.743118963 | 0.648826286 | 1.620019038 | 1.934911852 | 2.372558903 | 1.137683362 | 0.447042075 | 0.495642458 | 0.61631368 | 0.495410382 |
| ENSG00000118242 | MREG | 1232.956664 | 1.675469015 | 1.234065483 | 1.578822295 | 1.349971224 | 0.777104989 | 0.77903205 | 0.8663674 | 0.500713808 | 0.761008236 | 1.006884874 | 0.535589563 | 0.934971063 |
| ENSG00000196586 | MYO6 | 610.6459223 | 1.301385893 | 0.873380998 | 1.234848341 | 1.323495292 | 0.777459472 | 1.048154982 | 1.329226067 | 0.84966388 | 0.727427716 | 0.877702398 | 0.791532179 | 0.865722783 |
| ENSG00000108784 | NAGLU | 801.1364423 | 2.255456752 | 1.88749074 | 1.923092332 | 1.937152735 | 0.634157626 | 0.467400953 | 0.618427732 | 0.278728871 | 0.473030259 | 0.495790774 | 0.478538866 | 0.550732362 |
| ENSG00000131400 | NAPSA | 18.43183692 | 1.038253812 | 0.737560682 | 2.158346118 | 1.498579943 | 0.468312674 | 0.094490686 | 0.476593067 | 4.275846965 | 0.104102148 | 0.676238155 | 0.199039281 | 0.27263647 |
| ENSG00000188554 | NBR1 | 4899.393961 | 1.243540044 | 1.077243709 | 1.071818573 | 1.085789786 | 1.193760633 | 0.873592275 | 0.709838382 | 1.44586445 | 0.842219698 | 0.772882898 | 0.76321302 | 0.920236534 |
| ENSG00000114388 | NPRL2 | 576.4392032 | 1.137485927 | 1.191886456 | 1.10422146 | 1.110978347 | 0.986174814 | 0.711532961 | 0.797011374 | 1.428741152 | 0.92870761 | 0.782750497 | 0.943514559 | 0.876994844 |
| ENSG00000103148 | NPRL3 | 1359.293112 | 1.373773601 | 1.206301774 | 1.539172417 | 1.26890534 | 0.626861247 | 0.886006054 | 0.973904732 | 0.406825814 | 0.850495941 | 1.083247344 | 0.945306057 | 0.83919968 |
| ENSG00000073969 | NSF | 973.611024 | 1.21761372 | 1.013933648 | 1.282464137 | 1.185245698 | 0.671269442 | 1.087616324 | 1.207221525 | 1.073908073 | 0.76072928 | 0.852622987 | 0.874198215 | 0.773176952 |
| ENSG00000168824 | NSG1 | 962.2111055 | 2.678665434 | 2.087757363 | 2.417722512 | 2.407081684 | 0.501086708 | 0.45974919 | 0.405348849 | 0.271657834 | 0.129619786 | 0.291029201 | 0.072442036 | 0.277839402 |
| ENSG00000104044 | OCA2 | 1902.962228 | 2.527858166 | 2.074482674 | 1.943731821 | 1.873515908 | 0.374545136 | 0.469052357 | 0.43577084 | 0.236757352 | 0.460297826 | 0.702154862 | 0.278094927 | 0.62373813 |
| ENSG00000109180 | OCIAD1 | 2307.432533 | 1.347491923 | 1.207789434 | 1.2922861 | 1.289732015 | 0.831013105 | 0.812536059 | 0.89084713 | 0.704743624 | 0.959217067 | 0.841962069 | 0.918582898 | 0.903798578 |
| ENSG00000169174 | PCSK9 | 98.04127661 | 4.016856828 | 3.146560471 | 2.351626016 | 1.753012831 | 0.100620743 | 0.1154681 | 0.134399806 | 0.187567936 | 0.009785643 | 0.042377768 | 0.01870977 | 0.123014088 |
| ENSG00000154678 | PDE1C | 131.4484868 | 2.344684369 | 1.360390353 | 2.992062763 | 1.984583427 | 0.225144914 | 0.722101974 | 0.882134241 | 0.679505293 | 0.175167675 | 0.164359491 | 0.362823304 | 0.107042196 |
| ENSG00000129925 | PGAP6 | 3407.232671 | 1.373100988 | 1.273706395 | 1.124595119 | 1.096512907 | 1.007566989 | 0.756259423 | 0.755666843 | 1.14265712 | 0.849234952 | 0.875283413 | 0.904450717 | 0.840965133 |
| ENSG00000127252 | PLAAT1 | 58.83350709 | 2.85897453 | 3.448260096 | 2.581793942 | 2.277882556 | 0.356312185 | 0.266425257 | 0.179173117 | 0 | 0 | 0 | 0.031178318 | 0 |
| ENSG00000075651 | PLD1 | 1043.567042 | 1.750795739 | 1.226546697 | 1.509262666 | 1.395964601 | 0.573096656 | 0.735996681 | 0.860293974 | 0.616758648 | 0.752942586 | 0.908536378 | 0.896452409 | 0.773352965 |
| ENSG00000166289 | PLEKHF1 | 340.9157729 | 2.109452107 | 1.588933381 | 2.376281834 | 1.722461678 | 0.53894651 | 0.541522357 | 0.626145867 | 0.435382402 | 0.402427135 | 0.587417633 | 0.513846845 | 0.557182251 |
| ENSG00000163932 | PRKCD | 4876.090333 | 1.044781888 | 1.076387083 | 0.951409109 | 1.075453482 | 1.382559472 | 0.85133607 | 0.672155638 | 1.790309501 | 0.890908269 | 0.717783798 | 0.73770592 | 0.809209769 |
| ENSG00000080815 | PSEN1 | 2207.641669 | 0.896960842 | 1.06201353 | 1.017325761 | 1.007895368 | 1.565672797 | 0.824413944 | 0.578565008 | 2.01642702 | 0.859164652 | 0.625951519 | 0.755288231 | 0.790321328 |
| ENSG00000143801 | PSEN2 | 1329.600907 | 1.606711561 | 1.411777235 | 1.679626368 | 1.583466349 | 0.486905288 | 0.641193357 | 0.646811494 | 0.34181752 | 0.873096771 | 1.028067502 | 0.767752578 | 0.932773977 |
| ENSG00000103490 | PYCARD | 1657.730371 | 1.510448015 | 1.692504385 | 1.295348877 | 1.071323416 | 1.786757442 | 0.954484012 | 0.663977735 | 0.861301437 | 0.525497033 | 0.54787729 | 0.394478245 | 0.696002113 |
| ENSG00000069974 | RAB27A | 4042.731585 | 1.538938436 | 1.540645805 | 1.375426038 | 1.459604236 | 1.105705816 | 0.875184344 | 1.034739012 | 0.770364335 | 0.557687961 | 0.630399558 | 0.367979279 | 0.743325181 |
| ENSG00000179331 | RAB39A | 28.84072724 | 1.606460965 | 1.015254325 | 1.442077689 | 0.673956963 | 1.496471079 | 1.811643201 | 1.857975433 | 1.912857444 | 0.033265351 | 0.086435486 | 0.063602064 | 0 |
| ENSG00000123595 | RAB9A | 631.7931212 | 1.469855293 | 1.587328223 | 1.383848713 | 1.256526346 | 0.948566684 | 0.694677557 | 0.756380539 | 0.663216484 | 0.824561876 | 0.750997077 | 0.746166329 | 0.91787488 |

| | | | | | | | | | | | | | | |
|---|---|---|---|---|---|---|---|---|---|---|---|---|---|---|
| **ENSG00000167705** | RILP | *206.6893982* | 1.1792767 | 1.533018698 | 1.233581765 | 1.128499619 | 1.563110143 | 0.985882782 | 0.561011895 | 1.849329555 | 0.645200825 | 0.313583666 | 0.599049885 | 0.408454467 |
| **ENSG00000108523** | RNF167 | *2889.434313* | 1.374461261 | 1.300733337 | 1.372751231 | 1.364177725 | 0.958524939 | 0.665748988 | 0.820552548 | 1.209228384 | 0.754718376 | 0.687900113 | 0.751016253 | 0.740186845 |
| **ENSG00000025039** | RRAGD | *2247.481767* | 1.126198292 | 1.10925935 | 1.029463948 | 0.931309907 | 1.376061505 | 1.08334957 | 1.10495851 | 1.36292479 | 0.808930763 | 0.683255004 | 0.557444897 | 0.826843465 |
| **ENSG00000115884** | SDC1 | *258.040156* | 2.603497056 | 2.455886844 | 3.710613306 | 2.164662122 | 0.205488561 | 0.182235964 | 0.289366312 | 0.142531304 | 0.022308084 | 0.083726528 | 0.046206543 | 0.093477375 |
| **ENSG00000169439** | SDC2 | *355.1003635* | 1.80678663 | 1.540187506 | 1.555704567 | 1.24168354 | 1.257082974 | 1.495913024 | 1.244323226 | 0.843379911 | 0.27287803 | 0.184864281 | 0.240203457 | 0.316992853 |
| **ENSG00000133661** | SFTPD | *3.300209038* | 0.610389563 | 1.267481316 | 1.095861262 | 2.789881974 | 0 | 0 | 0.266179675 | 5.97020621 | 0 | 0 | 0 | 0 |
| **ENSG00000110013** | SIAE | *195.4401672* | 2.107793375 | 1.503541842 | 1.905989148 | 1.627913536 | 0.378568013 | 0.739642551 | 0.943890921 | 0.470461118 | 0.51543486 | 0.590987107 | 0.567830827 | 0.647946702 |
| **ENSG00000149577** | SIDT2 | *489.6824045* | 1.404833184 | 1.161735939 | 1.384789614 | 1.380930212 | 1.115567152 | 0.851821583 | 0.870048733 | 1.40021758 | 0.664176513 | 0.643134404 | 0.556274782 | 0.566470303 |
| **ENSG00000181045** | SLC26A11 | *655.4103486* | 1.303170111 | 1.281224619 | 1.045665897 | 1.145691092 | 0.950133377 | 1.114746947 | 1.29339256 | 0.941941835 | 0.750935067 | 0.797470052 | 0.654907525 | 0.720720919 |
| **ENSG00000198246** | SLC29A3 | *50.6729866* | 1.729264013 | 2.043062803 | 1.998382207 | 1.816982642 | 0.267684161 | 0.910808328 | 0.20802766 | 1.607143699 | 0.549060006 | 0.508349293 | 0.162897129 | 0.198338061 |
| **ENSG00000115194** | SLC30A3 | *27.7919986* | 2.754307127 | 2.633912146 | 2.309806519 | 2.76074475 | 0.044369722 | 0 | 0 | 0.945256644 | 0.172603079 | 0.089697122 | 0 | 0.28930289 |
| **ENSG00000104154** | SLC30A4 | *220.9983353* | 0.91606356 | 0.809152038 | 1.039158388 | 0.893414666 | 1.026680495 | 1.595855799 | 1.709211448 | 1.468072397 | 0.677226448 | 0.624160265 | 0.622514796 | 0.618489698 |
| **ENSG00000213699** | SLC35F6 | *1370.121775* | 1.202659493 | 1.032670218 | 1.191118777 | 1.084157101 | 0.897309944 | 1.082388338 | 1.402828203 | 1.028679477 | 0.777952722 | 0.767199636 | 0.789227225 | 0.743808866 |
| **ENSG00000123643** | SLC36A1 | *945.0731827* | 1.137149415 | 0.854230128 | 1.213083923 | 1.271912389 | 0.899001203 | 0.825600971 | 0.999215964 | 1.655334794 | 0.821261372 | 0.76318762 | 0.821987839 | 0.738034383 |
| **ENSG00000138821** | SLC39A8 | *814.396202* | 1.684456968 | 1.35340537 | 1.557610853 | 1.311443333 | 0.676827933 | 0.867186962 | 1.055998477 | 0.70967002 | 0.593735629 | 0.741780996 | 0.666704503 | 0.781178955 |
| **ENSG00000168003** | SLC3A2 | *7085.796555* | 1.688533791 | 1.669303668 | 1.610430997 | 1.357426548 | 0.71734124 | 0.706039469 | 0.717680606 | 0.578184198 | 0.727082884 | 0.767182862 | 0.76833857 | 0.692455167 |
| **ENSG00000103257** | SLC7A5 | *16356.79842* | 1.960742621 | 1.629586882 | 2.080985747 | 1.759366026 | 0.209279964 | 0.363462179 | 0.480341189 | 0.232964072 | 0.827788435 | 0.955733448 | 0.793872993 | 0.705876445 |
| **ENSG00000145335** | SNCA | *2703.218446* | 1.373013801 | 1.282405626 | 1.349248006 | 1.156151756 | 0.63635514 | 0.872358793 | 0.949544689 | 0.538391432 | 0.951511379 | 0.961529545 | 0.812251319 | 1.117238514 |
| **ENSG00000028528** | SNX1 | *3277.933227* | 1.023819608 | 0.962175422 | 1.11406565 | 1.036150288 | 1.185748536 | 1.165188394 | 1.17084197 | 1.079936424 | 0.815416178 | 0.874319417 | 0.812258669 | 0.760079446 |
| **ENSG00000129515** | SNX6 | *1757.915965* | 1.087468411 | 1.140373264 | 1.064655908 | 1.08592117 | 1.090784037 | 0.885721179 | 0.71258677 | 1.513095847 | 0.901046652 | 0.714238712 | 0.849905022 | 0.954203028 |
| **ENSG00000008294** | SPAG9 | *6798.71896* | 1.109913752 | 0.948571088 | 1.163638273 | 1.206640492 | 1.222473063 | 1.059140385 | 1.105244841 | 1.721626254 | 0.69992725 | 0.547799706 | 0.582914152 | 0.632110744 |
| **ENSG00000104133** | SPG11 | *1970.009134* | 1.134508137 | 0.990527241 | 0.958295145 | 0.953430867 | 1.260659257 | 1.043209152 | 0.879336318 | 1.969616952 | 0.79186403 | 0.690911926 | 0.631770044 | 0.695870931 |
| **ENSG00000063176** | SPHK2 | *536.0046668* | 1.272150961 | 1.279847701 | 1.561994315 | 1.555513586 | 0.821308157 | 0.984536176 | 0.947273978 | 0.44355709 | 0.737440456 | 0.998376449 | 0.732356634 | 0.665644498 |
| **ENSG00000131748** | STARD3 | *1678.630219* | 1.110031349 | 1.185514241 | 1.122482841 | 1.090894531 | 1.300978186 | 0.83780918 | 0.628498593 | 1.431192957 | 0.843014991 | 0.782625266 | 0.842513286 | 0.82444458 |
| **ENSG00000101846** | STS | *299.6053665* | 1.398499435 | 1.183240794 | 1.614511814 | 1.673136073 | 0.674995303 | 1.002760301 | 1.307680387 | 0.868070753 | 0.438701676 | 0.535284604 | 0.648983559 | 0.654135302 |
| **ENSG00000166900** | STX3 | *5596.04403* | 1.004138899 | 1.171494277 | 1.016264034 | 1.191017295 | 1.533018413 | 0.893219909 | 0.593528932 | 1.963470337 | 0.786232952 | 0.51822855 | 0.639191374 | 0.690195027 |
| **ENSG00000170310** | STX8 | *379.7209591* | 1.816956605 | 1.867188435 | 1.21196547 | 1.031855719 | 0.798871686 | 0.786605858 | 0.809691937 | 0.629573357 | 0.613959917 | 0.680570079 | 0.850208644 | 0.902552293 |
| **ENSG00000198203** | SULT1C2 | *54.33973251* | 2.539344501 | 2.444045287 | 2.079838197 | 1.506111975 | 0.068078542 | 0.689094184 | 0.678966167 | 1.474524051 | 0.158899792 | 0.091750996 | 0.084391738 | 0.184954571 |
| **ENSG00000100321** | SYNGR1 | *1347.182041* | 1.855641366 | 1.62001557 | 2.084549441 | 1.7146774 | 0.804579719 | 0.756934388 | 0.959185771 | 0.405608052 | 0.336847378 | 0.552660921 | 0.398269458 | 0.511030535 |
| **ENSG00000011347** | SYT7 | *13.10039217* | 1.99897611 | 1.037724523 | 2.070494071 | 1.093272715 | 1.223673475 | 0 | 0 | 3.208523346 | 0.219702638 | 0.317148556 | 0.140020982 | 0.690463582 |
| **ENSG00000185339** | TCN2 | *155.7673749* | 1.939828369 | 1.382973134 | 2.19407936 | 1.300389517 | 0.894557854 | 0.486374029 | 0.817726064 | 0.303574663 | 0.51736983 | 0.666822958 | 0.612356398 | 0.883947823 |
| **ENSG00000205356** | TECPR1 | *1677.279908* | 1.17337699 | 1.139084722 | 1.074332682 | 1.147276299 | 1.191746708 | 1.005142031 | 0.748939666 | 1.358730914 | 0.821385836 | 0.818925398 | 0.728906841 | 0.792151912 |
| **ENSG00000164342** | TLR3 | *126.9742124* | 1.911701441 | 1.358912334 | 2.043635331 | 1.45024624 | 0.41759897 | 0.596666081 | 0.940891879 | 0.868966993 | 0.619578928 | 0.608617523 | 0.621198424 | 0.561985856 |
| **ENSG00000244045** | TMEM199 | *371.5305229* | 1.309396527 | 1.362303546 | 1.496641041 | 1.3134352 | 0.740145077 | 0.799258945 | 0.799168838 | 0.72476779 | 0.854735633 | 0.881208656 | 0.891169111 | 0.827769637 |
| **ENSG00000164841** | TMEM74 | *26.01579777* | 2.20676588 | 1.849028937 | 2.29373806 | 2.477354587 | 0.142197054 | 0.401672152 | 0.168829835 | 0.252448258 | 0.516284637 | 0.479105485 | 0.246779065 | 0.96579605 |
| **ENSG00000175348** | TMEM9B | *854.7614803* | 1.279686049 | 1.317630942 | 1.277788647 | 1.199243987 | 1.058906485 | 0.895512305 | 0.95988294 | 0.58395338 | 0.853035223 | 0.88562622 | 0.856259436 | 0.832474385 |
| **ENSG00000162341** | TPCN2 | *3215.668596* | 1.883695465 | 1.571045551 | 2.166398728 | 1.798107447 | 0.411466923 | 0.553524989 | 0.608093296 | 0.295329299 | 0.567462991 | 0.817344113 | 0.533356686 | 0.794174512 |
| **ENSG00000170043** | TRAPPC1 | *1343.441904* | 1.35924413 | 1.317835164 | 1.336587474 | 1.044768311 | 0.871071516 | 0.750615095 | 0.774846721 | 0.721567549 | 0.876955973 | 0.999538432 | 0.925055578 | 1.021914057 |
| **ENSG00000113595** | TRIM23 | *619.3007726* | 1.16284806 | 0.866241064 | 1.040938188 | 1.116682846 | 1.089161284 | 1.098188869 | 1.048236206 | 1.493174483 | 0.878374565 | 0.762120678 | 0.731598403 | 0.712435354 |
| **ENSG00000142185** | TRPM2 | *112.8549104* | 0.70505855 | 0.963686784 | 0.881270552 | 0.979012115 | 1.901232715 | 0.910519334 | 0.420329275 | 3.56156183 | 0.527071512 | 0.18407575 | 0.511997111 | 0.45418447 |
| **ENSG00000103197** | TSC2 | *2794.597615* | 1.400561116 | 1.098276891 | 1.366924295 | 1.349704277 | 0.848528611 | 0.799274045 | 0.804705595 | 0.996451348 | 0.815347313 | 0.971716988 | 0.819167509 | 0.729342012 |
| **ENSG00000077498** | TYR | *6937.863625* | 2.125076344 | 1.606471337 | 1.707714079 | 1.456411387 | 0.450388551 | 0.509221667 | 0.735515717 | 0.311822438 | 0.668327529 | 0.890496559 | 0.582460067 | 0.956094326 |
| **ENSG00000221983** | UBA52 | *11861.3658* | 1.339192736 | 1.568515029 | 1.336925361 | 1.077671336 | 1.118519858 | 0.735485226 | 0.704751605 | 0.676068172 | 0.833107108 | 0.774884055 | 0.891387809 | 0.943491705 |
| **ENSG00000114316** | USP4 | *2636.455817* | 1.023077722 | 1.065786446 | 1.028130314 | 0.978609338 | 1.358726763 | 0.916579448 | 0.800995923 | 1.75920618 | 0.875170958 | 0.686458693 | 0.772289088 | 0.734969127 |
| **ENSG00000111667** | USP5 | *2432.122313* | 1.694606101 | 1.453726753 | 1.815629692 | 1.549597761 | 0.610953456 | 0.638759045 | 0.768242656 | 0.396955158 | 0.673753088 | 0.810412823 | 0.891475643 | 0.695887827 |
| **ENSG00000129204** | USP6 | *27.22461404* | 1.294866114 | 1.766929104 | 1.726945553 | 0.901849941 | 0.860594092 | 0.511783024 | 0.709867493 | 1.881665393 | 0.634320998 | 0.457832437 | 0.404265738 | 0.849080113 |
| **ENSG00000160695** | VPS11 | *966.1609263* | 1.142561635 | 1.004434291 | 1.191285806 | 1.042969316 | 0.860234622 | 1.103213514 | 1.502933357 | 1.137929904 | 0.66729538 | 0.839416264 | 0.75563318 | 0.752092732 |
| **ENSG00000104142** | VPS18 | *2498.331746* | 1.079236979 | 1.100432742 | 1.171103933 | 1.092088113 | 1.164845663 | 1.058576651 | 0.965180595 | 1.138275311 | 0.78991889 | 0.845478311 | 0.736791637 | 0.858071175 |
| **ENSG00000069329** | VPS35 | *3590.873679* | 1.375526263 | 1.121783829 | 1.250132824 | 1.354960046 | 0.76098504 | 0.76608529 | 0.843986125 | 0.812067947 | 0.934316071 | 0.936505017 | 0.931500425 | 0.912151123 |
| **ENSG00000127580** | WDR24 | *360.128809* | 1.507472691 | 1.356069787 | 1.295474504 | 1.110717625 | 0.75672991 | 0.916450411 | 0.970826331 | 0.554402583 | 0.916429145 | 1.091390848 | 0.870995002 | 0.653041163 |
| **ENSG00000164253** | WDR41 | *1060.293006* | 1.205464091 | 1.103639444 | 1.144362591 | 1.055543646 | 1.018790063 | 1.081651865 | 1.256000013 | 0.875856665 | 0.818881381 | 0.801727477 | 0.849440596 | 0.788642166 |
| **ENSG00000114742** | WDR48 | *1905.194986* | 1.911117655 | 1.343677385 | 1.968504224 | 1.943810445 | 0.599346602 | 0.536606949 | 0.48597902 | 0.871459415 | 0.64809315 | 0.572231073 | 0.580570948 | 0.538603134 |
| **ENSG00000186187** | ZNRF1 | *418.6767393* | 1.14751428 | 1.051541185 | 1.159664829 | 1.163088845 | 0.830571005 | 1.214679324 | 1.521162158 | 1.047869394 | 0.634744944 | 0.791901369 | 0.633091425 | 0.804171242 |
