## Supplementary material for "Peripheral positioning of lysosomes supports melanoma aggressiveness": Table 2

### Table 2: Positive Regulation of cell migration genes differentially expressed (refers to Figure 1A)

| Gene | GeneSymbol | *Average* | WM1862 | WM1862 | WM1862 | WM1862 | WM983A | WM983A | WM983A | WM983A | WM983B | WM983B | WM983B | WM983B |
|---|---|---|---|---|---|---|---|---|---|---|---|---|---|---|
| **ENSG00000127837** | AAMP | *3560.206563* | 0.858056265 | 0.843297903 | 0.841870651 | 0.797105914 | 0.804600882 | 0.950506793 | 1.118723237 | 0.526118837 | 1.209416714 | 1.361191882 | 1.332388088 | 1.356722834 |
| **ENSG00000143322** | ABL2 | *3545.55694* | 0.862737907 | 0.511726655 | 1.001668573 | 0.865031525 | 0.506040201 | 0.875102618 | 0.989554996 | 0.596088992 | 1.259873154 | 1.565088794 | 1.7360019 | 1.231084687 |
| **ENSG00000137845** | ADAM10 | *7478.350113* | 0.878537164 | 0.810765839 | 0.886086176 | 0.879609579 | 0.969567441 | 1.032170824 | 0.976608649 | 1.321195414 | 1.090977446 | 0.98647567 | 1.151860039 | 1.016145759 |
| **ENSG00000151694** | ADAM17 | *1147.517213* | 0.658295077 | 0.554073519 | 0.675240754 | 0.762239897 | 0.901590325 | 1.212677141 | 1.193447299 | 1.540725864 | 1.0517675 | 1.170197535 | 1.280414915 | 0.999330172 |
| **ENSG00000168615** | ADAM9 | *2738.891445* | 0.664142816 | 0.461227442 | 0.615660159 | 0.569986389 | 0.582593919 | 0.88611436 | 1.017360095 | 0.698274331 | 1.524797468 | 1.6735029 | 1.798906562 | 1.50743356 |
| **ENSG00000154734** | ADAMTS1 | *1138.655271* | 0.338785696 | 0.235109798 | 0.352555702 | 0.380043122 | 0.597796417 | 0.98273968 | 1.195792367 | 0.672536433 | 1.782878374 | 1.528863801 | 2.747494807 | 1.185403803 |
| **ENSG00000135744** | AGT | *17.35060763* | 0.696602631 | 0 | 0.885872594 | 0.294808558 | 0.710708979 | 0.200758031 | 0.354404878 | 0.984165611 | 1.382367881 | 1.819893356 | 2.643033924 | 2.027383558 |
| **ENSG00000131016** | AKAP12 | *5310.314013* | 0.788647326 | 0.550604794 | 0.823555587 | 0.707403559 | 0.609094361 | 1.139540419 | 1.324046436 | 0.676513784 | 1.194026219 | 1.6430324 | 1.119531314 | 1.4240038 |
| **ENSG00000126016** | AMOT | *492.4528976* | 0.734257353 | 0.380111805 | 0.618731514 | 0.535969134 | 0.485784355 | 0.938982394 | 1.204080201 | 0.698837363 | 1.229314421 | 1.829117549 | 1.557001396 | 1.787812516 |
| **ENSG00000135046** | ANXA1 | *2705.859771* | 0.333147303 | 0.343573374 | 0.463455679 | 0.377320174 | 1.201742263 | 1.48265652 | 1.561550846 | 0.983013006 | 1.248060654 | 1.021395618 | 1.95373703 | 1.030347532 |
| **ENSG00000138772** | ANXA3 | *243.1858912* | 0.13253487 | 0.348312987 | 0.156152143 | 0.222957592 | 1.211897854 | 0.777050034 | 0.617694984 | 1.652808636 | 1.633278633 | 1.906658242 | 1.538754052 | 1.801899972 |
| **ENSG00000142192** | APP | *22269.0729* | 0.770520233 | 0.561162695 | 0.715752006 | 0.575203671 | 0.803640008 | 1.295020002 | 1.420763327 | 0.642104502 | 1.188891751 | 1.477421018 | 1.219628857 | 1.329891929 |
| **ENSG00000137135** | ARHGEF39 | *440.3677585* | 0.91259048 | 0.786023451 | 1.090224757 | 0.836318602 | 0.46483526 | 0.654545896 | 1.095148895 | 0.653233009 | 1.259246169 | 1.651085525 | 1.291289431 | 1.305458525 |
| **ENSG00000168874** | ATOH8 | *457.7686909* | 0.292633545 | 0.269562171 | 0.481926462 | 0.181018809 | 1.21758374 | 1.862362556 | 2.153094595 | 0.880910551 | 1.035330904 | 1.472149107 | 0.897592778 | 1.255834782 |
| **ENSG00000124406** | ATP8A1 | *270.6517005* | 0.480062191 | 0.55252038 | 0.350764451 | 0.566976775 | 1.977358705 | 1.177600416 | 0.746506194 | 2.9556027 | 0.737311299 | 0.878076399 | 0.630303336 | 0.946917155 |
| **ENSG00000156735** | BAG4 | *680.5558177* | 0.822866901 | 0.763687465 | 0.944588998 | 1.022186009 | 0.677663886 | 1.064601811 | 1.024880172 | 1.080845947 | 1.122141518 | 1.117208869 | 1.115871042 | 1.243457382 |
| **ENSG00000141376** | BCAS3 | *5226.632594* | 1.091875574 | 1.043610967 | 0.95887046 | 0.801915785 | 0.732564928 | 0.715764122 | 0.72438865 | 0.349578472 | 1.30015037 | 1.480146627 | 1.290297973 | 1.510836071 |
| **ENSG00000171791** | BCL2 | *953.0533291* | 0.716524499 | 0.50583252 | 0.849068767 | 0.720261313 | 0.626231126 | 1.279200022 | 1.388110722 | 1.237652308 | 1.029809148 | 1.26946638 | 1.267406686 | 1.110436511 |
| **ENSG00000142871** | CCN1 | *894.1550064* | 0.075471076 | 0.091223097 | 0.176954768 | 0.04576484 | 0.486819968 | 1.485199028 | 1.325303902 | 0.794737992 | 1.420605484 | 1.461816776 | 3.078226726 | 1.557876342 |
| **ENSG00000177697** | CD151 | *4209.380626* | 0.938443097 | 0.881928098 | 0.958833107 | 0.72229628 | 0.77279283 | 0.76047498 | 1.160724433 | 0.488979127 | 1.196573609 | 1.326958503 | 1.568343058 | 1.223652878 |
| **ENSG00000196776** | CD47 | *2828.684112* | 0.686144863 | 0.814428701 | 0.620728674 | 0.714639899 | 1.54495471 | 1.160913085 | 1.135682278 | 1.55328516 | 0.927283161 | 0.82458417 | 0.927642907 | 1.089712391 |
| **ENSG00000002586** | CD99 | *5179.731882* | 0.551075519 | 0.541672038 | 0.527153017 | 0.400737088 | 1.257946062 | 1.169109075 | 1.015355949 | 0.47700292 | 1.440466403 | 1.514083144 | 1.414596447 | 1.690802339 |
| **ENSG00000185043** | CIB1 | *1803.519456* | 0.793581982 | 0.808285832 | 0.684303644 | 0.686355802 | 1.396863681 | 1.057428249 | 0.932260839 | 1.177683825 | 1.058596757 | 1.130195953 | 1.247961388 | 1.026482048 |
| **ENSG00000163347** | CLDN1 | *648.0037899* | 0 | 0 | 0.001395274 | 0.004736183 | 0.239772565 | 0.486472898 | 0.501580354 | 0.383110258 | 2.305204088 | 2.71212597 | 2.128715939 | 3.236886471 |
| **ENSG00000085719** | CPNE3 | *2303.742072* | 0.861293015 | 0.713578427 | 0.883835753 | 0.869488024 | 0.971514454 | 1.096204977 | 1.188555013 | 0.956748136 | 1.096100424 | 1.235389067 | 0.987336628 | 1.139956083 |
| **ENSG00000099942** | CRKL | *3380.834821* | 0.843103778 | 0.741424794 | 0.873699757 | 0.825779987 | 0.987713974 | 1.02283024 | 1.023737492 | 1.137592295 | 1.100763806 | 1.19155938 | 1.189035824 | 1.062758672 |
| **ENSG00000138061** | CYP1B1 | *90.68928002* | 0.155485765 | 0.334399075 | 0.209363213 | 0.180488194 | 0.802236742 | 0.489713242 | 0.164668036 | 1.477351168 | 1.819578545 | 1.493511881 | 2.922734125 | 1.950470014 |
| **ENSG00000162733** | DDR2 | *2514.921336* | 0.997225774 | 0.538062711 | 0.996206001 | 0.739120576 | 0.553084114 | 1.128117405 | 1.224976338 | 0.690995089 | 1.164282684 | 1.464375242 | 1.377796159 | 1.125757907 |
| **ENSG00000198171** | DDRGK1 | *2488.349066* | 0.619296578 | 0.588775671 | 0.555926224 | 0.566529703 | 1.222047987 | 1.348739362 | 1.448107132 | 0.673036169 | 1.186740125 | 1.255272614 | 1.312526535 | 1.2230019 |
| **ENSG00000087470** | DNM1L | *3113.593811* | 0.987605277 | 0.76878847 | 1.041903537 | 1.010340903 | 0.569116666 | 0.865618089 | 1.20132369 | 0.588507192 | 1.181068749 | 1.30343906 | 1.286672728 | 1.195615639 |
| **ENSG00000116641** | DOCK7 | *2884.952089* | 0.578498791 | 0.34109397 | 0.499872351 | 0.532263711 | 0.547114034 | 1.064921503 | 1.41467585 | 0.753071864 | 1.375435532 | 1.777724033 | 1.627080717 | 1.488247642 |
| **ENSG00000146648** | EGFR | *32.98892489* | 0.519038188 | 0.095099036 | 0.109629861 | 0.217077419 | 0.6728385 | 0.475151351 | 0.878743483 | 3.026111683 | 1.19237815 | 1.158688512 | 1.278901484 | 2.376342334 |
| **ENSG00000134014** | ELP3 | *1185.23113* | 0.68833606 | 0.515267584 | 0.649940467 | 0.73625923 | 0.78238638 | 0.90591542 | 1.194012381 | 0.739200592 | 1.341274824 | 1.498931141 | 1.521345609 | 1.427130311 |
| **ENSG00000161671** | EMC10 | *1058.882039* | 0.406161586 | 0.366394843 | 0.420955677 | 0.427997188 | 0.597415208 | 0.679297617 | 0.842044028 | 0.614040674 | 1.702462337 | 2.168254891 | 1.820675515 | 1.954300438 |
| **ENSG00000136960** | ENPP2 | *4056.319567* | 0.014898332 | 0.013405845 | 0.01337384 | 0.014627853 | 0.728385249 | 1.10926146 | 1.234841993 | 0.652826669 | 1.811262993 | 2.407651393 | 1.363429139 | 2.636035233 |
| **ENSG00000142627** | EPHA2 | *338.8271874* | 0.609388372 | 0.376534352 | 0.603069301 | 0.392509229 | 0.225641994 | 0.72476712 | 0.469263379 | 0.345025567 | 1.707408251 | 1.954596434 | 3.064189343 | 1.527606658 |
| **ENSG00000134954** | ETS1 | *5141.540396* | 0.708359499 | 0.550576953 | 0.793965325 | 0.742363768 | 0.6739374 | 0.901382751 | 1.13395261 | 0.805253236 | 1.300582303 | 1.378582656 | 1.412260506 | 1.598782994 |
| **ENSG00000181104** | F2R | *1755.809287* | 0.406712431 | 0.275161493 | 0.451605565 | 0.230729229 | 1.241684928 | 1.593532807 | 1.833135056 | 0.929145603 | 1.034359688 | 1.021768082 | 1.864825916 | 1.117339201 |
| **ENSG00000117525** | F3 | *131.133006* | 0.399401673 | 0.486452951 | 0.682590455 | 0.553899665 | 0.507794782 | 1.201971536 | 1.179008648 | 1.322213042 | 1.265704723 | 1.425763632 | 1.664609473 | 1.31058942 |
| **ENSG00000180921** | FAM83H | *1060.105053* | 0.997605758 | 0.800009186 | 1.137742439 | 0.846321667 | 0.556013687 | 0.916733106 | 1.023374032 | 0.455972275 | 1.196412288 | 1.497136829 | 1.131634713 | 1.44104402 |
| **ENSG00000112029** | FBXO5 | *873.8936883* | 0.64427571 | 0.62823731 | 0.857695164 | 0.687170124 | 0.447308499 | 0.80714961 | 1.562099708 | 0.667365712 | 1.315214323 | 1.387310651 | 1.737997516 | 1.258175673 |
| **ENSG00000077782** | FGFR1 | *817.0851462* | 0.706327083 | 0.520894915 | 0.663927975 | 0.647303564 | 0.780242708 | 1.221363472 | 1.638453012 | 0.726625511 | 1.086107297 | 1.405457634 | 1.496270989 | 1.10702584 |
| **ENSG00000174951** | FUT1 | *7.110292594* | 0.14165473 | 0 | 0 | 1.151031702 | 0 | 0.122472943 | 0.247092101 | 0.554208655 | 3.508198752 | 2.10359468 | 2.192849723 | 1.978896714 |
| **ENSG00000196371** | FUT4 | *253.8295481* | 0.10316912 | 0.201872391 | 0.167414363 | 0.217638816 | 0.728711416 | 0.778773753 | 1.315096917 | 1.754272453 | 1.734877535 | 1.607371808 | 1.803041782 | 1.587759648 |
| **ENSG00000179348** | GATA2 | *138.6332853* | 0.472241766 | 0.482764674 | 0.554355606 | 0.420622123 | 1.245279995 | 1.702273745 | 1.913620291 | 1.004333322 | 1.00345708 | 1.174804463 | 0.946054038 | 1.080192898 |
| **ENSG00000111846** | GCNT2 | *1217.78685* | 0.145565997 | 0.116785965 | 0.149231949 | 0.127690056 | 1.035883331 | 1.402277413 | 1.275343932 | 0.936241671 | 1.774170773 | 1.858715218 | 1.422682861 | 1.755410833 |
| **ENSG00000104522** | GFUS | *1651.193102* | 0.824092641 | 0.867652306 | 0.887062422 | 0.853141062 | 0.899902943 | 0.875990491 | 0.96931927 | 0.627651628 | 1.266069886 | 1.339637255 | 1.42752157 | 1.161958526 |
| **ENSG00000136235** | GPNMB | *24554.61229* | 0.970223308 | 0.693166594 | 0.978058259 | 0.856926855 | 0.674600136 | 1.004604517 | 1.203981875 | 0.550348005 | 1.162625337 | 1.302203966 | 1.205313019 | 1.397948129 |
| **ENSG00000170961** | HAS2 | *440.5393005* | 0.361235964 | 0.405914417 | 0.332481427 | 0.327430571 | 1.648682878 | 1.879851245 | 2.261229987 | 0.983940426 | 0.81013351 | 1.094007671 | 0.460102971 | 1.434988934 |
| **ENSG00000113070** | HBEGF | *336.7332912* | 0.4995155 | 0.593157924 | 0.630984717 | 0.519511094 | 1.00705486 | 1.184423589 | 1.205236457 | 0.635830082 | 1.37043151 | 1.332553751 | 1.922941479 | 1.098359038 |
| **ENSG00000094631** | HDAC6 | *1092.22261* | 0.930461818 | 0.775526925 | 0.836078863 | 0.821434996 | 0.876106796 | 1.147300694 | 1.368073688 | 0.642198987 | 1.077783956 | 1.239329972 | 1.181490833 | 1.104212471 |
| **ENSG00000048052** | HDAC9 | *954.4758456* | 0.308131758 | 0.199401982 | 0.292705289 | 0.341908983 | 0.856554649 | 1.422357601 | 1.844374522 | 1.230303044 | 1.177037412 | 0.846210391 | 1.979473534 | 1.501540835 |
| **ENSG00000090339** | ICAM1 | *3303.124062* | 0.61259522 | 0.733540627 | 0.625184565 | 0.668051633 | 1.096815039 | 0.944867129 | 0.806876432 | 1.55843933 | 1.197530997 | 1.200725075 | 1.361118206 | 1.194255747 |
| **ENSG00000157368** | IL34 | *154.2795177* | 0 | 0.054225647 | 0.011720841 | 0.006630961 | 0.575480636 | 0.812796537 | 0.854081522 | 0.442725557 | 2.089437184 | 2.256746522 | 2.407654538 | 2.488500055 |
| **ENSG00000115232** | ITGA4 | *2385.545359* | 0.194217655 | 0.245045738 | 0.300175012 | 0.22385541 | 0.522083762 | 0.974655657 | 0.892240793 | 0.920636346 | 1.902268325 | 1.994182238 | 2.241061622 | 1.589577442 |
| **ENSG00000091409** | ITGA6 | *3655.286829* | 0.613645153 | 0.417976417 | 0.826155957 | 0.671978641 | 0.55865715 | 1.184506051 | 1.300626702 | 0.743136504 | 1.342262869 | 1.406259552 | 1.693924572 | 1.240870432 |
| **ENSG00000138448** | ITGAV | *4331.328526* | 0.534609163 | 0.346943429 | 0.576553459 | 0.5158415 | 0.549753039 | 1.152626078 | 1.306925241 | 0.732378407 | 1.445742012 | 1.681158719 | 1.546420494 | 1.611048459 |
| **ENSG00000150093** | ITGB1 | *6127.826389* | 0.670613456 | 0.458547403 | 0.694356497 | 0.620040728 | 0.478331762 | 0.895143744 | 1.211054791 | 0.695366742 | 1.557968197 | 1.597408052 | 1.703119284 | 1.418049343 |
| **ENSG00000119185** | ITGB1BP1 | *3279.914475* | 0.803329606 | 0.811743658 | 0.733256887 | 0.688062295 | 1.380532528 | 1.398655869 | 1.541342481 | 0.784933878 | 0.901213094 | 0.972849671 | 1.001078027 | 0.983002007 |
| **ENSG00000138182** | KIF20B | *2326.43529* | 0.942510082 | 0.973622271 | 0.901641808 | 0.959507867 | 0.942426024 | 0.869906892 | 0.934545748 | 0.577596002 | 1.191392547 | 1.060464786 | 1.439356182 | 1.20702979 |
| **ENSG00000049130** | KITLG | *884.8381823* | 0.730785171 | 0.575556722 | 1.052471642 | 0.714511776 | 0.564413846 | 0.985139761 | 1.536821547 | 0.877330339 | 1.109200593 | 1.106263203 | 1.708208089 | 1.039297311 |
| **ENSG00000091136** | LAMB1 | *5887.794889* | 1.088669489 | 0.70671497 | 1.143424235 | 0.866332033 | 0.436257683 | 0.810060944 | 1.05796464 | 0.48411282 | 1.177127158 | 1.462123693 | 1.515991791 | 1.251220544 |
| **ENSG00000172061** | LRRC15 | *33.11137498* | 0 | 0 | 0.054612218 | 0 | 1.601392281 | 2.629967669 | 0.822433549 | 5.03809123 | 0.434622652 | 0.451723122 | 0.360092071 | 0.607065209 |
| **ENSG00000198909** | MAP3K3 | *3602.470078* | 0.471105393 | 0.654009164 | 0.472592659 | 0.591809733 | 1.318879499 | 0.96328671 | 0.742267773 | 1.814343493 | 1.255415569 | 1.375670055 | 1.019900922 | 1.32071903 |
| **ENSG00000071054** | MAP4K4 | *7669.261921* | 0.746875496 | 0.665546282 | 0.743425459 | 0.705645982 | 1.045602649 | 1.098108968 | 1.047367243 | 1.119603989 | 1.161269701 | 1.246442746 | 1.159302752 | 1.260808735 |
| **ENSG00000166974** | MAPRE2 | *1257.846643* | 0.778318086 | 0.740752342 | 0.819434396 | 0.831204664 | 0.980344678 | 1.143694343 | 1.302469255 | 1.071418615 | 0.990023063 | 1.26375913 | 1.00550444 | 1.073076988 |
| **ENSG00000076706** | MCAM | *10158.9692* | 0.61340742 | 0.517466532 | 0.639728145 | 0.529991003 | 0.941444338 | 1.734184626 | 2.512306536 | 0.885428772 | 0.764762252 | 0.94596055 | 1.021081442 | 0.894238384 |
| **ENSG00000087245** | MMP2 | *353.4520307* | 0.572775099 | 0.46746557 | 0.549977613 | 0.532564376 | 0.694271098 | 0.569126914 | 0.529377479 | 0.969950449 | 1.411468456 | 1.821999497 | 1.606229469 | 2.274793979 |
| **ENSG00000065534** | MYLK | *286.7957801* | 0.642683108 | 0.455785264 | 0.696717229 | 0.642073165 | 0.653547747 | 0.68925627 | 0.581965425 | 0.874782593 | 1.364852503 | 1.697860053 | 2.305737853 | 1.39473879 |
| **ENSG00000116044** | NFE2L2 | *4913.586326* | 0.890655477 | 0.844280258 | 0.90274686 | 0.940034807 | 1.106240327 | 1.005052347 | 0.971487872 | 1.097372552 | 1.095179142 | 1.024659081 | 1.038010451 | 1.084280826 |
| **ENSG00000164867** | NOS3 | *52.86462569* | 0.095262812 | 0.079125753 | 0.034205967 | 0.038703443 | 0.816411989 | 1.251918496 | 0.980399745 | 0.795104733 | 2.03259651 | 2.15343832 | 1.821677772 | 1.90115446 |
| **ENSG00000099250** | NRP1 | *138.973196* | 0.456591747 | 0.383762164 | 0.507458569 | 0.27972889 | 0.700975012 | 1.084033449 | 0.998716859 | 1.172006882 | 1.380693451 | 1.303474321 | 2.481442529 | 1.251116127 |
| **ENSG00000118257** | NRP2 | *4716.454802* | 0.975290261 | 0.617272001 | 1.075051727 | 0.8647993 | 0.337272183 | 0.71841134 | 0.829379617 | 0.3528584 | 1.388306296 | 1.790184555 | 1.695696913 | 1.355477406 |
| **ENSG00000153989** | NUS1 | *899.0738176* | 0.797633151 | 0.645536282 | 0.763279254 | 0.803330251 | 0.726920656 | 0.985038551 | 0.981944746 | 0.70273122 | 1.329599999 | 1.244942452 | 1.496518827 | 1.52252461 |
| **ENSG00000135124** | P2RX4 | *833.2736154* | 0.869080114 | 0.73541589 | 0.732408162 | 0.624906328 | 1.425099367 | 0.983398673 | 1.232375966 | 0.786597274 | 0.997076719 | 1.166743157 | 1.061052383 | 1.385845967 |
| **ENSG00000169313** | P2RY12 | *9.464533968* | 0.319257107 | 0 | 0.382118259 | 0 | 2.084621628 | 1.65615468 | 1.763480681 | 0.693921418 | 0.912308224 | 1.316949301 | 1.065959913 | 1.805228791 |
| **ENSG00000197461** | PDGFA | *716.2478931* | 0.008437357 | 0.029200458 | 0.007573993 | 0.009998146 | 2.017765745 | 2.470517709 | 2.656509873 | 1.252555183 | 0.755464499 | 0.765698171 | 1.288196292 | 0.738082573 |
| **ENSG00000100311** | PDGFB | *6.743077036* | 0.149368986 | 0 | 0.268169207 | 0 | 0.731489945 | 1.807996309 | 1.563289692 | 1.947966128 | 1.280509199 | 1.478768382 | 2.1762525 | 0.596189653 |
| **ENSG00000170962** | PDGFD | *223.2787986* | 0.279680866 | 0.266962581 | 0.307753598 | 0.274908732 | 0.938875325 | 1.704360951 | 2.258295376 | 1.076572111 | 0.872261825 | 1.566793065 | 0.84208086 | 1.61145471 |
| **ENSG00000124181** | PLCG1 | *7319.70751* | 0.849554903 | 0.617324571 | 0.848594169 | 0.801119356 | 0.658702817 | 1.235850219 | 1.645716746 | 0.670249349 | 1.102564109 | 1.320497847 | 1.168680159 | 1.081145755 |
| **ENSG00000162407** | PLPP3 | *542.9313982* | 0.558393161 | 0.556641919 | 0.57619326 | 0.663257968 | 1.015240784 | 1.121139988 | 1.465883914 | 1.062084533 | 1.309397672 | 1.245823384 | 1.400415775 | 1.025527642 |
| **ENSG00000128567** | PODXL | *96.24387023* | 0.125581805 | 0.130386069 | 0.056365739 | 0.10629472 | 0.960936468 | 1.058620764 | 1.32346135 | 1.282904406 | 2.193046884 | 1.769937022 | 1.791563443 | 1.20090133 |
| **ENSG00000100034** | PPM1F | *4368.451934* | 0.681315825 | 0.80289457 | 0.686936707 | 0.730654062 | 1.371877074 | 0.984353635 | 0.844775791 | 1.90604308 | 1.049561239 | 0.942335326 | 1.001680632 | 0.997572059 |
| **ENSG00000154229** | PRKCA | *233.7571243* | 0.590302013 | 0.420519386 | 0.50669133 | 0.428890069 | 0.722706901 | 0.704084159 | 0.605030627 | 0.651827465 | 1.543196762 | 1.97290039 | 2.138351364 | 1.715499533 |
| **ENSG00000136238** | RAC1 | *11787.13427* | 0.861417989 | 0.95949191 | 0.814156483 | 0.760118812 | 1.080265187 | 0.991526374 | 0.933438787 | 0.826977556 | 1.125673037 | 1.152903066 | 1.225362532 | 1.268668267 |
| **ENSG00000155366** | RHOC | *2139.545767* | 0.758860608 | 0.767852192 | 0.798688179 | 0.647891803 | 0.962501425 | 1.134339389 | 1.41813342 | 0.634188354 | 1.206675323 | 1.192711116 | 1.458344574 | 1.019813618 |
| **ENSG00000126785** | RHOJ | *976.598208* | 0.267117532 | 0.247354082 | 0.279594132 | 0.226267719 | 0.43688453 | 0.825700769 | 0.82663907 | 0.330871002 | 1.76633095 | 2.344986364 | 2.337525704 | 2.110728147 |
| **ENSG00000132669** | RIN2 | *413.724044* | 0.87641599 | 0.75070397 | 0.841369959 | 0.853086569 | 0.625914517 | 0.879818614 | 0.94273265 | 0.758798846 | 1.226713728 | 1.325593011 | 1.507459223 | 1.411392923 |
| **ENSG00000134318** | ROCK2 | *3904.200341* | 0.555689457 | 0.376864324 | 0.684788192 | 0.57856445 | 0.918793929 | 1.611957491 | 1.930507674 | 1.124718409 | 0.936985833 | 1.052686488 | 1.317417207 | 0.911026546 |
| **ENSG00000108443** | RPS6KB1 | *1242.030478* | 0.754169993 | 0.66262147 | 0.761441363 | 0.837670981 | 0.654273985 | 0.741088987 | 0.976737282 | 0.837591865 | 1.392713509 | 1.359466412 | 1.538170361 | 1.484053791 |
| **ENSG00000126458** | RRAS | *413.0540146* | 0.872960779 | 0.724072761 | 0.833979089 | 0.577077105 | 0.913526331 | 1.123693805 | 1.193087658 | 0.508806508 | 1.184572491 | 1.291532122 | 1.620926917 | 1.155764435 |
| **ENSG00000115310** | RTN4 | *8949.093601* | 0.880804137 | 0.756396622 | 0.837250319 | 0.835418828 | 1.040062246 | 1.167793384 | 1.043938185 | 1.054451952 | 1.137241698 | 1.112384168 | 1.118337089 | 1.015921373 |
| **ENSG00000111961** | SASH1 | *2868.573204* | 0.675901405 | 0.432356284 | 0.780092853 | 0.707198853 | 0.704132599 | 1.258609443 | 1.592405782 | 0.793088171 | 1.156865685 | 1.471259591 | 1.242465335 | 1.185623999 |
| **ENSG00000073060** | SCARB1 | *10782.19487* | 0.973372551 | 0.847961874 | 0.826225015 | 0.690256638 | 0.829043682 | 1.194264496 | 1.130670461 | 0.551862072 | 1.11028921 | 1.454258983 | 1.009356511 | 1.382438508 |
| **ENSG00000075213** | SEMA3A | *386.9659107* | 0.564814163 | 0.502647191 | 0.894876491 | 0.674143332 | 0.31229128 | 1.003667302 | 1.314383792 | 0.882551713 | 1.214847282 | 1.292709125 | 1.974329865 | 1.368738463 |
| **ENSG00000012171** | SEMA3B | *2308.493638* | 0.355588314 | 0.287652443 | 0.45510801 | 0.366046397 | 1.483918503 | 1.943079156 | 2.286978751 | 0.795459387 | 0.906826867 | 1.161215201 | 1.114824248 | 0.843302724 |
| **ENSG00000168758** | SEMA4C | *3232.814514* | 1.087646118 | 0.901527971 | 0.893287296 | 0.883209679 | 0.501211548 | 0.954911399 | 1.207019 | 0.426220391 | 1.176686359 | 1.494672221 | 1.28291296 | 1.190695058 |
| **ENSG00000138623** | SEMA7A | *1262.897866* | 0.782382866 | 0.70549573 | 0.713779322 | 0.694220387 | 1.155109102 | 1.634899081 | 1.756343641 | 0.876796621 | 0.85691784 | 1.050784646 | 0.903440536 | 0.86983023 |
| **ENSG00000117090** | SLAMF1 | *32.67992368* | 0.246562773 | 0.639988229 | 0.359665973 | 0.375651375 | 1.320667533 | 1.172463328 | 0.403205609 | 3.657630926 | 0.822006614 | 0.940800895 | 0.954212947 | 1.107143798 |
| **ENSG00000166949** | SMAD3 | *2880.409819* | 0.495838792 | 0.393911336 | 0.558103193 | 0.44502208 | 0.839517593 | 1.518878285 | 1.906694115 | 1.249498681 | 0.889980488 | 1.082971473 | 1.541764082 | 1.07781988 |
| **ENSG00000124216** | SNAI1 | *139.6357131* | 0.180327538 | 0.157269973 | 0.220150382 | 0.307707126 | 0.732973167 | 1.415653526 | 1.283364435 | 0.950089216 | 1.710807534 | 1.779310381 | 1.858820057 | 1.403526666 |
| **ENSG00000198648** | STK39 | *925.955149* | 0.908270225 | 0.678746409 | 0.911998152 | 0.817572998 | 0.729788639 | 1.229173707 | 1.6251137 | 0.699353078 | 1.010213061 | 1.147777283 | 1.148987912 | 1.093004837 |
| **ENSG00000100242** | SUN2 | *3945.199091* | 0.670415969 | 0.73741374 | 0.692568236 | 0.712059655 | 1.167735367 | 0.989967981 | 0.802030942 | 1.49291885 | 1.319499346 | 1.130840909 | 1.276759288 | 1.007789717 |
| **ENSG00000054654** | SYNE2 | *2752.622561* | 0.828781587 | 0.555802742 | 0.751530115 | 0.671206033 | 0.836828953 | 0.919657235 | 1.207274467 | 1.279828068 | 1.115674029 | 1.460482073 | 1.364107494 | 1.008827202 |
| **ENSG00000106799** | TGFBR1 | *3134.25773* | 0.866692011 | 0.733024306 | 0.934069458 | 0.963859011 | 0.862013689 | 0.789340076 | 0.850909556 | 1.222478544 | 1.233583793 | 1.123313012 | 1.211178471 | 1.209538071 |
| **ENSG00000137801** | THBS1 | *3521.717234* | 0.049763775 | 0.105413661 | 0.051603435 | 0.07262235 | 0.709400433 | 2.129744914 | 2.235958326 | 1.452382431 | 0.833886919 | 0.889300807 | 2.970217337 | 0.499705613 |
| **ENSG00000156299** | TIAM1 | *810.2562211* | 0.745843019 | 0.508506925 | 0.842483901 | 0.75881668 | 0.637673163 | 1.300440904 | 1.52107853 | 0.852713016 | 1.129599041 | 1.391664359 | 1.446624784 | 0.86455568 |
| **ENSG00000104067** | TJP1 | *730.5117534* | 0.699035617 | 0.455221679 | 0.847813635 | 0.625986671 | 0.562112854 | 1.207563181 | 1.630605194 | 0.773180282 | 1.222702465 | 1.450307218 | 1.412448873 | 1.113022332 |
| **ENSG00000111199** | TRPV4 | *561.5686177* | 0.096852198 | 0.046554343 | 0.096601852 | 0.076512295 | 1.370213526 | 2.112038851 | 3.217716676 | 1.050226647 | 0.984051818 | 1.065385294 | 1.053426663 | 0.830419837 |
| **ENSG00000100219** | XBP1 | *2962.345817* | 0.772486901 | 0.712374607 | 0.782257767 | 0.628521883 | 1.032339193 | 1.395733749 | 0.984803219 | 0.80434256 | 1.24234197 | 1.321180585 | 1.395711231 | 0.927906334 |
