## Supplementary material for "Peripheral positioning of lysosomes supports melanoma aggressiveness": Table 3

**Table 3**: Proteomic analysis of cell medium, list of hits corresponding to Figure 2D)

| protein_set_id | accession | Clustered Vs Spread log Fold Change | Clustered Vs Spread pValue |
|---|---|---|---|
| 19314 | sp\|P04908\|H2A1B_HUMAN | 6,342191451 | 0,000204616 |
| 19395 | sp\|P53634\|CATC_HUMAN | -0,781580636 | 0,004082027 |
| 20188 | sp\|Q96KG7\|MEG10_HUMAN | -1,351320057 | 0,004085503 |
| 19592 | sp\|Q07954\|LRP1_HUMAN | -1,102509205 | 0,004949343 |
| 20455 | sp\|P06280\|AGAL_HUMAN | -0,823936279 | 0,005930019 |
| 20320 | sp\|P98155\|VLDLR_HUMAN | -1,642686833 | 0,010427447 |
| 20016 | sp\|P01130\|LDLR_HUMAN | -0,799216732 | 0,010513742 |
| 20193 | sp\|P02786\|TFR1_HUMAN | -0,954069546 | 0,012939708 |
| 20447 | sp\|Q99519\|NEUR1_HUMAN | -0,960603563 | 0,014698715 |
| 19462 | sp\|P53041\|PPP5_HUMAN | 0,660528097 | 0,01525975 |
| 19856 | sp\|P54578\|UBP14_HUMAN | 0,770161562 | 0,017425941 |
| 20487 | sp\|O75054\|IGSF3_HUMAN | -0,587333651 | 0,024072151 |
| 20291 | sp\|Q5VU97\|CAHD1_HUMAN | 1,024205139 | 0,02457342 |
| 19904 | sp\|P24752\|THIL_HUMAN | -1,508107986 | 0,024917899 |
| 19862 | sp\|P25774\|CATS_HUMAN | -1,121522126 | 0,025834738 |
| 19874 | sp\|Q68BL7\|OLM2A_HUMAN | 1,209553813 | 0,026552665 |
| 19808 | sp\|P28062\|PSB8_HUMAN | 0,561182673 | 0,02768074 |
| 20068 | sp\|Q92484\|ASM3A_HUMAN | -0,898032715 | 0,031112035 |
| 19901 | sp\|Q8NHP8\|PLBL2_HUMAN | -0,795350697 | 0,031819803 |
| 19650 | sp\|O43237\|DC1L2_HUMAN | -0,539174655 | 0,033622041 |
| 20492 | sp\|P00390\|GSHR_HUMAN | 1,121204278 | 0,033653576 |
| 19764 | sp\|P10155\|RO60_HUMAN | 0,962255171 | 0,041429745 |
| 20477 | sp\|P30048\|PRDX3_HUMAN | -1,325967988 | 0,046757674 |
| 19403 | sp\|Q13510\|ASAH1_HUMAN | -0,861434931 | 0,048969694 |
| 19496 | sp\|P43235\|CATK_HUMAN | -1,246452127 | 0,050948014 |
| 19974 | sp\|P61163\|ACTZ_HUMAN | 0,637810883 | 0,051177857 |
| 20267 | sp\|O00462\|MANBA_HUMAN | -0,840116856 | 0,055797238 |
| 19479 | sp\|O00584\|RNT2_HUMAN | -0,426035275 | 0,056138492 |
| 19729 | sp\|P13686\|PPA5_HUMAN | -0,917029322 | 0,056954602 |
| 20111 | sp\|O14979\|HNRDL_HUMAN | -1,103315758 | 0,058053677 |
| 19770 | sp\|P07686\|HEXB_HUMAN | -0,578173086 | 0,060057885 |
| 20031 | sp\|O95834\|EMAL2_HUMAN | 0,664530063 | 0,066556922 |
| 19680 | sp\|P62263\|RS14_HUMAN | 1,692340296 | 0,067359269 |
| 19785 | sp\|Q92820\|GGH_HUMAN | -0,53056449 | 0,080189891 |
| 19887 | sp\|Q13332\|PTPRS_HUMAN | -0,693339196 | 0,086579212 |
| 20268 | sp\|Q14956\|GPNMB_HUMAN | -0,509313945 | 0,087543759 |
| 19685 | sp\|Q00688\|FKBP3_HUMAN | 0,496418166 | 0,087682207 |
| 20029 | sp\|P07339\|CATD_HUMAN | -0,474418939 | 0,0905938 |
| 19504 | sp\|Q13740\|CD166_HUMAN | 0,960462512 | 0,093384 |
| 19916 | sp\|Q9Y646\|CBPQ_HUMAN | -0,775399816 | 0,094158689 |
| 19568 | sp\|O15031\|PLXB2_HUMAN | -0,641130383 | 0,09484923 |
| 19594 | sp\|P52888\|THOP1_HUMAN | 0,637383658 | 0,09906184 |
| 19360 | sp\|P00505\|AATM_HUMAN | 0,604862976 | 0,101569501 |
| 20198 | sp\|P23396\|RS3_HUMAN | 0,495583805 | 0,101653404 |
| 19918 | sp\|O75828\|CBR3_HUMAN | 0,366402824 | 0,101719989 |
| 20452 | sp\|O00469\|PLOD2_HUMAN | -0,406758182 | 0,102630827 |
| 19636 | sp\|P67809\|YBOX1_HUMAN | 0,577354666 | 0,104428002 |
| 19381 | sp\|P07858\|CATB_HUMAN | -0,641357707 | 0,104730356 |
| 20191 | sp\|Q16719\|KYNU_HUMAN | 0,58085926 | 0,109296523 |
| 20363 | sp\|Q9H1B5\|XYLT2_HUMAN | -0,462439383 | 0,112307675 |
| 20394 | sp\|Q07021\|C1QBP_HUMAN | 0,564248429 | 0,114980943 |
| 19580 | sp\|P38571\|LICH_HUMAN | -0,813616093 | 0,116565321 |
| 19996 | sp\|P08581\|MET_HUMAN | -1,01237578 | 0,117613183 |
| 20204 | sp\|Q9GZZ1\|NAA50_HUMAN | 0,584853366 | 0,117811787 |
| 19669 | sp\|P60900\|PSA6_HUMAN | 0,51233974 | 0,122745949 |
| 19753 | sp\|P05556\|ITB1_HUMAN | -0,318060155 | 0,126986356 |
| 20396 | sp\|P51580\|TPMT_HUMAN | 0,332468195 | 0,131667075 |
| 20485 | sp\|O95831\|AIFM1_HUMAN | 0,603768164 | 0,133724764 |
| 20212 | sp\|P11279\|LAMP1_HUMAN | -0,471238093 | 0,136141487 |
| 19440 | sp\|P07602\|SAP_HUMAN | -0,845667279 | 0,138931415 |
| 20048 | sp\|Q5JRX3\|PREP_HUMAN | 0,53407336 | 0,140466216 |
| 20073 | sp\|O00115\|DNS2A_HUMAN | -0,593579123 | 0,145879496 |
| 20345 | sp\|Q8N0V5\|GNT2A_HUMAN | -0,984637537 | 0,14652591 |
| 19749 | sp\|O00560\|SDCB1_HUMAN | -0,386052389 | 0,149287619 |
| 19551 | sp\|Q9NSE4\|SYIM_HUMAN | 0,765166743 | 0,149405932 |
| 20127 | sp\|P42785\|PCP_HUMAN | -0,948774743 | 0,149809019 |
| 19494 | sp\|P15586\|GNS_HUMAN | -0,469025844 | 0,152350736 |
| 19524 | sp\|Q9UHL4\|DPP2_HUMAN | -0,875986087 | 0,153997026 |
| 19955 | sp\|P11717\|MPRI_HUMAN | -0,434947732 | 0,155188993 |
| 20229 | sp\|P20042\|IF2B_HUMAN | 0,559429653 | 0,1608213 |
| 20223 | sp\|Q9UBG0\|MRC2_HUMAN | -0,467522369 | 0,161186314 |
| 19882 | sp\|Q9UBR2\|CATZ_HUMAN | -0,764769746 | 0,162214039 |
| 19418 | sp\|Q99733\|NP1L4_HUMAN | -0,688828784 | 0,162573606 |
| 20202 | sp\|Q8IZQ5\|SELH_HUMAN | -0,497119433 | 0,16338948 |
| 19979 | sp\|O95166\|GBRAP_HUMAN | 0,352088108 | 0,164250139 |
| 19829 | sp\|Q00577\|PURA_HUMAN | 0,324374895 | 0,169310137 |

| | | | |
|---|---|---|---|
| 19477 | sp\|A0A0B4J2D5\|GAL3B_HUMAN | 1,004289476 | 0,169671131 |
| 20232 | sp\|P10619\|PPGB_HUMAN | -0,623265608 | 0,170424446 |
| 19844 | sp\|Q9NXG2\|THUM1_HUMAN | 0,793238817 | 0,170971405 |
| 19626 | sp\|P62820\|RAB1A_HUMAN | -0,410350392 | 0,172315869 |
| 19796 | sp\|P06276\|CHLE_HUMAN | -0,325520478 | 0,175348472 |
| 19899 | sp\|Q8NCW5\|NNRE_HUMAN | 0,447412713 | 0,177393391 |
| 20261 | sp\|P12955\|PEPD_HUMAN | 0,499488701 | 0,17998212 |
| 20051 | sp\|P13798\|ACPH_HUMAN | 1,079106286 | 0,180941087 |
| 19647 | sp\|Q9BTY2\|FUCO2_HUMAN | -0,469939138 | 0,191530237 |
| 19526 | sp\|P48723\|HSP13_HUMAN | -0,504639277 | 0,193047605 |
| 20182 | sp\|P31150\|GDIA_HUMAN | 0,325726543 | 0,194609506 |
| 20005 | sp\|P00750\|TPA_HUMAN | -0,540550672 | 0,196408905 |
| 19880 | sp\|P07996\|TSP1_HUMAN | 0,531063323 | 0,197405966 |
| 19489 | sp\|P13796\|PLSL_HUMAN | 0,502689524 | 0,199807522 |
| 19845 | sp\|P25789\|PSA4_HUMAN | 0,380824277 | 0,200913619 |
| 19365 | sp\|Q08431\|MFGM_HUMAN | -0,76248342 | 0,203396907 |
| 20258 | sp\|P07585\|PGS2_HUMAN | -0,660025086 | 0,204794629 |
| 19715 | sp\|P40306\|PSB10_HUMAN | 0,492184997 | 0,205226774 |
| 20177 | sp\|P18206\|VINC_HUMAN | 0,33217295 | 0,208752527 |
| 20315 | sp\|P29401\|TKT_HUMAN | 0,29848633 | 0,210753982 |
| 20311 | sp\|O96019\|ACL6A_HUMAN | 0,715911253 | 0,212629013 |
| 19835 | sp\|Q99436\|PSB7_HUMAN | 0,376456244 | 0,215424742 |
| 19633 | sp\|P37802\|TAGL2_HUMAN | -0,401612701 | 0,215479609 |
| 20393 | sp\|P43007\|SATT_HUMAN | -0,362918302 | 0,221404547 |
| 19991 | sp\|O00410\|IPO5_HUMAN | 0,711052926 | 0,222191197 |
| 20094 | sp\|P06865\|HEXA_HUMAN | -0,65148129 | 0,22489233 |
| 20422 | sp\|P06858\|LIPL_HUMAN | -1,216557785 | 0,225318257 |
| 19967 | sp\|Q14240\|IF4A2_HUMAN | 0,787257513 | 0,23097056 |
| 19857 | sp\|P40926\|MDHM_HUMAN | 0,470975023 | 0,232535684 |
| 20163 | sp\|P41271\|NBL1_HUMAN | -0,477395541 | 0,232970055 |
| 19570 | sp\|Q9UHD1\|CHRD1_HUMAN | 0,335863415 | 0,234447478 |
| 19548 | sp\|Q9GZL7\|WDR12_HUMAN | 0,633702155 | 0,235440771 |
| 20161 | sp\|Q7Z739\|YTHD3_HUMAN | -0,506569863 | 0,235952264 |
| 20331 | sp\|Q14103\|HNRPD_HUMAN | 0,857458699 | 0,236685532 |
| 20361 | sp\|P84090\|ERH_HUMAN | 0,683821421 | 0,238653728 |
| 19926 | sp\|P41091\|IF2G_HUMAN | 0,378368966 | 0,238863433 |
| 19693 | sp\|Q92598\|HS105_HUMAN | 0,333712968 | 0,239281462 |
| 19742 | sp\|P52907\|CAZA1_HUMAN | 0,298947829 | 0,240038359 |
| 20080 | sp\|Q9NQR4\|NIT2_HUMAN | -0,756872854 | 0,24021154 |
| 19696 | sp\|P12273\|PIP_HUMAN | -0,846677275 | 0,242244284 |
| 20066 | sp\|Q96G03\|PGM2_HUMAN | 1,056244984 | 0,245086628 |
| 20276 | sp\|Q06481\|APLP2_HUMAN | -0,399594017 | 0,249748529 |
| 20178 | sp\|Q9NPH2\|INO1_HUMAN | 0,46110533 | 0,24976896 |
| 19376 | sp\|Q02790\|FKBP4_HUMAN | 0,247093463 | 0,25093351 |
| 20356 | sp\|P07711\|CATL1_HUMAN | -0,267649734 | 0,253534499 |
| 20460 | sp\|P07311\|ACYP1_HUMAN | -0,515183466 | 0,254661747 |
| 19851 | sp\|P36957\|ODO2_HUMAN | -0,423040511 | 0,258802263 |
| 20471 | sp\|P62241\|RS8_HUMAN | 0,284921894 | 0,259191643 |
| 20116 | sp\|P04083\|ANXA1_HUMAN | -0,388432066 | 0,260945237 |
| 20303 | sp\|P02100\|HBE_HUMAN | -0,522487496 | 0,263374959 |
| 20289 | sp\|P28066\|PSA5_HUMAN | 0,27220948 | 0,263952296 |
| 20036 | sp\|O14773\|TPP1_HUMAN | -0,735515739 | 0,264531862 |
| 19363 | sp\|Q9NR28\|DBLOH_HUMAN | -0,781042887 | 0,265844898 |
| 19488 | sp\|Q32MZ4\|LRRF1_HUMAN | 0,537467186 | 0,266544076 |
| 20137 | sp\|P23284\|PPIB_HUMAN | -0,365385521 | 0,270376025 |
| 19561 | sp\|Q02809\|PLOD1_HUMAN | -0,327214183 | 0,271353944 |
| 20466 | sp\|P30740\|ILEU_HUMAN | 0,293917016 | 0,273626305 |
| 20364 | sp\|P17931\|LEG3_HUMAN | -0,383901954 | 0,277633438 |
| 19919 | sp\|Q9BY44\|EIF2A_HUMAN | 0,313042035 | 0,279149419 |
| 19826 | sp\|Q16658\|FSCN1_HUMAN | -0,237963485 | 0,281666807 |
| 19413 | sp\|P82970\|HMGN5_HUMAN | 0,295511673 | 0,283802561 |
| 19574 | sp\|P13987\|CD59_HUMAN | -0,360296483 | 0,283868638 |
| 20486 | sp\|Q14974\|IMB1_HUMAN | 0,226410018 | 0,28496871 |
| 20446 | sp\|O00499\|BIN1_HUMAN | 0,269174544 | 0,286274895 |
| 19303 | sp\|P62081\|RS7_HUMAN | 0,251613728 | 0,286686275 |
| 19797 | sp\|P25786\|PSA1_HUMAN | 0,378418414 | 0,296182902 |
| 20334 | sp\|Q8WUM4\|PDC6I_HUMAN | -0,338845939 | 0,296606239 |
| 19550 | sp\|Q13011\|ECH1_HUMAN | 0,359272255 | 0,299190854 |
| 20184 | sp\|O15240\|VGF_HUMAN | -0,805603238 | 0,29976068 |
| 19831 | sp\|A6NMY6\|AXA2L_HUMAN | -0,402825138 | 0,300793725 |
| 20386 | sp\|P61916\|NPC2_HUMAN | -0,469919617 | 0,301020064 |
| 20427 | sp\|P25787\|PSA2_HUMAN | 0,317971201 | 0,301220171 |
| 20298 | sp\|Q9BTT0\|AN32E_HUMAN | 0,270103221 | 0,302961979 |
| 19834 | sp\|P46782\|RS5_HUMAN | 0,218009863 | 0,303499236 |
| 19728 | sp\|P14324\|FPPS_HUMAN | 0,24624998 | 0,304138138 |
| 19353 | sp\|P41250\|GARS_HUMAN | -0,348846905 | 0,305471437 |
| 19921 | sp\|Q9H4A4\|AMPB_HUMAN | 0,278740279 | 0,306151176 |
| 20366 | sp\|Q02750\|MP2K1_HUMAN | 1,119124164 | 0,307540369 |

| | | | |
|---|---|---|---|
| 20310 | sp\|P15104\|GLNA_HUMAN | 0,527444117 | 0,307732418 |
| 19840 | sp\|P43034\|LIS1_HUMAN | 0,280767194 | 0,308098792 |
| 19690 | sp\|P00491\|PNPH_HUMAN | 0,25614642 | 0,308308433 |
| 19430 | sp\|P04075\|ALDOA_HUMAN | 0,262915447 | 0,308751646 |
| 19983 | sp\|Q02880\|TOP2B_HUMAN | 0,315303863 | 0,309451844 |
| 19535 | sp\|O43324\|MCA3_HUMAN | 0,389831133 | 0,309954394 |
| 19646 | sp\|P50281\|MMP14_HUMAN | -0,318451573 | 0,310412973 |
| 20153 | sp\|O15143\|ARC1B_HUMAN | 0,238616777 | 0,311550014 |
| 20426 | sp\|Q9HC38\|GLOD4_HUMAN | 0,290828233 | 0,315700332 |
| 20170 | sp\|P61247\|RS3A_HUMAN | 0,33203745 | 0,319114313 |
| 20207 | sp\|P62495\|ERF1_HUMAN | 0,774176785 | 0,320257122 |
| 19827 | sp\|P62826\|RAN_HUMAN | 0,252786338 | 0,324009143 |
| 20101 | sp\|P09622\|DLDH_HUMAN | 0,313579981 | 0,325249625 |
| 20279 | sp\|Q04446\|GLGB_HUMAN | 0,459706233 | 0,328272099 |
| 20290 | sp\|P27105\|STOM_HUMAN | -0,426220807 | 0,328548555 |
| 19861 | sp\|P14543\|NID1_HUMAN | -0,405712727 | 0,334091107 |
| 20371 | sp\|Q8WVY7\|UBCP1_HUMAN | 0,80531858 | 0,340428516 |
| 19746 | sp\|P49746\|TSP3_HUMAN | -0,253277127 | 0,340719915 |
| 20392 | sp\|Q9NPJ3\|ACO13_HUMAN | 0,453306505 | 0,340799762 |
| 20179 | sp\|Q15819\|UB2V2_HUMAN | 0,593043169 | 0,342293536 |
| 20375 | sp\|Q96JB1\|DYH8_HUMAN | -0,859712674 | 0,342746166 |
| 20012 | sp\|P42677\|RS27_HUMAN | 0,567640039 | 0,343879813 |
| 19676 | sp\|P06733\|ENOA_HUMAN | 0,371027832 | 0,345682624 |
| 19814 | sp\|Q9Y617\|SERC_HUMAN | 0,262086052 | 0,345753093 |
| 19295 | sp\|P05023\|AT1A1_HUMAN | -0,374863793 | 0,345934339 |
| 20217 | sp\|Q9UJ70\|NAGK_HUMAN | -0,456503319 | 0,347011457 |
| 19932 | sp\|P07954\|FUMH_HUMAN | 0,29177074 | 0,347733271 |
| 19755 | sp\|Q14152\|EIF3A_HUMAN | -0,470343335 | 0,351307419 |
| 19471 | sp\|P12109\|CO6A1_HUMAN | -0,58826246 | 0,352691549 |
| 19708 | sp\|Q9GZT8\|NIF3L_HUMAN | 0,299702551 | 0,352695102 |
| 19930 | sp\|P61224\|RAP1B_HUMAN | -0,303869904 | 0,353147389 |
| 19817 | sp\|P10451\|OSTP_HUMAN | -0,343967625 | 0,353224253 |
| 19460 | sp\|P36551\|HEM6_HUMAN | 0,320729472 | 0,353585017 |
| 19359 | sp\|Q9Y2S6\|TMA7_HUMAN | 0,271430675 | 0,353869311 |
| 19947 | sp\|Q09328\|MGT5A_HUMAN | -0,634912753 | 0,354910654 |
| 20272 | sp\|P19021\|AMD_HUMAN | -0,68111263 | 0,358737672 |
| 20445 | sp\|O00625\|PIR_HUMAN | -0,232570958 | 0,359674582 |
| 20135 | sp\|P55786\|PSA_HUMAN | 0,256557237 | 0,360984958 |
| 19705 | sp\|P30084\|ECHM_HUMAN | 0,704511585 | 0,361505528 |
| 20186 | sp\|Q13214\|SEM3B_HUMAN | -0,386984986 | 0,362299301 |
| 20023 | sp\|P52292\|IMA1_HUMAN | 0,319994707 | 0,362695104 |
| 19394 | sp\|P09871\|C1S_HUMAN | -0,275109807 | 0,363835005 |
| 19497 | sp\|O60911\|CATL2_HUMAN | -0,283414952 | 0,366933387 |
| 20401 | sp\|P49207\|RL34_HUMAN | -0,946935444 | 0,36737537 |
| 19564 | sp\|P28838\|AMPL_HUMAN | 0,703287294 | 0,369439243 |
| 19928 | sp\|P17174\|AATC_HUMAN | 0,622082151 | 0,372152457 |
| 20297 | sp\|Q53H82\|LACB2_HUMAN | 0,244433984 | 0,373232733 |
| 19317 | sp\|Q16610\|ECM1_HUMAN | -0,407886656 | 0,376249239 |
| 19900 | sp\|Q16363\|LAMA4_HUMAN | -0,348789302 | 0,377927834 |
| 20389 | sp\|P51884\|LUM_HUMAN | -0,340810966 | 0,378678368 |
| 20349 | sp\|P20827\|EFNA1_HUMAN | -0,236023884 | 0,385923324 |
| 19630 | sp\|P00736\|C1R_HUMAN | -0,312529355 | 0,386240806 |
| 20360 | sp\|P32004\|L1CAM_HUMAN | -0,282876782 | 0,389028702 |
| 19684 | sp\|O43396\|TXNL1_HUMAN | 0,221146795 | 0,393126241 |
| 20167 | sp\|P30153\|2AAA_HUMAN | 0,23241594 | 0,394235873 |
| 20155 | sp\|Q99598\|TSNAX_HUMAN | 0,528885425 | 0,395147505 |
| 20123 | sp\|P62750\|RL23A_HUMAN | 0,354708813 | 0,395265109 |
| 19385 | sp\|P09341\|GROA_HUMAN | -0,333489397 | 0,397373594 |
| 19800 | sp\|Q12841\|FSTL1_HUMAN | -0,297622112 | 0,398547726 |
| 20026 | sp\|P78504\|JAG1_HUMAN | -0,568500212 | 0,399331957 |
| 20417 | sp\|P50552\|VASP_HUMAN | 0,211669817 | 0,399912075 |
| 19389 | sp\|P30566\|PUR8_HUMAN | 0,200888755 | 0,400832882 |
| 20254 | sp\|Q9NRG0\|CHRC1_HUMAN | 0,281690816 | 0,40283903 |
| 20302 | sp\|O95782\|AP2A1_HUMAN | -0,320154899 | 0,405151966 |
| 20156 | sp\|P61158\|ARP3_HUMAN | 0,210183748 | 0,405505536 |
| 19370 | sp\|Q13685\|AAMP_HUMAN | 0,215095992 | 0,406782972 |
| 20200 | sp\|P05067\|A4_HUMAN | -0,342046781 | 0,406847239 |
| 19472 | sp\|P63313\|TYB10_HUMAN | 0,294349918 | 0,40688756 |
| 20425 | sp\|Q15758\|AAAT_HUMAN | -0,236383955 | 0,407765019 |
| 20183 | sp\|P50395\|GDIB_HUMAN | 0,195756879 | 0,411103044 |
| 20173 | sp\|P31949\|S10AB_HUMAN | 0,179666919 | 0,414650389 |
| 19436 | sp\|O00533\|NCHL1_HUMAN | -0,455595002 | 0,414910195 |
| 19556 | sp\|Q8NBJ4\|GOLM1_HUMAN | -0,427709133 | 0,415813187 |
| 20374 | sp\|P04062\|GLCM_HUMAN | -0,47242389 | 0,416016098 |
| 20072 | sp\|P62888\|RL30_HUMAN | 0,191345155 | 0,417156152 |
| 20064 | sp\|P34932\|HSP74_HUMAN | 0,173128513 | 0,418042313 |
| 19956 | sp\|Q8N475\|FSTL5_HUMAN | -0,487444587 | 0,419435088 |
| 20476 | sp\|Q9NZU0\|FLRT3_HUMAN | -0,205304703 | 0,419882512 |

| | | | |
|---|---|---|---|
| 19923 | sp\|P46777\|RL5_HUMAN | 0,320362163 | 0,421235073 |
| 19279 | sp\|P30050\|RL12_HUMAN | 0,228321055 | 0,421370372 |
| 19954 | sp\|P49773\|HINT1_HUMAN | -0,469164132 | 0,422703697 |
| 20097 | sp\|P84098\|RL19_HUMAN | 0,221868179 | 0,422990964 |
| 19740 | sp\|O43175\|SERA_HUMAN | 0,63075972 | 0,423020031 |
| 19571 | sp\|P13693\|TCTP_HUMAN | 0,22181904 | 0,423035971 |
| 20475 | sp\|Q14008\|CKAP5_HUMAN | 0,39995356 | 0,423854754 |
| 20018 | sp\|O60568\|PLOD3_HUMAN | -0,31227536 | 0,424815451 |
| 20493 | sp\|Q9Y376\|CAB39_HUMAN | 0,295693059 | 0,430223795 |
| 20192 | sp\|Q9UKZ9\|PCOC2_HUMAN | -0,262973503 | 0,431501324 |
| 19841 | sp\|P61201\|CSN2_HUMAN | 0,220429746 | 0,432338099 |
| 19396 | sp\|O95373\|IPO7_HUMAN | 0,21539174 | 0,432782245 |
| 19456 | sp\|P49458\|SRP09_HUMAN | 0,232734567 | 0,434065319 |
| 20019 | sp\|Q9BXS5\|AP1M1_HUMAN | 0,43388456 | 0,434352622 |
| 20353 | sp\|P62269\|RS18_HUMAN | 0,229809677 | 0,436467691 |
| 20037 | sp\|P06756\|ITAV_HUMAN | -0,296989736 | 0,437687964 |
| 20317 | sp\|P30520\|PURA2_HUMAN | 0,28157813 | 0,439983376 |
| 19518 | sp\|O75347\|TBCA_HUMAN | 1,331816615 | 0,442251909 |
| 19536 | sp\|O76003\|GLRX3_HUMAN | -0,447766238 | 0,44289622 |
| 19508 | sp\|O75083\|WDR1_HUMAN | -0,265053443 | 0,444779343 |
| 19323 | sp\|P11021\|BIP_HUMAN | 0,185227315 | 0,447309911 |
| 19422 | sp\|O95336\|6PGL_HUMAN | -0,459658815 | 0,447343865 |
| 19612 | sp\|P36871\|PGM1_HUMAN | 0,190119432 | 0,449367034 |
| 19474 | sp\|Q16531\|DDB1_HUMAN | 0,260629774 | 0,450437478 |
| 20464 | sp\|P48681\|NEST_HUMAN | -0,469302568 | 0,45176519 |
| 19725 | sp\|Q92499\|DDX1_HUMAN | 0,644099121 | 0,451935044 |
| 19623 | sp\|P62273\|RS29_HUMAN | 0,253839865 | 0,46242427 |
| 19585 | sp\|Q08380\|LG3BP_HUMAN | -0,243549845 | 0,465153771 |
| 20352 | sp\|Q9H1E3\|NUCKS_HUMAN | 0,294631736 | 0,466078211 |
| 19619 | sp\|A0A1B0GUS4\|UB2L5_HUMAN | 0,261055128 | 0,466078403 |
| 20039 | sp\|P10809\|CH60_HUMAN | 0,186010065 | 0,466910752 |
| 20318 | sp\|O43278\|SPIT1_HUMAN | -0,191231956 | 0,467759962 |
| 19498 | sp\|P00367\|DHE3_HUMAN | -0,294946432 | 0,471540102 |
| 20122 | sp\|P08195\|4F2_HUMAN | -0,238765397 | 0,473490173 |
| 20388 | sp\|Q15370\|ELOB_HUMAN | 0,237771353 | 0,473846964 |
| 20206 | sp\|P39019\|RS19_HUMAN | 0,18079449 | 0,476248855 |
| 19300 | sp\|P00558\|PGK1_HUMAN | 0,163949882 | 0,476560652 |
| 19896 | sp\|Q9UQ80\|PA2G4_HUMAN | 0,177883838 | 0,478134433 |
| 20350 | sp\|P62258\|1433E_HUMAN | 0,206828365 | 0,47848502 |
| 20408 | sp\|P11387\|TOP1_HUMAN | 0,321469356 | 0,480084091 |
| 20369 | sp\|Q06323\|PSME1_HUMAN | 0,592122532 | 0,480945491 |
| 19280 | sp\|Q96RW7\|HMCN1_HUMAN | -0,266127904 | 0,484125263 |
| 19652 | sp\|P26038\|MOES_HUMAN | 0,151997415 | 0,485178127 |
| 20286 | sp\|Q5T6V5\|QSPP_HUMAN | -0,602660378 | 0,485542595 |
| 19378 | sp\|P21399\|ACOC_HUMAN | 0,183002914 | 0,48608378 |
| 19820 | sp\|P09382\|LEG1_HUMAN | -0,180993509 | 0,486514124 |
| 20113 | sp\|Q9BWS9\|CHID1_HUMAN | -0,330924456 | 0,487202944 |
| 19980 | sp\|Q00796\|DHSO_HUMAN | 0,192445611 | 0,48799794 |
| 20380 | sp\|P55809\|SCOT1_HUMAN | 0,249386605 | 0,488119933 |
| 19682 | sp\|Q13310\|PABP4_HUMAN | -0,319181473 | 0,489394183 |
| 19905 | sp\|Q16851\|UGPA_HUMAN | 0,161313014 | 0,49052627 |
| 19691 | sp\|Q6YHK3\|CD109_HUMAN | -0,164612187 | 0,493916897 |
| 19544 | sp\|O15347\|HMGB3_HUMAN | 0,543273937 | 0,494747513 |
| 20130 | sp\|O60462\|NRP2_HUMAN | -0,263154555 | 0,497857037 |
| 20228 | sp\|Q99497\|PARK7_HUMAN | 0,237555049 | 0,498286035 |
| 20084 | sp\|P27695\|APEX1_HUMAN | 0,209787966 | 0,499741699 |
| 19815 | sp\|P08603\|CFAH_HUMAN | -0,482359523 | 0,49988689 |
| 19978 | sp\|P01034\|CYTC_HUMAN | -0,306370319 | 0,501244768 |
| 20413 | sp\|Q86TI2\|DPP9_HUMAN | 0,178015696 | 0,501494417 |
| 19324 | sp\|P0DMV9\|HS71B_HUMAN | 0,139333215 | 0,50156207 |
| 19294 | sp\|P61981\|1433G_HUMAN | 0,183454231 | 0,502066909 |
| 19557 | sp\|P01023\|A2MG_HUMAN | -0,434360015 | 0,502604552 |
| 20336 | sp\|Q08945\|SSRP1_HUMAN | -0,577202299 | 0,503375534 |
| 19842 | sp\|P62328\|TYB4_HUMAN | 0,196059528 | 0,503771303 |
| 19373 | sp\|P18124\|RL7_HUMAN | 0,342596233 | 0,504079847 |
| 19435 | sp\|P46940\|IQGA1_HUMAN | 0,157342699 | 0,507356214 |
| 20075 | sp\|Q9UBT2\|SAE2_HUMAN | 0,24559222 | 0,508786447 |
| 19912 | sp\|Q13616\|CUL1_HUMAN | 0,209756986 | 0,509394546 |
| 20007 | sp\|P52565\|GDIR1_HUMAN | 0,18371615 | 0,509740269 |
| 19283 | sp\|P33176\|KINH_HUMAN | 0,319985678 | 0,510366291 |
| 20175 | sp\|P49189\|AL9A1_HUMAN | 0,189034833 | 0,511582692 |
| 19670 | sp\|P11216\|PYGB_HUMAN | 0,141915713 | 0,514712724 |
| 19604 | sp\|P22090\|RS4Y1_HUMAN | -0,338194441 | 0,515781732 |
| 19402 | sp\|P62906\|RL10A_HUMAN | 0,152660603 | 0,516743745 |
| 19399 | sp\|Q9UJJ9\|GNPTG_HUMAN | -0,329491428 | 0,517603823 |
| 19783 | sp\|P30086\|PEBP1_HUMAN | 0,170392315 | 0,517915922 |
| 19466 | sp\|P07195\|LDHB_HUMAN | 0,171751445 | 0,517988407 |
| 20245 | sp\|P18669\|PGAM1_HUMAN | 0,222967393 | 0,51841158 |

| | | | |
|---|---|---|---|
| 20461 | sp\|Q9NR30\|DDX21_HUMAN | 0,382889574 | 0,519696578 |
| 20210 | sp\|P46781\|RS9_HUMAN | 0,277552729 | 0,520103277 |
| 20410 | sp\|Q9NY33\|DPP3_HUMAN | 0,168934318 | 0,520373784 |
| 19500 | sp\|P02649\|APOE_HUMAN | -0,734480883 | 0,521143537 |
| 20244 | sp\|P98160\|PGBM_HUMAN | -0,180923718 | 0,521673126 |
| 19738 | sp\|P28482\|MK01_HUMAN | -0,214650345 | 0,522094165 |
| 19481 | sp\|B2RXH8\|HNRC2_HUMAN | -0,212550878 | 0,522562729 |
| 20257 | sp\|Q9ULV4\|COR1C_HUMAN | 0,154883944 | 0,522849179 |
| 19897 | sp\|P40925\|MDHC_HUMAN | 0,149116541 | 0,523072926 |
| 19892 | sp\|P61160\|ARP2_HUMAN | -0,168301123 | 0,52511151 |
| 19336 | sp\|P60981\|DEST_HUMAN | 0,203706131 | 0,527455181 |
| 20351 | sp\|Q13263\|TIF1B_HUMAN | 0,467808731 | 0,528122882 |
| 19992 | sp\|P05455\|LA_HUMAN | 0,209267371 | 0,529213479 |
| 20081 | sp\|P23381\|SYWC_HUMAN | 0,212306548 | 0,530367557 |
| 19653 | sp\|P15311\|EZRI_HUMAN | 0,136059634 | 0,532003336 |
| 19813 | sp\|P09132\|SRP19_HUMAN | 0,141439967 | 0,532495867 |
| 20474 | sp\|P11047\|LAMC1_HUMAN | -0,317307702 | 0,535945466 |
| 19709 | sp\|O75368\|SH3L1_HUMAN | 0,36068671 | 0,537664579 |
| 19600 | sp\|P01024\|CO3_HUMAN | -0,17849424 | 0,537860512 |
| 19995 | sp\|O75144\|ICOSL_HUMAN | 0,15991439 | 0,539595462 |
| 20242 | sp\|P07942\|LAMB1_HUMAN | -0,314910272 | 0,540481986 |
| 19701 | sp\|P09486\|SPRC_HUMAN | -0,219546696 | 0,540861586 |
| 20038 | sp\|O00468\|AGRIN_HUMAN | -0,446476219 | 0,541314046 |
| 19538 | sp\|Q9NPH3\|IL1AP_HUMAN | -0,168907538 | 0,542387428 |
| 20189 | sp\|P16035\|TIMP2_HUMAN | -0,236800388 | 0,543096588 |
| 19605 | sp\|Q96KP4\|CNDP2_HUMAN | 0,187460221 | 0,544465143 |
| 20470 | sp\|O75326\|SEM7A_HUMAN | -0,266510695 | 0,546456684 |
| 20333 | sp\|Q9NY27\|PP4R2_HUMAN | -0,294851409 | 0,546545739 |
| 20201 | sp\|Q9UKM7\|MA1B1_HUMAN | -0,233099777 | 0,549109317 |
| 19675 | sp\|P09104\|ENOG_HUMAN | 0,347334921 | 0,549217336 |
| 19993 | sp\|P08582\|TRFM_HUMAN | -0,397939401 | 0,549310695 |
| 19674 | sp\|Q9HB71\|CYBP_HUMAN | 0,201697063 | 0,550211042 |
| 20160 | sp\|Q13867\|BLMH_HUMAN | 0,179122795 | 0,551407063 |
| 19616 | sp\|Q9Y4L1\|HYOU1_HUMAN | -0,182248226 | 0,551444165 |
| 19516 | sp\|P51114\|FXR1_HUMAN | 0,272855141 | 0,552729722 |
| 19602 | sp\|O00154\|BACH_HUMAN | 0,159813538 | 0,553232874 |
| 20227 | sp\|P47914\|RL29_HUMAN | 0,155021015 | 0,553438144 |
| 20283 | sp\|P54136\|SYRC_HUMAN | 0,329609773 | 0,558566661 |
| 19788 | sp\|Q9P265\|DIP2B_HUMAN | -0,339152102 | 0,558716494 |
| 19681 | sp\|P11940\|PABP1_HUMAN | 0,214636326 | 0,559184244 |
| 19405 | sp\|P26583\|HMGB2_HUMAN | 0,344750148 | 0,561377409 |
| 20047 | sp\|Q9ULC4\|MCTS1_HUMAN | -0,255357207 | 0,561696949 |
| 19380 | sp\|P31153\|METK2_HUMAN | 0,368752682 | 0,561845217 |
| 19492 | sp\|P15531\|NDKA_HUMAN | 0,238080838 | 0,562053264 |
| 19703 | sp\|P60174\|TPIS_HUMAN | 0,184856431 | 0,562102982 |
| 19313 | sp\|P17050\|NAGAB_HUMAN | -0,316343554 | 0,562627623 |
| 19707 | sp\|Q9BR76\|COR1B_HUMAN | 0,230639721 | 0,564942471 |
| 19442 | sp\|P03950\|ANGI_HUMAN | -0,348151987 | 0,565245522 |
| 20091 | sp\|P08708\|RS17_HUMAN | 0,143882164 | 0,565819757 |
| 19629 | sp\|P62851\|RS25_HUMAN | 0,142041478 | 0,569041874 |
| 19565 | sp\|P09960\|LKHA4_HUMAN | 0,135520839 | 0,569281378 |
| 20002 | sp\|P35268\|RL22_HUMAN | 0,180128631 | 0,56939491 |
| 19736 | sp\|Q9UHD8\|SEPT9_HUMAN | 0,27342256 | 0,569961914 |
| 19486 | sp\|Q92859\|NEO1_HUMAN | 0,154588873 | 0,571578853 |
| 20033 | sp\|O43583\|DENR_HUMAN | 0,292703721 | 0,571601558 |
| 19547 | sp\|Q96C86\|DCPS_HUMAN | -0,270107391 | 0,572269448 |
| 19464 | sp\|P00338\|LDHA_HUMAN | 0,142498024 | 0,574318659 |
| 20041 | sp\|P58215\|LOXL3_HUMAN | 0,226165039 | 0,574556512 |
| 20166 | sp\|Q0VDG4\|SCRN3_HUMAN | 0,145085939 | 0,575414382 |
| 19319 | sp\|Q96AG4\|LRC59_HUMAN | 0,266351306 | 0,575819055 |
| 19628 | sp\|P04179\|SODM_HUMAN | 0,219176208 | 0,576791469 |
| 20098 | sp\|Q9UHY7\|ENOPH_HUMAN | 0,140860947 | 0,576909516 |
| 20432 | sp\|O00567\|NOP56_HUMAN | -0,178082022 | 0,577501501 |
| 19866 | sp\|Q01459\|DIAC_HUMAN | -0,41727652 | 0,578587753 |
| 19775 | sp\|P29966\|MARCS_HUMAN | 0,19202726 | 0,581536375 |
| 20022 | sp\|Q969E4\|TCAL3_HUMAN | 0,275031451 | 0,581861044 |
| 19846 | sp\|P62266\|RS23_HUMAN | 0,171618173 | 0,582041935 |
| 19951 | sp\|Q9UNF0\|PACN2_HUMAN | 0,128067554 | 0,583884651 |
| 20174 | sp\|Q14CX7\|NAA25_HUMAN | 0,192725138 | 0,583922271 |
| 19933 | sp\|P43487\|RANG_HUMAN | 0,186726855 | 0,584291443 |
| 19863 | sp\|P02545\|LMNA_HUMAN | 0,140688742 | 0,584330992 |
| 19455 | sp\|Q9NQW7\|XPP1_HUMAN | 0,161860899 | 0,585142069 |
| 19514 | sp\|P25398\|RS12_HUMAN | 0,135861138 | 0,585453525 |
| 20262 | sp\|Q8WWM7\|ATX2L_HUMAN | 0,196931236 | 0,585790703 |
| 20157 | sp\|P13667\|PDIA4_HUMAN | 0,140381658 | 0,586851281 |
| 20110 | sp\|P07237\|PDIA1_HUMAN | 0,15006307 | 0,587051809 |
| 19732 | sp\|P26012\|ITB8_HUMAN | -0,321006405 | 0,588151523 |
| 19779 | sp\|P69849\|NOMO3_HUMAN | -0,12549977 | 0,589413493 |

| | | | |
|---|---|---|---|
| 20138 | sp\|Q13822\|ENPP2_HUMAN | -0,179545845 | 0,590014689 |
| 20083 | sp\|Q86UD1\|OAF_HUMAN | -0,276938741 | 0,590235753 |
| 19805 | sp\|O15372\|EIF3H_HUMAN | 0,3514113 | 0,590280123 |
| 20059 | sp\|P12270\|TPR_HUMAN | 0,322522145 | 0,591144258 |
| 20287 | sp\|P25391\|LAMA1_HUMAN | -0,240930278 | 0,59138679 |
| 20015 | sp\|P62244\|RS15A_HUMAN | 0,177276122 | 0,592083386 |
| 20259 | sp\|Q9BWD1\|THIC_HUMAN | 0,155474922 | 0,592539705 |
| 20222 | sp\|P11362\|FGFR1_HUMAN | -0,174244916 | 0,592566399 |
| 19318 | sp\|P37108\|SRP14_HUMAN | 0,168766329 | 0,593601228 |
| 19572 | sp\|Q969P0\|IGSF8_HUMAN | -0,564340086 | 0,593841786 |
| 20154 | sp\|P12956\|XRCC6_HUMAN | 0,199332575 | 0,594585057 |
| 20060 | sp\|P41236\|IPP2_HUMAN | 0,209911973 | 0,595145194 |
| 19482 | sp\|P32119\|PRDX2_HUMAN | -0,260495497 | 0,595304528 |
| 20247 | sp\|O43493\|TGON2_HUMAN | -0,131080818 | 0,595815407 |
| 19869 | sp\|Q15424\|SAFB1_HUMAN | 0,265249226 | 0,596449385 |
| 19945 | sp\|Q14061\|COX17_HUMAN | 0,152310963 | 0,5966061 |
| 20004 | sp\|P29218\|IMPA1_HUMAN | -0,210134502 | 0,596960698 |
| 19292 | sp\|P63104\|1433Z_HUMAN | 0,132496885 | 0,597439201 |
| 19519 | sp\|P35052\|GPC1_HUMAN | -0,142366913 | 0,604572432 |
| 19493 | sp\|Q02818\|NUCB1_HUMAN | -0,326785684 | 0,604826334 |
| 19491 | sp\|P22392\|NDKB_HUMAN | 0,196301518 | 0,60486843 |
| 19608 | sp\|Q14914\|PTGR1_HUMAN | 0,210967299 | 0,605499629 |
| 19644 | sp\|P50990\|TCPQ_HUMAN | 0,144141339 | 0,607086526 |
| 19520 | sp\|P16152\|CBR1_HUMAN | 0,115231591 | 0,607258777 |
| 19333 | sp\|Q99832\|TCPH_HUMAN | 0,164067658 | 0,609086971 |
| 20014 | sp\|P61956\|SUMO2_HUMAN | 0,230671576 | 0,609181105 |
| 20062 | sp\|P62993\|GRB2_HUMAN | -0,201739843 | 0,610478157 |
| 19390 | sp\|P62917\|RL8_HUMAN | 0,160135142 | 0,612442066 |
| 19877 | sp\|P36955\|PEDF_HUMAN | -0,300877223 | 0,613379928 |
| 19778 | sp\|Q08257\|QOR_HUMAN | 0,100968539 | 0,615546451 |
| 20124 | sp\|Q04760\|LGUL_HUMAN | 0,167904401 | 0,615819782 |
| 19986 | sp\|P06744\|G6PI_HUMAN | 0,16878753 | 0,616208461 |
| 20479 | sp\|Q12904\|AIMP1_HUMAN | 0,133200327 | 0,619356385 |
| 19710 | sp\|Q9H3G5\|CPVL_HUMAN | -0,5982555 | 0,620177621 |
| 19969 | sp\|P13497\|BMP1_HUMAN | -0,136817848 | 0,621088742 |
| 19615 | sp\|P30405\|PPIF_HUMAN | 0,338227464 | 0,626337499 |
| 20465 | sp\|Q8IUX7\|AEBP1_HUMAN | -0,327723956 | 0,627275379 |
| 19438 | sp\|P07225\|PROS_HUMAN | -0,206684625 | 0,628324214 |
| 20365 | sp\|Q02878\|RL6_HUMAN | 0,122968365 | 0,630452816 |
| 19903 | sp\|Q10471\|GALT2_HUMAN | -0,156324349 | 0,63054904 |
| 20330 | sp\|P13489\|RINI_HUMAN | -0,22939552 | 0,631204444 |
| 19282 | sp\|P00492\|HPRT_HUMAN | 0,119710876 | 0,632838154 |
| 19553 | sp\|Q01581\|HMCS1_HUMAN | 0,205751381 | 0,633637203 |
| 20282 | sp\|P48745\|CCN3_HUMAN | -0,136292589 | 0,633936994 |
| 20294 | sp\|Q9Y2W1\|TR150_HUMAN | -0,270098424 | 0,634466344 |
| 20376 | sp\|Q15393\|SF3B3_HUMAN | 0,411047884 | 0,636341711 |
| 19321 | sp\|P43251\|BTD_HUMAN | -0,215831553 | 0,636991328 |
| 20453 | sp\|O94985\|CSTN1_HUMAN | -0,172751251 | 0,638307974 |
| 19907 | sp\|P37837\|TALDO_HUMAN | 0,192072575 | 0,639546252 |
| 19743 | sp\|P27797\|CALR_HUMAN | 0,105682377 | 0,640629979 |
| 19982 | sp\|O75223\|GGCT_HUMAN | 0,106125451 | 0,64218029 |
| 19640 | sp\|Q9UNZ2\|NSF1C_HUMAN | 0,242406397 | 0,64252958 |
| 20488 | sp\|P62318\|SMD3_HUMAN | -0,231437157 | 0,643257575 |
| 20056 | sp\|O60832\|DKC1_HUMAN | 0,460948094 | 0,644483349 |
| 20280 | sp\|P78417\|GSTO1_HUMAN | 0,119738666 | 0,645701731 |
| 19433 | sp\|P08134\|RHOC_HUMAN | -0,120156766 | 0,645949651 |
| 19635 | sp\|Q8N1G4\|LRC47_HUMAN | -0,152104535 | 0,646860471 |
| 19975 | sp\|Q14764\|MVP_HUMAN | 0,276822125 | 0,647228992 |
| 19946 | sp\|Q16836\|HCDH_HUMAN | 0,149838015 | 0,649984974 |
| 19375 | sp\|Q9Y2V2\|CHSP1_HUMAN | 0,213846045 | 0,650457488 |
| 19790 | sp\|P01033\|TIMP1_HUMAN | -0,253418939 | 0,650775611 |
| 19849 | sp\|P40429\|RL13A_HUMAN | 0,1256786 | 0,653108579 |
| 19925 | sp\|P49589\|SYCC_HUMAN | 0,165033567 | 0,653723059 |
| 19531 | sp\|P30101\|PDIA3_HUMAN | 0,115294636 | 0,65384479 |
| 20424 | sp\|P62316\|SMD2_HUMAN | -0,154510601 | 0,654309483 |
| 20211 | sp\|O95433\|AHSA1_HUMAN | 0,099810335 | 0,655801077 |
| 19965 | sp\|Q9Y266\|NUDC_HUMAN | 0,106839153 | 0,657563476 |
| 20256 | sp\|P48444\|COPD_HUMAN | -0,268044783 | 0,658413657 |
| 20322 | sp\|Q16666\|IF16_HUMAN | 0,13053873 | 0,65985706 |
| 19286 | sp\|P05120\|PAI2_HUMAN | 0,325779346 | 0,660159843 |
| 19424 | sp\|Q15149\|PLEC_HUMAN | 0,346595498 | 0,662573251 |
| 20082 | sp\|P53999\|TCP4_HUMAN | 0,368996411 | 0,663733742 |
| 20011 | sp\|O75937\|DNJC8_HUMAN | 0,204031015 | 0,66549075 |
| 19902 | sp\|Q15185\|TEBP_HUMAN | 0,174643467 | 0,667341333 |
| 19802 | sp\|P58546\|MTPN_HUMAN | -0,096891142 | 0,667717247 |
| 20415 | sp\|P43490\|NAMPT_HUMAN | 0,107196904 | 0,667810205 |
| 20450 | sp\|O95861\|BPNT1_HUMAN | 0,089674238 | 0,66880144 |
| 20221 | sp\|Q99523\|SORT_HUMAN | -0,165686431 | 0,669032945 |

| | | | |
|---|---|---|---|
| 20434 | sp\|P22234\|PUR6_HUMAN | 0,215815756 | 0,670302048 |
| 19505 | sp\|O14602\|IF1AY_HUMAN | 0,14367384 | 0,671010261 |
| 20181 | sp\|O15145\|ARPC3_HUMAN | 0,089283121 | 0,673898339 |
| 19741 | sp\|P27824\|CALX_HUMAN | -0,105063017 | 0,674041237 |
| 19698 | sp\|P0C0S5\|H2AZ_HUMAN | 0,20951596 | 0,674154608 |
| 20383 | sp\|Q9UN86\|G3BP2_HUMAN | 0,143652168 | 0,675443517 |
| 19512 | sp\|P51858\|HDGF_HUMAN | 0,129643974 | 0,676360206 |
| 20085 | sp\|P19823\|ITIH2_HUMAN | -0,420496661 | 0,678879787 |
| 20077 | sp\|P39748\|FEN1_HUMAN | 0,162785122 | 0,679074857 |
| 20118 | sp\|P13500\|CCL2_HUMAN | 0,1477292 | 0,679922792 |
| 20266 | sp\|P11233\|RALA_HUMAN | -0,145460042 | 0,680598591 |
| 20104 | sp\|O75116\|ROCK2_HUMAN | -0,335348214 | 0,68253711 |
| 19312 | sp\|P50579\|MAP2_HUMAN | 0,258001558 | 0,682829242 |
| 20195 | sp\|P17987\|TCPA_HUMAN | -0,117603606 | 0,683050854 |
| 19473 | sp\|P52209\|6PGD_HUMAN | 0,101683026 | 0,683878053 |
| 20034 | sp\|Q16881\|TRXR1_HUMAN | 0,160299003 | 0,684977597 |
| 19959 | sp\|P46821\|MAP1B_HUMAN | 0,249794321 | 0,68650658 |
| 19864 | sp\|Q14847\|LASP1_HUMAN | 0,167797006 | 0,688682612 |
| 19429 | sp\|P39023\|RL3_HUMAN | 0,11767073 | 0,688814742 |
| 20055 | sp\|Q8NBS9\|TXND5_HUMAN | 0,123814631 | 0,689836977 |
| 19521 | sp\|Q8NE71\|ABCF1_HUMAN | 0,195710074 | 0,690016201 |
| 20165 | sp\|P27816\|MAP4_HUMAN | -0,160201342 | 0,691139604 |
| 20402 | sp\|Q92520\|FAM3C_HUMAN | -0,174839643 | 0,691195524 |
| 19499 | sp\|Q8NC51\|PAIRB_HUMAN | -0,149007627 | 0,693114632 |
| 20323 | sp\|Q9NYU2\|UGGG1_HUMAN | -0,243893705 | 0,693605979 |
| 19427 | sp\|O43399\|TPD54_HUMAN | 0,332412686 | 0,694318843 |
| 20252 | sp\|Q16543\|CDC37_HUMAN | 0,205666207 | 0,694366619 |
| 19654 | sp\|P35241\|RADI_HUMAN | 0,104170148 | 0,694802455 |
| 19745 | sp\|P48643\|TCPE_HUMAN | 0,141644149 | 0,695518745 |
| 19386 | sp\|Q9NY97\|B3GN2_HUMAN | -0,126372335 | 0,699594329 |
| 19534 | sp\|P32455\|GBP1_HUMAN | 0,249771177 | 0,700577789 |
| 20354 | sp\|P09661\|RU2A_HUMAN | 0,220900963 | 0,70088412 |
| 19621 | sp\|P14314\|GLU2B_HUMAN | 0,101462566 | 0,701025357 |
| 19458 | sp\|P07108\|ACBP_HUMAN | 0,116990749 | 0,703860135 |
| 20187 | sp\|P61254\|RL26_HUMAN | -0,119834359 | 0,706209729 |
| 20472 | sp\|Q9Y6E2\|BZW2_HUMAN | 0,216033575 | 0,707780815 |
| 20274 | sp\|P04792\|HSPB1_HUMAN | 0,167235014 | 0,7086699 |
| 19843 | sp\|P61313\|RL15_HUMAN | 0,099486249 | 0,708700826 |
| 20231 | sp\|P07814\|SYEP_HUMAN | 0,127915832 | 0,709092903 |
| 19789 | sp\|P09417\|DHPR_HUMAN | 0,088593865 | 0,710230383 |
| 19724 | sp\|Q9H354\|YJ001_HUMAN | -0,077838235 | 0,7109772 |
| 19397 | sp\|Q9NPF2\|CHSTB_HUMAN | -0,307458878 | 0,712786121 |
| 20442 | sp\|P35520\|CBS_HUMAN | 0,104167487 | 0,712955237 |
| 19484 | sp\|Q99470\|SDF2_HUMAN | 0,20763224 | 0,713428694 |
| 19411 | sp\|P67936\|TPM4_HUMAN | 0,219165095 | 0,715146121 |
| 19664 | sp\|Q12906\|ILF3_HUMAN | 0,205533543 | 0,715239472 |
| 19495 | sp\|P50502\|F10A1_HUMAN | -0,111649799 | 0,715275291 |
| 19288 | sp\|P02795\|MT2_HUMAN | 0,176895767 | 0,716033253 |
| 20069 | sp\|Q15046\|SYK_HUMAN | 0,125083868 | 0,716971094 |
| 19441 | sp\|P07737\|PROF1_HUMAN | 0,107387008 | 0,717752688 |
| 19824 | sp\|Q15274\|NADC_HUMAN | 0,113761273 | 0,718800231 |
| 20128 | sp\|Q04637\|IF4G1_HUMAN | 0,136052186 | 0,720852992 |
| 20344 | sp\|P15880\|RS2_HUMAN | 0,109670711 | 0,722172936 |
| 20197 | sp\|P43121\|MUC18_HUMAN | -0,162248423 | 0,722352222 |
| 20391 | sp\|Q86UP2\|KTN1_HUMAN | 0,092627527 | 0,722410213 |
| 20131 | sp\|P09543\|CN37_HUMAN | -0,204425478 | 0,72244989 |
| 19751 | sp\|P07093\|GDN_HUMAN | -0,086406793 | 0,722866303 |
| 19388 | sp\|P06899\|H2B1J_HUMAN | 0,116332642 | 0,723313467 |
| 20299 | sp\|Q02543\|RL18A_HUMAN | 0,108478044 | 0,724394447 |
| 19342 | sp\|Q14517\|FAT1_HUMAN | -0,163911173 | 0,724754455 |
| 20484 | sp\|P49354\|FNTA_HUMAN | 0,097930873 | 0,727196647 |
| 20489 | sp\|O75882\|ATRN_HUMAN | -0,125170416 | 0,727755278 |
| 19795 | sp\|P08253\|MMP2_HUMAN | 0,101121591 | 0,727920425 |
| 19747 | sp\|Q9NRX4\|PHP14_HUMAN | 0,176972551 | 0,728883984 |
| 20338 | sp\|P04181\|OAT_HUMAN | -0,11502061 | 0,730436216 |
| 20378 | sp\|P23526\|SAHH_HUMAN | 0,080800095 | 0,730933558 |
| 20316 | sp\|Q96P70\|IPO9_HUMAN | 0,069553568 | 0,731547132 |
| 20319 | sp\|O00244\|ATOX1_HUMAN | 0,08814097 | 0,73291285 |
| 20430 | sp\|P01036\|CYTS_HUMAN | -0,338286982 | 0,733115058 |
| 20441 | sp\|Q9NSD9\|SYFB_HUMAN | 0,228415181 | 0,735713876 |
| 19392 | sp\|P38159\|RBMX_HUMAN | 0,152400288 | 0,736530508 |
| 20385 | sp\|Q9NQX1\|PRDM5_HUMAN | 0,099146121 | 0,737016592 |
| 19949 | sp\|P35659\|DEK_HUMAN | 0,216445796 | 0,738399416 |
| 20042 | sp\|P62280\|RS11_HUMAN | 0,118648061 | 0,739009569 |
| 20132 | sp\|P52788\|SPSY_HUMAN | -0,118107304 | 0,739284391 |
| 19511 | sp\|P60660\|MYL6_HUMAN | -0,107184178 | 0,740227229 |
| 19434 | sp\|P78539\|SRPX_HUMAN | -0,19132219 | 0,740881185 |
| 20218 | sp\|Q6UW49\|SPESP_HUMAN | 0,307087816 | 0,74107562 |

| | | | |
|---|---|---|---|
| 19767 | sp\|P31939\|PUR9_HUMAN | 0,099997485 | 0,741212331 |
| 20423 | sp\|Q13449\|LSAMP_HUMAN | -0,085082264 | 0,741401031 |
| 19476 | sp\|P43243\|MATR3_HUMAN | -0,18934623 | 0,743086544 |
| 19798 | sp\|Q4J6C6\|PPCEL_HUMAN | -0,243394713 | 0,745565666 |
| 19906 | sp\|P63165\|SUMO1_HUMAN | 0,296064083 | 0,746373157 |
| 19968 | sp\|P22455\|FGFR4_HUMAN | -0,140996351 | 0,747796427 |
| 19281 | sp\|P24534\|EF1B_HUMAN | -0,110266728 | 0,748298755 |
| 20078 | sp\|Q15084\|PDIA6_HUMAN | -0,099670861 | 0,749942872 |
| 19964 | sp\|P17405\|ASM_HUMAN | 0,123232582 | 0,750255882 |
| 19549 | sp\|P53004\|BIEA_HUMAN | -0,129018183 | 0,751130707 |
| 19867 | sp\|Q99536\|VAT1_HUMAN | 0,073694119 | 0,75118358 |
| 20185 | sp\|P05362\|ICAM1_HUMAN | 0,0906635 | 0,751201086 |
| 20285 | sp\|Q9BXK5\|B2L13_HUMAN | 0,110016621 | 0,75148441 |
| 20203 | sp\|P63167\|DYL1_HUMAN | 0,084692265 | 0,752094425 |
| 19838 | sp\|P35270\|SPRE_HUMAN | -0,137549382 | 0,753037572 |
| 19787 | sp\|P29692\|EF1D_HUMAN | -0,131260506 | 0,753729505 |
| 19963 | sp\|Q01082\|SPTB2_HUMAN | -0,141641177 | 0,753940105 |
| 20387 | sp\|Q9Y4K0\|LOXL2_HUMAN | -0,121837915 | 0,755444991 |
| 19914 | sp\|P61088\|UBE2N_HUMAN | 0,106725476 | 0,755810297 |
| 19614 | sp\|P50991\|TCPD_HUMAN | -0,070741336 | 0,760111631 |
| 19727 | sp\|P48147\|PPCE_HUMAN | 0,109813434 | 0,762270598 |
| 19702 | sp\|Q9Y383\|LC7L2_HUMAN | 0,160669416 | 0,762673448 |
| 19560 | sp\|Q9NZU5\|LMCD1_HUMAN | 0,167938621 | 0,763266441 |
| 20196 | sp\|Q01844\|EWS_HUMAN | -0,183465605 | 0,763279204 |
| 19352 | sp\|P54727\|RD23B_HUMAN | 0,082201726 | 0,765089601 |
| 19450 | sp\|Q03252\|LMNB2_HUMAN | -0,153821125 | 0,765324335 |
| 20151 | sp\|O95881\|TXD12_HUMAN | 0,196427369 | 0,768541578 |
| 19832 | sp\|P54819\|KAD2_HUMAN | -0,151554747 | 0,769321376 |
| 19850 | sp\|P35237\|SPB6_HUMAN | -0,09132654 | 0,769424076 |
| 20339 | sp\|P60033\|CD81_HUMAN | 0,294081224 | 0,769932373 |
| 20046 | sp\|P17096\|HMGA1_HUMAN | 0,29432657 | 0,771761782 |
| 19461 | sp\|P21291\|CSRP1_HUMAN | 0,075530272 | 0,773382466 |
| 19297 | sp\|O43768\|ENSA_HUMAN | -0,168251599 | 0,775448626 |
| 19768 | sp\|P15121\|ALDR_HUMAN | -0,057233587 | 0,777731206 |
| 19818 | sp\|P11413\|G6PD_HUMAN | 0,095776972 | 0,778181303 |
| 20403 | sp\|Q9Y315\|DEOC_HUMAN | 0,109823733 | 0,778202769 |
| 20102 | sp\|Q6ZMJ2\|SCAR5_HUMAN | 0,148576168 | 0,780405426 |
| 20065 | sp\|P68104\|EF1A1_HUMAN | -0,096057516 | 0,780456797 |
| 20341 | sp\|Q02952\|AKA12_HUMAN | -0,085860022 | 0,782853023 |
| 19958 | sp\|Q00839\|HNRPU_HUMAN | 0,127494815 | 0,782963438 |
| 20199 | sp\|O00391\|QSOX1_HUMAN | -0,171183087 | 0,783588245 |
| 19302 | sp\|P60709\|ACTB_HUMAN | 0,065272542 | 0,78428877 |
| 19546 | sp\|Q9ULB5\|CADH7_HUMAN | 0,110258625 | 0,784555285 |
| 20429 | sp\|O60869\|EDF1_HUMAN | 0,158820346 | 0,785703202 |
| 19997 | sp\|Q99075\|HBEGF_HUMAN | -0,082927591 | 0,787660802 |
| 20168 | sp\|P02751\|FINC_HUMAN | -0,09213635 | 0,788462581 |
| 20348 | sp\|P08579\|RU2B_HUMAN | 0,11900899 | 0,788510659 |
| 20292 | sp\|P55010\|IF5_HUMAN | 0,099578112 | 0,788894682 |
| 20121 | sp\|P63208\|SKP1_HUMAN | -0,07648748 | 0,788905776 |
| 19470 | sp\|P08238\|HS90B_HUMAN | 0,064652573 | 0,789096693 |
| 19731 | sp\|Q6UVY6\|MOXD1_HUMAN | -0,118217169 | 0,79480301 |
| 19613 | sp\|P23588\|IF4B_HUMAN | 0,07435878 | 0,800004112 |
| 19624 | sp\|Q16831\|UPP1_HUMAN | 0,061828166 | 0,80030385 |
| 19750 | sp\|P63220\|RS21_HUMAN | -0,122772362 | 0,800440595 |
| 20301 | sp\|P62913\|RL11_HUMAN | 0,089534278 | 0,800485254 |
| 20398 | sp\|Q9Y5B9\|SP16H_HUMAN | 0,146780771 | 0,80048659 |
| 19586 | sp\|Q9NUQ9\|CYRIB_HUMAN | 0,092942368 | 0,801458279 |
| 19671 | sp\|P06737\|PYGL_HUMAN | -0,058652386 | 0,803540669 |
| 20076 | sp\|Q05682\|CALD1_HUMAN | 0,085915763 | 0,805059264 |
| 20120 | sp\|O75874\|IDHC_HUMAN | 0,060947873 | 0,805954098 |
| 20237 | sp\|O00299\|CLIC1_HUMAN | 0,060762795 | 0,807084157 |
| 20025 | sp\|P51572\|BAP31_HUMAN | 0,116715606 | 0,807169551 |
| 20089 | sp\|Q7KZF4\|SND1_HUMAN | 0,089600457 | 0,807513315 |
| 19688 | sp\|Q9BRK5\|CAB45_HUMAN | -0,091161051 | 0,807949728 |
| 19506 | sp\|Q92823\|NRCAM_HUMAN | 0,10209871 | 0,811091094 |
| 19981 | sp\|P46063\|RECQ1_HUMAN | -0,132148235 | 0,814391314 |
| 20335 | sp\|Q96FW1\|OTUB1_HUMAN | -0,152100964 | 0,814990236 |
| 19335 | sp\|P23528\|COF1_HUMAN | 0,062667222 | 0,816144901 |
| 19733 | sp\|P62854\|RS26_HUMAN | 0,090220572 | 0,816217374 |
| 20248 | sp\|P62277\|RS13_HUMAN | 0,146160137 | 0,816645173 |
| 20420 | sp\|Q9UNW1\|MINP1_HUMAN | -0,103347959 | 0,816935235 |
| 20491 | sp\|P38919\|IF4A3_HUMAN | 0,067463465 | 0,817032775 |
| 19898 | sp\|O15067\|PUR4_HUMAN | 0,062589642 | 0,817496413 |
| 19941 | sp\|P19338\|NUCL_HUMAN | 0,085483318 | 0,81954043 |
| 19517 | sp\|Q01105\|SET_HUMAN | 0,069041698 | 0,820416303 |
| 20463 | sp\|Q13813\|SPTN1_HUMAN | 0,074940797 | 0,823125077 |
| 19940 | sp\|Q15121\|PEA15_HUMAN | -0,081156562 | 0,82320187 |
| 20115 | sp\|P61970\|NTF2_HUMAN | 0,075672806 | 0,824530206 |

| | | | |
|---|---|---|---|
| 19645 | sp\|O76021\|RL1D1_HUMAN | 0,056323256 | 0,826049433 |
| 20234 | sp\|Q86UE4\|LYRIC_HUMAN | 0,064706137 | 0,826818769 |
| 19343 | sp\|O95394\|AGM1_HUMAN | 0,097015575 | 0,829117622 |
| 19966 | sp\|P60842\|IF4A1_HUMAN | 0,058693303 | 0,829157954 |
| 19475 | sp\|P40121\|CAPG_HUMAN | 0,073617423 | 0,829591063 |
| 19786 | sp\|B5ME19\|EIFCL_HUMAN | 0,082096604 | 0,830439284 |
| 19819 | sp\|Q9BRA2\|TXD17_HUMAN | 0,07923581 | 0,830950848 |
| 19962 | sp\|Q13126\|MTAP_HUMAN | 0,050521365 | 0,831571449 |
| 19309 | sp\|P16401\|H15_HUMAN | 0,132814286 | 0,832098468 |
| 19315 | sp\|P16104\|H2AX_HUMAN | 0,16414675 | 0,833759649 |
| 20150 | sp\|O43847\|NRDC_HUMAN | 0,069727434 | 0,835635313 |
| 20043 | sp\|P14618\|KPYM_HUMAN | 0,044897988 | 0,835722497 |
| 20340 | sp\|P04406\|G3P_HUMAN | -0,09828121 | 0,838430375 |
| 19957 | sp\|Q9Y6N7\|ROBO1_HUMAN | 0,04369401 | 0,839066555 |
| 19804 | sp\|P31948\|STIP1_HUMAN | 0,078561448 | 0,84187179 |
| 19922 | sp\|P30085\|KCY_HUMAN | 0,045793396 | 0,842155221 |
| 20454 | sp\|Q13564\|ULA1_HUMAN | -0,047334123 | 0,84221938 |
| 19537 | sp\|O00339\|MATN2_HUMAN | 0,090846823 | 0,842670264 |
| 19637 | sp\|P11766\|ADHX_HUMAN | 0,06293872 | 0,843085812 |
| 19791 | sp\|Q9H2E6\|SEM6A_HUMAN | -0,10659633 | 0,844309722 |
| 20045 | sp\|P30041\|PRDX6_HUMAN | 0,063699152 | 0,847234559 |
| 20250 | sp\|Q9P2E9\|RRBP1_HUMAN | -0,094880669 | 0,848013796 |
| 20053 | sp\|P05114\|HMGN1_HUMAN | 0,081425537 | 0,848326176 |
| 19595 | sp\|P26641\|EF1G_HUMAN | 0,050415962 | 0,849021931 |
| 20281 | sp\|O00622\|CCN1_HUMAN | -0,062228097 | 0,849293689 |
| 19943 | sp\|P80303\|NUCB2_HUMAN | -0,099653463 | 0,851040006 |
| 20251 | sp\|Q12905\|ILF2_HUMAN | -0,084428181 | 0,851057479 |
| 20313 | sp\|Q9Y3F4\|STRAP_HUMAN | 0,090639323 | 0,851888534 |
| 19803 | sp\|Q15113\|PCOC1_HUMAN | -0,062975822 | 0,852473602 |
| 19871 | sp\|P49915\|GUAA_HUMAN | 0,095902368 | 0,853678405 |
| 19533 | sp\|P62805\|H4_HUMAN | 0,065729676 | 0,854660409 |
| 19700 | sp\|P26358\|DNMT1_HUMAN | 0,088066803 | 0,85527165 |
| 19777 | sp\|Q14738\|2A5D_HUMAN | -0,042983562 | 0,856522765 |
| 19780 | sp\|P10768\|ESTD_HUMAN | 0,04946613 | 0,859274517 |
| 20008 | sp\|Q9UHB6\|LIMA1_HUMAN | 0,053560359 | 0,862844514 |
| 19523 | sp\|Q13428\|TCOF_HUMAN | 0,111805515 | 0,863684459 |
| 20418 | sp\|Q16270\|IBP7_HUMAN | 0,057022216 | 0,8645794 |
| 20467 | sp\|P26373\|RL13_HUMAN | 0,05504476 | 0,866597646 |
| 19328 | sp\|P35579\|MYH9_HUMAN | -0,075245995 | 0,869479085 |
| 20421 | sp\|P61604\|CH10_HUMAN | -0,05853422 | 0,871259096 |
| 19603 | sp\|O75390\|CISY_HUMAN | -0,051841743 | 0,872066682 |
| 19771 | sp\|Q15459\|SF3A1_HUMAN | 0,092496022 | 0,872266816 |
| 19935 | sp\|P49588\|SYAC_HUMAN | 0,144204216 | 0,872493998 |
| 19936 | sp\|P49327\|FAS_HUMAN | 0,056480561 | 0,873292703 |
| 19284 | sp\|Q9NR45\|SIAS_HUMAN | 0,128967483 | 0,875823169 |
| 19759 | sp\|Q14204\|DYHC1_HUMAN | -0,071252354 | 0,875838367 |
| 19354 | sp\|P39687\|AN32A_HUMAN | 0,052908357 | 0,87616171 |
| 19469 | sp\|P07900\|HS90A_HUMAN | 0,035495179 | 0,876308484 |
| 19970 | sp\|P14866\|HNRPL_HUMAN | -0,049608554 | 0,876880633 |
| 20225 | sp\|P24821\|TENA_HUMAN | -0,046205478 | 0,880750119 |
| 19407 | sp\|O43707\|ACTN4_HUMAN | 0,039690948 | 0,884894012 |
| 20209 | sp\|P27694\|RFA1_HUMAN | 0,043280543 | 0,886415606 |
| 19734 | sp\|P53396\|ACLY_HUMAN | -0,038940722 | 0,887106807 |
| 20063 | sp\|Q13185\|CBX3_HUMAN | -0,04616719 | 0,888258396 |
| 19762 | sp\|P55145\|MANF_HUMAN | -0,066161793 | 0,888279686 |
| 19416 | sp\|P30040\|ERP29_HUMAN | 0,038173212 | 0,897681607 |
| 19999 | sp\|Q9BY32\|ITPA_HUMAN | -0,039972619 | 0,89861719 |
| 19938 | sp\|A6NN14\|ZN729_HUMAN | 0,046775968 | 0,898987672 |
| 20468 | sp\|P46778\|RL21_HUMAN | 0,03863363 | 0,899404383 |
| 19587 | sp\|P36578\|RL4_HUMAN | -0,035296226 | 0,899427496 |
| 19687 | sp\|P14550\|AK1A1_HUMAN | 0,0546711 | 0,899566894 |
| 19735 | sp\|O15394\|NCAM2_HUMAN | -0,052588359 | 0,900582755 |
| 20087 | sp\|O15305\|PMM2_HUMAN | 0,056963753 | 0,902617648 |
| 19891 | sp\|P62857\|RS28_HUMAN | 0,030583583 | 0,902940532 |
| 19837 | sp\|P40123\|CAP2_HUMAN | 0,044229832 | 0,906767314 |
| 20444 | sp\|P61086\|UBE2K_HUMAN | 0,03481839 | 0,907868908 |
| 20049 | sp\|Q15642\|CIP4_HUMAN | -0,023119385 | 0,909432258 |
| 20000 | sp\|P16949\|STMN1_HUMAN | -0,053485583 | 0,909530228 |
| 20134 | sp\|Q9UKX7\|NUP50_HUMAN | -0,05256768 | 0,910425504 |
| 20370 | sp\|Q14978\|NOLC1_HUMAN | -0,08112567 | 0,912638978 |
| 20146 | sp\|Q07955\|SRSF1_HUMAN | 0,036286887 | 0,913599615 |
| 19910 | sp\|Q9GZN8\|CT027_HUMAN | 0,074363453 | 0,915760564 |
| 19668 | sp\|P09874\|PARP1_HUMAN | 0,061816211 | 0,916949638 |
| 20213 | sp\|P35998\|PRS7_HUMAN | -0,070505134 | 0,9169601 |
| 19853 | sp\|P54577\|SYYC_HUMAN | 0,049067742 | 0,918442413 |
| 19692 | sp\|P55884\|EIF3B_HUMAN | 0,059729426 | 0,918849223 |
| 19924 | sp\|Q13907\|IDI1_HUMAN | -0,020869641 | 0,923049268 |
| 19502 | sp\|Q9Y490\|TLN1_HUMAN | 0,037523837 | 0,927674742 |

| | | | |
|---|---|---|---|
| 19683 | sp\|Q13435\|SF3B2_HUMAN | -0,034458268 | 0,929006005 |
| 19822 | sp\|Q13433\|S39A6_HUMAN | -0,02896282 | 0,929035038 |
| 19760 | sp\|Q15075\|EEA1_HUMAN | 0,034118595 | 0,93406648 |
| 19420 | sp\|P19022\|CADH2_HUMAN | -0,01765922 | 0,935047549 |
| 19776 | sp\|P49321\|NASP_HUMAN | 0,036523493 | 0,935863817 |
| 19382 | sp\|P13010\|XRCC5_HUMAN | 0,038042496 | 0,935895228 |
| 19364 | sp\|Q01469\|FABP5_HUMAN | 0,016197087 | 0,938243807 |
| 20208 | sp\|Q9P1F3\|ABRAL_HUMAN | -0,028238421 | 0,941171698 |
| 20440 | sp\|P62249\|RS16_HUMAN | -0,021802188 | 0,94170526 |
| 20327 | sp\|Q15942\|ZYX_HUMAN | 0,039183527 | 0,943125594 |
| 19793 | sp\|P31431\|SDC4_HUMAN | 0,019416297 | 0,943627496 |
| 20437 | sp\|P63244\|RACK1_HUMAN | -0,023613326 | 0,945374722 |
| 19657 | sp\|P22314\|UBA1_HUMAN | -0,036905233 | 0,949065704 |
| 19870 | sp\|O60841\|IF2P_HUMAN | -0,044881967 | 0,949085454 |
| 20296 | sp\|Q9UKK9\|NUDT5_HUMAN | -0,019842334 | 0,949282746 |
| 19875 | sp\|Q9ULF5\|S39AA_HUMAN | 0,038555525 | 0,949495095 |
| 20384 | sp\|P23246\|SFPQ_HUMAN | -0,023950568 | 0,949664546 |
| 19344 | sp\|P68363\|TBA1B_HUMAN | -0,040904838 | 0,950417975 |
| 19697 | sp\|P43686\|PRS6B_HUMAN | 0,01922973 | 0,951071783 |
| 19555 | sp\|Q14247\|SRC8_HUMAN | 0,035264659 | 0,951100299 |
| 20307 | sp\|O75369\|FLNB_HUMAN | -0,03526509 | 0,951659283 |
| 19563 | sp\|P60953\|CDC42_HUMAN | -0,016668211 | 0,951726003 |
| 19576 | sp\|P14625\|ENPL_HUMAN | -0,013749834 | 0,952527211 |
| 20147 | sp\|P15018\|LIF_HUMAN | -0,046762468 | 0,953720657 |
| 19799 | sp\|P04080\|CYTB_HUMAN | 0,013268802 | 0,956660355 |
| 19656 | sp\|P60866\|RS20_HUMAN | -0,017022637 | 0,956678202 |
| 20372 | sp\|O14974\|MYPT1_HUMAN | 0,017689732 | 0,95686358 |
| 20328 | sp\|P49368\|TCPG_HUMAN | 0,015139055 | 0,957684105 |
| 19412 | sp\|P63241\|IF5A1_HUMAN | 0,011054097 | 0,958030286 |
| 19686 | sp\|P78371\|TCPB_HUMAN | 0,011798644 | 0,958686771 |
| 19895 | sp\|Q14118\|DAG1_HUMAN | 0,018243209 | 0,960195504 |
| 20136 | sp\|Q9UBQ7\|GRHPR_HUMAN | -0,015033516 | 0,960701422 |
| 20275 | sp\|Q969H8\|MYDGF_HUMAN | 0,011113187 | 0,960811636 |
| 19414 | sp\|Q16181\|SEPT7_HUMAN | 0,023677126 | 0,961134788 |
| 20129 | sp\|P26368\|U2AF2_HUMAN | 0,037497777 | 0,961405056 |
| 19730 | sp\|Q9BXJ9\|NAA15_HUMAN | -0,022547903 | 0,961742113 |
| 19756 | sp\|P06748\|NPM_HUMAN | -0,012377665 | 0,962205914 |
| 19971 | sp\|O94973\|AP2A2_HUMAN | -0,017014444 | 0,962786897 |
| 20103 | sp\|Q5VY80\|ULBP6_HUMAN | -0,024799323 | 0,963240299 |
| 19806 | sp\|Q01813\|PFKAP_HUMAN | 0,020755565 | 0,963641505 |
| 20480 | sp\|O15540\|FABP7_HUMAN | 0,032131779 | 0,963979773 |
| 20106 | sp\|P14780\|MMP9_HUMAN | 0,016849947 | 0,965400857 |
| 20431 | sp\|Q9UKY7\|CDV3_HUMAN | -0,041667428 | 0,965666066 |
| 20093 | sp\|O75475\|PSIP1_HUMAN | -0,021024028 | 0,965795305 |
| 20241 | sp\|P48507\|GSH0_HUMAN | -0,013277274 | 0,966144216 |
| 20100 | sp\|P46060\|RAGP1_HUMAN | -0,019872713 | 0,966412342 |
| 19661 | sp\|Q99715\|COCA1_HUMAN | -0,02896038 | 0,971233644 |
| 19748 | sp\|P26639\|SYTC_HUMAN | -0,008788629 | 0,971849996 |
| 20009 | sp\|Q14697\|GANAB_HUMAN | -0,012755415 | 0,973388225 |
| 19539 | sp\|Q86VP6\|CAND1_HUMAN | 0,00943536 | 0,978315277 |
| 20265 | sp\|P11234\|RALB_HUMAN | 0,007441367 | 0,978983931 |
| 20044 | sp\|P42167\|LAP2B_HUMAN | -0,010249105 | 0,979736569 |
| 19723 | sp\|Q09028\|RBBP4_HUMAN | 0,013337261 | 0,980058823 |
| 19510 | sp\|Q15181\|IPYR_HUMAN | -0,010901045 | 0,9804951 |
| 19828 | sp\|P13591\|NCAM1_HUMAN | -0,004534065 | 0,980778008 |
| 20052 | sp\|Q9H299\|SH3L3_HUMAN | -0,008618776 | 0,981173778 |
| 19423 | sp\|P30044\|PRDX5_HUMAN | 0,014234992 | 0,983044383 |
| 19792 | sp\|Q99798\|ACON_HUMAN | 0,005137053 | 0,983232486 |
| 20143 | sp\|Q15293\|RCN1_HUMAN | -0,007632568 | 0,983800008 |
| 19334 | sp\|Q9Y281\|COF2_HUMAN | 0,006656281 | 0,983898302 |
| 20342 | sp\|P13612\|ITA4_HUMAN | 0,005431193 | 0,985083396 |
| 19720 | sp\|Q16555\|DPYL2_HUMAN | -0,009342097 | 0,985888705 |
| 19931 | sp\|P82979\|SARNP_HUMAN | 0,013168358 | 0,985980987 |
| 19848 | sp\|Q86SQ4\|AGRG6_HUMAN | 0,005663472 | 0,987380681 |
| 19532 | sp\|P40227\|TCPZ_HUMAN | -0,006910311 | 0,987824984 |
| 19401 | sp\|P13639\|EF2_HUMAN | -0,004254428 | 0,988684385 |
| 19836 | sp\|Q01518\|CAP1_HUMAN | -0,003123745 | 0,991824623 |
| 20010 | sp\|P18621\|RL17_HUMAN | 0,002482544 | 0,992406506 |
| 19431 | sp\|P09972\|ALDOC_HUMAN | 0,002532922 | 0,992622531 |
| 19293 | sp\|P27348\|1433T_HUMAN | -0,003093108 | 0,992898037 |
| 20293 | sp\|P83731\|RL24_HUMAN | -0,002603466 | 0,99457974 |
| 20473 | sp\|Q7L1Q6\|BZW1_HUMAN | 0,000935706 | 0,99852905 |
| 19356 | sp\|O60907\|TBL1X_HUMAN | 0,000561688 | 0,999082641 |
