## Supplementary material for "Peripheral positioning of lysosomes supports melanoma aggressiveness": Table 3

### Table 4: Human Protease Array (corresponding to Figure S4A)

| | Control | Control | Control | Control | Rapalog 5nM | Rapalog 5nM | Rapalog 5nM | Rapalog 5nM |
|---|---|---|---|---|---|---|---|---|
| ADAM8 | 1.147717 | 0.852283 | 1.15873 | 0.84127 | 0.774411 | 0.533206 | 0.746363 | 0.587302 |
| ADAM9 | 1.027607 | 0.972393 | 1.024194 | 0.975806 | 0.620266 | 0.592668 | 0.839214 | 0.808132 |
| ADAMTS1 | 1.023897 | 0.976103 | 0.992701 | 1.007299 | 0.556102 | 0.547179 | 0.792214 | 0.810219 |
| ADAMTS13 | 1.004161 | 0.995839 | 0.974874 | 1.025126 | 0.583189 | 0.587307 | 0.875419 | 0.89866 |
| Cathepsin A | 0.997188 | 1.002812 | 0.970847 | 1.029153 | 0.651642 | 0.689132 | 0.606976 | 0.61868 |
| Cathepsin B | 1.02308 | 0.97692 | 0.983442 | 1.016558 | 0.690258 | 0.642286 | 0.62038 | 0.617672 |
| Cathepsin C | 0.896558 | 1.103442 | 0.930233 | 1.069767 | 0.516407 | 0.625453 | 0.624862 | 0.665836 |
| Cathepsin D | 0.999935 | 1.000065 | 0.985497 | 1.014503 | 0.731973 | 0.623161 | 0.562947 | 0.60059 |
| Cathepsin E | 1.094293 | 0.905707 | 1.057143 | 0.942857 | 0.541856 | 0.47676 | 0.63869 | 0.697421 |
| Cathepsin L | 0.99064 | 1.00936 | 0.972477 | 1.027523 | 0.58572 | 0.590158 | 0.742546 | 0.912271 |
| Cathepsin S | 1.020089 | 0.979911 | 0.971026 | 1.028974 | 0.571797 | 0.596897 | 0.692362 | 0.7153 |
| Cathepsin V | 1.006443 | 0.993557 | 0.954325 | 1.045675 | 0.566174 | 0.584872 | 0.800049 | 0.791059 |
| Cathepsin X/Z/P | 0.966757 | 1.033243 | 0.971185 | 1.028815 | 0.701833 | 0.730565 | 0.644144 | 0.654838 |
| DPPIV/CD26 | 1.117334 | 0.882666 | 1.042017 | 0.957983 | 0.801751 | 0.659795 | 0.796744 | 0.757878 |
| Kallikrein 3/PSA | 0.985857 | 1.014143 | 1.054054 | 0.945946 | 0.551682 | 0.72544 | 0.833334 | 0.822917 |
| Kallikrein 5 | 0.974425 | 1.025575 | 1 | 1 | 0.670176 | 0.718398 | 0.639628 | 0.811835 |
| Kallikrein 6 | 0.942437 | 1.057563 | 0.965986 | 1.034014 | 0.706947 | 0.676587 | 0.89144 | 0.839002 |
| Kallikrein 7 | 0.961601 | 1.038399 | 1.008772 | 0.991228 | 0.497709 | 0.556055 | 0.872259 | 0.966923 |
| Kallikrein 10 | 0.961426 | 1.038574 | 1.019355 | 0.980645 | 0.600281 | 0.659802 | 0.974731 | 0.944893 |
| Kallikrein 11 | 1.056922 | 0.943078 | 1.034965 | 0.965035 | 0.59863 | 0.684269 | 0.776224 | 0.765443 |
| Kallikrein 13 | 0.943741 | 1.056259 | 0.944724 | 1.055276 | 0.641374 | 0.736734 | 0.774707 | 0.852178 |
| MMP-1 | 1.045864 | 0.954136 | 0.990291 | 1.009709 | 0.732064 | 0.699026 | 0.778317 | 0.785801 |
| MMP-2 | 0.977902 | 1.022098 | 0.967157 | 1.032843 | 0.696487 | 0.70715 | 0.700996 | 0.701869 |
| MMP-3 | 1.002066 | 0.997934 | 1.0033 | 0.9967 | 0.679249 | 0.815638 | 0.778465 | 0.717409 |
| MMP-7 | 0.981057 | 1.018943 | 1.001368 | 0.998632 | 0.549956 | 0.554512 | 0.795087 | 0.76767 |
| MMP-8 | 1.004194 | 0.995806 | 1.007722 | 0.992278 | 0.589924 | 0.604858 | 0.732143 | 0.761905 |
| MMP-9 | 0.987 | 1.013 | 1.017751 | 0.982249 | 0.648874 | 0.66088 | 0.723701 | 0.675049 |
| MMP-10 | 1.0573 | 0.9427 | 1.063291 | 0.936709 | 0.508022 | 0.449979 | 0.731804 | 0.868407 |
| MMP-12 | 0.956382 | 1.043618 | 0.93007 | 1.06993 | 0.603263 | 0.645063 | 0.927156 | 0.981061 |
| MMP-13 | 1.017502 | 0.982498 | 1 | 1 | 0.824547 | 0.658938 | 0.79974 | 0.790104 |
| Nepriltsin/CD10 | 1.020975 | 0.979025 | 0.938547 | 1.061453 | 0.638855 | 0.614965 | 0.861266 | 0.766527 |
| Presenilin | 1.031023 | 0.968977 | 1.035294 | 0.964706 | 0.695308 | 0.721402 | 0.825245 | 0.761765 |
| Proprotein Convertase 9 | 1.088068 | 0.911932 | 1.100324 | 0.899676 | 0.577882 | 0.473507 | 0.893069 | 0.708468 |
| Proteinase 3 | 1.044048 | 0.955952 | 0.907692 | 1.092308 | 0.433047 | 0.407147 | 0.830128 | 0.853846 |
| uPA/Urokinase | 1.002609 | 0.997391 | 1.078125 | 0.921875 | 0.582279 | 0.564111 | 0.794922 | 0.963542 |

| Control [%] | Rapalog [% of control] |
|---|---|
| 100 | 66.0 |
| 100 | 71.5 |
| 100 | 67.6 |
| 100 | 73.6 |
| 100 | 64.2 |
| 100 | 64.3 |
| 100 | 60.8 |
| 100 | 63.0 |
| 100 | 58.9 |
| 100 | 70.8 |
| 100 | 64.4 |
| 100 | 68.6 |
| 100 | 68.3 |
| 100 | 75.4 |
| 100 | 73.3 |
| 100 | 71.0 |
| 100 | 77.8 |
| 100 | 72.3 |
| 100 | 79.5 |
| 100 | 70.6 |
| 100 | 75.1 |
| 100 | 74.9 |
| 100 | 70.2 |
| 100 | 74.8 |
| 100 | 66.7 |
| 100 | 67.2 |
| 100 | 67.7 |
| 100 | 64.0 |
| 100 | 78.9 |
| 100 | 76.8 |
| 100 | 72.0 |
| 100 | 75.1 |
| 100 | 66.3 |
| 100 | 63.1 |
| 100 | 72.6 |

*Left* : Human protease array, levels of secretion in cells with spread and clustered lysosomes (control, 5nM Rapalog, respectively).

*Right* : Secretion levels are shown as percentage from the control.
